# Hierarchical substrates of prediction in visual cortical spiking

**DOI:** 10.1101/2024.10.02.616378

**Authors:** Jacob A. Westerberg, Yihan S. Xiong, Eli Sennesh, Hamed Nejat, David Ricci, Mazyar Azmi, Séverine Durand, Ben Hardcastle, Hannah Cabasco, Hannah Belski, Ahad Bawany, Ryan Gillis, Henry Loeffler, Carter R. Peene, Warren Han, Katrina Nguyen, Vivian Ha, Tye Johnson, Conor Grasso, Ahrial Young, Jackie Swapp, Ben Ouellette, Shiella Caldejon, Ali Williford, Peter A. Groblewski, Shawn R. Olsen, Carly Kiselycznyk, Christof Koch, Jerome A. Lecoq, Alexander Maier, André M. Bastos

## Abstract

Hierarchical predictive coding (HPC) models have recently flourished in neuroscience^1–9^. Feedforward and feedback processing are at the heart of HPC models. Previous experimental studies using fMRI, EEG/MEG, and LFP^9–11^ do not reliably resolve feedback modulation from local computations and feedforward outputs. Here, using open-science^8^, multi-species, multi-area, high-density^12^, laminar neurophysiology^13^, we empirically test whether hierarchical predictive coding is a key component shaping cortical processing of visual stimuli. To isolate visual information processing and eliminate motor/reward confounders^9–11^, we use a no-report task. Our task leveraged so-called global oddballs (GO) as unpredictable, deviant stimuli that circumvent low-level adaptation. We examined their responses relative to local oddballs (LO) that we habituated into highly predictable priors. Four surprising findings in this dataset challenge many existing hierarchical predictive coding models. First, GO responses were exclusive to higher-order, more cognitive areas rather than early-to-mid visual cortex. Second, inhibitory interneuron-targeted optogenetics in primates and mice and waveform shape analysis in primates revealed no evidence that predictive suppression was implemented via these interneurons. Third, highly predictable LO responses dominated in over 50% of all neurons, including in higher-order cortex, which should have anticipated them, indicating limited evidence for predictive suppression. Lastly, prediction error responses evoked by GOs did not evoke feedforward processing. These results reveal circuit dynamics that govern how prediction shapes visual processing, motivating more neurally constrained predictive processing models.

## Introduction

Predictive Processing (PP) models state that brains have evolved to model and create predictions about the sensory statistical regularities in the world^1,14,15^. In one prominent example of this, Hierarchical Predictive Coding (HPC), deviations from these internal predictions are neuronally signaled with prediction error responses^1–3^. Predictable stimuli are uninformative, so they are hypothesized to drive less overall neuronal activity to minimize uncertainty^2^. By contrast, surprising/unpredictable stimuli evoke prediction errors (ε, see Fig. 1a, Equation 1 for Hierarchical Predictive Coding), which enhance neuronal activity as a function of their magnitude and precision (Π), resulting in a precision-weighted prediction error (ξ). Numerous neuronal and computational instantiations for prediction and prediction error computations have been proposed, comprising a larger family of Predictive Processing models (representative examples are listed in Supplementary Table 1).

**Fig. 1.**
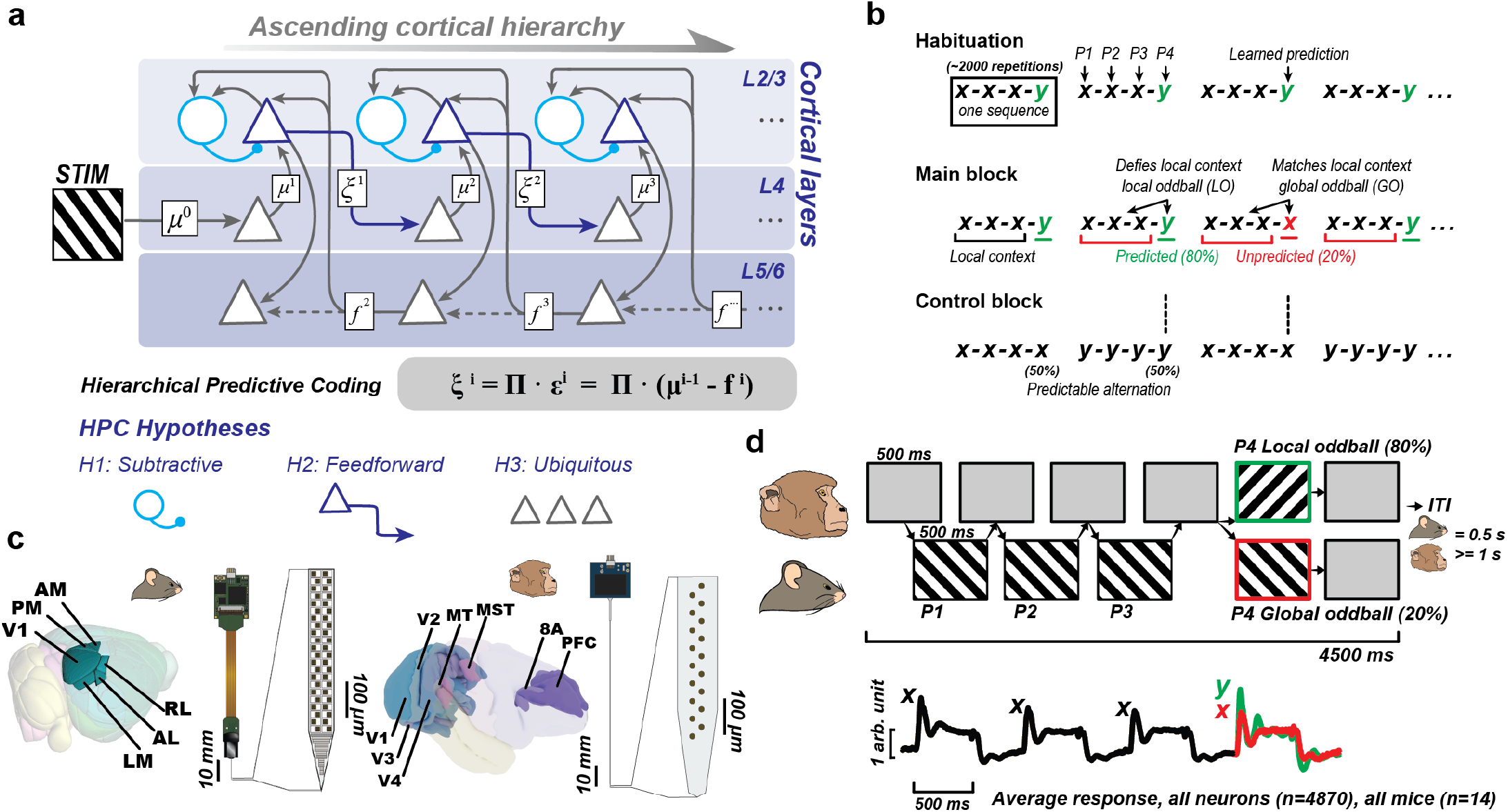
Hierarchical Predictive Coding (HPC) hypotheses, Local/global oddball paradigm, and experimental setup. **a**, Schematic for sensory processing via hierarchical predictive processing. Stimulus information enters the cortical hierarchy through layer 4, and prediction error is computed in superficial layers 2/3, with inhibitory cell involvement in the subtractive process (H1). Prediction error is then fed forward (H2) via superficial layers to higher-order areas. Deep layers in higher-order areas provide a feedback prediction to the previous stage. Error signals emerge early and propagate throughout the cortical hierarchy, utilizing this circuitry in a ubiquitous fashion. (H3) **b**, Visual stimuli were presented in 4-stimulus sequences consisting of 2-oriented, drifting gratings symbolized as ‘x’ or ‘y’. Animals were habituated to the x-x-x-y sequence (counterbalanced over sessions/animals with y-y-y-x). In the main block of a recording session (n=400 trials), animals were presented with 80% x-x-x-y and 20% x-x-x-x (global oddballs). Control block (n=200 trials) consisted of x-x-x-x (control for global oddball, 50% of trials) and y-y-y-y (control for local oddball, 50% of trials). **c**, Neuropixels were introduced into 6 visual cortical regions in all mice (V1, LM, RL, AL, PM, AM). 128-channel or 32-channel laminar probes were introduced into 8 cortical regions in 2 monkeys (V1, V2, V3, V4, MT, MST, 8A, PFC). **d**, Monkeys viewed sequences of drifting grating stimuli while fixating centrally. They were rewarded with liquid juice at the end of the trial for maintaining fixation. Mice viewed sequences of full-screen drifting grating stimuli. An Inter-Trial Interval (ITI) period was included between trials. The trace shows the population average response (all neurons in all mice) to the 4-stimulus sequences with the GO in red and the LO in green.

Sequence-based oddball paradigms have been widely used to test HPC and other PP models^9^. Subjects are exposed to a repeated pattern of sensory inputs to implicitly learn the hidden rules governing them. Studies of brain activity during oddball tasks using fMRI, MEG/EEG, and LFP have been largely consistent with HPC models^9–11,13,16–20^ and have documented neuronal signatures of prediction errors both in higher-order areas (e.g., prefrontal cortex) as well as sensory areas, including primary and secondary auditory and visual cortex (for a review, see^13^).

However, because it is difficult to unambiguously resolve feedback modulation from local computations and feedforward output^21^ using these approaches, prior work could not resolve key hypotheses made by the HPC models. So far, studies of spiking activity have been more equivocal, with different results depending on species, paradigm, cortical area, and methodology^3,13,20,22–26^. Importantly, no study has yet used large-scale high-density neuronal electrophysiological recording methods in multiple connected brain regions to test HPC models. Brain recordings with this level of detail are necessary to experimentally test HPC because these models propose multiple stages of cortical processing, with distinct cell types, cortical layers, areas, and directions of feedforward/feedback processing contributing to the hypothesized computations.

The simplest pattern that can create a predictive model is that the current stimulus will continue to repeat. A first-order pattern violation in this context is a change from repeated stimulation (e.g., during a stimulus sequence such as x-x-x-y, the “y” violates the local context established by repeating “x”)^13,27^. Neuronal responses to these stimuli are “local oddball” (LO) responses^9,10,13,28^. LO responses persist during deep anesthesia, when feedback connections are functionally wea^20^. As such, they do not provide sufficiently rich predictive context to test HPC, in which feedback was proposed to modulate early sensory processing through rich, multifaceted predictions^2,3,29,30^. We used second-order pattern violations to test more complex prediction error computations through higher-order predictions. To evoke higher-order predictions, we repeated the presentation of a first-order pattern (x-x-x-y / x-x-x-y / x-x-x-y) and violated the pattern with x-x-x-x, where the fourth “x” is called a “global oddball” (GO). Importantly, GO responses cannot be explained by recent stimulus history, i.e., release from adaptation.

In this work, using responses to LO and GO, we test three of the main hypotheses made by HPC models in the visual domain. The first hypothesis is that predictions are generated in higher-order areas from neurons in deep cortical layers (L5/6) and project via feedback connections to non-granular layers 1-3 and 5/6 of an earlier area in the hierarchy^31,32^. In the earlier hierarchical area, higher-order predictions are compared with and subtracted from sensory inputs (Fig. 1a, Hypothesis 1, “Predictions are suppressive or inhibitory”). This subtractive or inhibitory component of predictions is a consequence of Equation 1 (Fig. 1a), where bottom-up stimulus information (μ^i-1^) is subtracted from top-down predictions (f^i^) (a shared feature of HPC models in Rao and Ballard, 1999; Friston, 2010; Bastos et al., 2012, amongst others). The subtractive feedback predictions are thought to be implemented via the activity of local inhibitory interneurons in superficial layers 1-3^1,3,28^ (Fig. 1a). To implement predictive suppression, interneurons should increase their firing rates, resulting in a net decrease in activity of the population. Conversely, during prediction errors, inhibitory interneuron activity should decrease spiking activity, allowing the overall firing rate of the population to increase, signaling prediction error. In the second hypothesis, precision-weighted prediction errors (hereafter referred to as prediction errors for simplicity) are signaled in the feedforward direction, emanating from layer 2/3 pyramidal neurons of an earlier area and targeting layer 4 neurons of a higher area. They feed forward up the hierarchy to update internal models to make better predictions (Fig. 1a, Hypothesis 2, “Prediction Errors are feedforward”). In the third hypothesis, HPC models state that brains construct complex models that are compared to sensory inputs at multiple levels of cortical processing (Fig. 1a, Hypothesis 3, “HPC circuitry is ubiquitous in cortex”). Computational instantiations of HPC models have proposed that prediction error computations occur both in early sensory as well as higher-order levels of neuronal circuits, even for high-level predictions^11,27,33^.

We tested the above hypotheses regarding HPC using Multi-Area, high-Density, Laminar-resolved Neurophysiology (MaDeLaNe) recordings of thousands of spiking neurons in mice and monkeys throughout the visual cortex (areas V1, LM, RL, AL, PM, AM in mice and areas V1, V2, V3, V4, MT, MST in monkeys) and prefrontal cortex (areas 8A and lateral PFC in monkeys). We used a habituation-based no-report variant^20,34^ of the global-local oddball task^1^, thereby avoiding motor and reward related confounds on sensory responses. This paradigm allowed us to disentangle short-term stimulus repetition from prediction. Four surprising findings challenge the hypotheses made by HPC models. First, global oddballs evoked neuronal spiking responses consistent with a prediction error signal, but these responses were restricted to higher-order cortical areas in both species. Second, we used cell-type specific optogenetics in both species to test whether inhibitory interneurons performed predictive suppression as proposed. If so, inhibitory interneurons would be expected to increase their activity during a predictable stimulus, or equivalently to decrease their activity to an unpredicted stimulus, enabling the expression of a prediction error response when predictions are violated. Global oddballs did not modulate the inhibitory interneuron populations in this hypothesized manner. Third, highly expected local oddballs did not evoke a reduced neuronal response compared to the same sequence when it was contextually deviant. Fourth, global oddballs evoked a laminar- and area-wise pattern of activity more consistent with feedbackinvolving neuronal activity in L2/3 and L5/6 in higher areas - rather than the hypothesized pattern of feedforward processing (e.g., involving neuronal activity in lower areas feeding forward to layer 4 in higher-order areas).

## Results

### Isolating sensation and establishing priors

We first extensively habituated animals to the local oddball x-x-x-y stimulus sequence for 2000-3000 trials over several days (Fig. 1b and Extended Data Fig. 1). The “x” and “y” visual stimuli within a sequence denote drifting gratings at 135° or 45° from horizontal and were counterbalanced across animals. The unpredictable global oddball x-x-x-x stimulus sequence was never presented during habituation and introduced only on neural recording sessions (Fig. 1c), on 20% of trials in the main block (Fig. 1b). This extensive habituation ensured that animals could learn to predict the x-x-x-y (local oddball) stimulus sequence. Pupillary responses in monkeys (Extended Data Fig. 2) and running speed changes in mice (Extended Data Fig. 3) indicated that animals behaviorally registered the violation of the habituated sequence during unpredicted x-x-x-x global oddballs during the recording sessions.

To investigate the neuronal signaling underlying these predictions and their violations, we compared responses in the main block to a control block where the x-x-x-x sequence alternated with y-y-y-y and thus the sequence was predictable after the first stimulus (Fig. 1b). To analyze global and local oddballs, we compared spiking responses to the fourth component in the sequence (P4) to the same stimulus in the same sequence position of the control block (vertical dotted lines in Fig. 1b). To account for cross-session firing rate drift which occurs even in well-isolated single units^35^, we normalized responses to the third component in the sequence (P3). Our main contrast was therefore P4-P3 in the main block vs. P4-P3 in the control block (see *Methods*). For global oddballs, this contrast should reveal prediction errors while controlling for short-term adaptation. For local oddballs, this contrast should reveal the release of adaptation caused by changing the stimulus after a few repetitions. Neurons responded to these sequences with robust firing rate increases in all recorded areas (Fig. 1d, Supplementary Figs. 1-3). Significant oddball detection was determined by comparing the neuronal spiking response for each oddball type using a nonparametric, cluster-based permutation test^36^ (at P < 0.05, corrected for multiple comparisons, see *Methods*).

### Predictable local oddballs are widely signaled

Because of extensive habituation, local oddballs were non-surprising stimuli and, according to HPC models, should be suppressed and explained away (H1). Despite their extensive habituation, we observed that neuronal signaling of local oddballs was ubiquitous (present in all areas and both species, Fig. 2a), signaled early and fed forward up the visual cortical hierarchy (within the first ~150 ms of stimulus processing, Fig. 2b), strong (local oddball responses by area were on average 60-98% above the control stimulus in mice, and 43%-93% in monkeys, Fig. 2c), and involved more than 50% of recorded neurons (Fig. 2d). To ensure that these local oddball responses at the neuronal population level were not a consequence of simple stimulus selectivity differences in the population, we ensured that each neuronal population we tested contained just as many ‘x’ as ‘y’ preferring neurons and where there were imbalances, performed stratification (Supplementary Table 2, Supplementary Fig. 4, see *Methods*). In mice but not monkeys, local oddball signaling gradually increased in strength with ascending cortical hierarchy (Linear correlation between local oddball response and hierarchical area, R^2^ = 0.83, P=5.75e^−5^, Fig. 2c, left subpanel). In V1 of both species, local oddball signaling was most prominent amongst neurons in L2/3 neurons (Fig. 2e; for details on layer identification, see *Methods* and Supplementary Fig. 5).

**Fig. 2.**
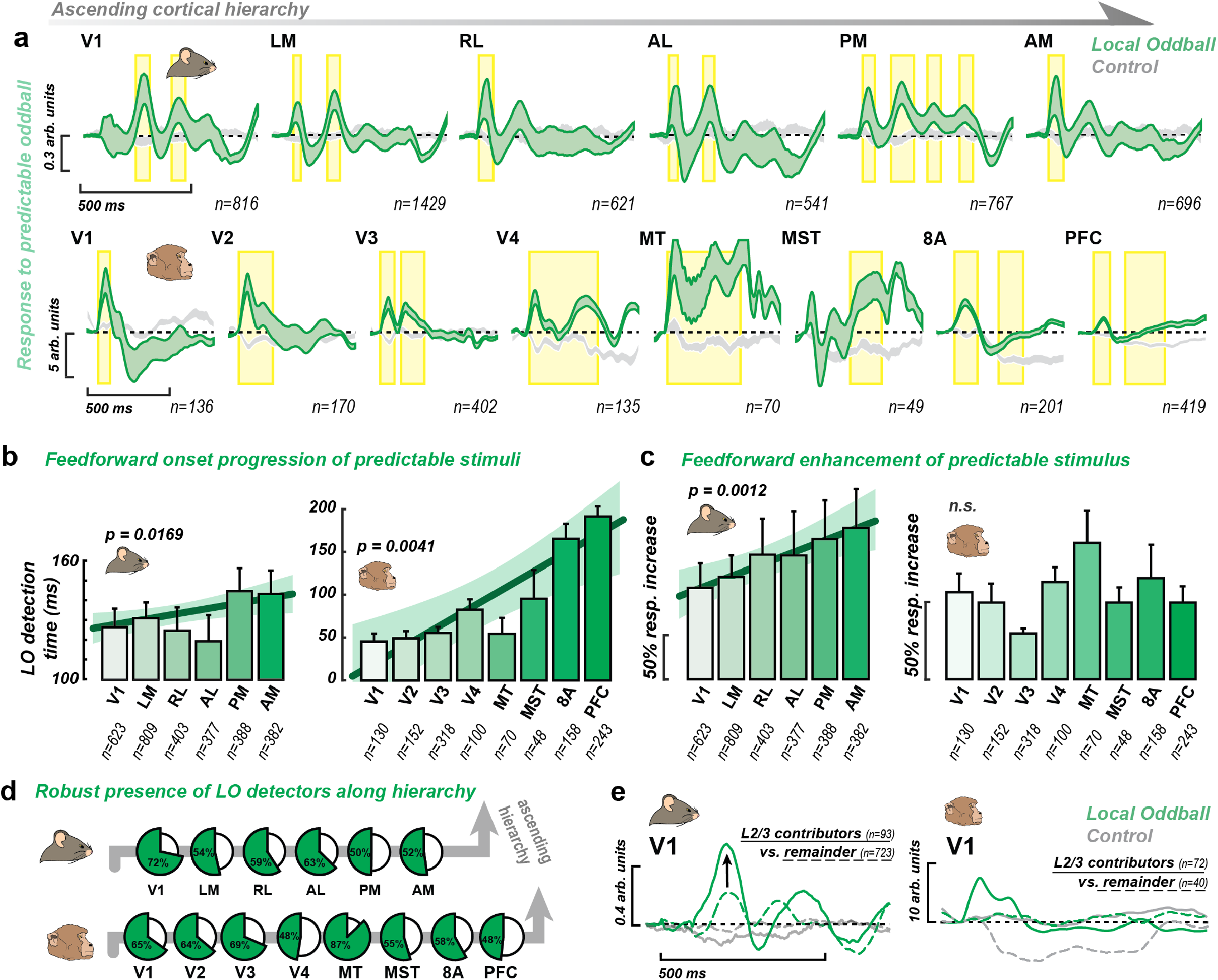
Predictable local oddball response is widespread, emerges early, and feeds forward. **a**, Local oddball detection across cortical areas in mice and monkeys. Bands are 95% confidence intervals across units in an area. Unit responses are normalized by dividing instantaneous firing rate by average firing rate at the single-unit level in mice and via a z-score method for the multiunit data in monkeys (see *Methods*). Only visually responsive units are included. Green bands are the P4-P3 local oddball in the main block; gray bands are the P4-P3 local oddball in the control block. For non-subtracted neuronal activity, see Supplementary Fig. 1. Yellow highlights reflect significant population local oddball detection periods, P<0.05, corrected using nonparametric, cluster-based permutation tests. Local oddball signaling in mice was accompanied by a stimulus-induced temporal modulation corresponding to the drifting grating presentation rate of 4 Hz (upper subpanel). This oscillatory component was much less prominent in monkeys (lower subpanel). **b**, Average onset time for significant local oddball effect across hierarchy in mice and monkeys. Error bars indicate 2 SEM across units. Linear regression shows significant relationship between hierarchical position and onset timing (R^2^ = 0.83, P=0.006 in mice; R^2^ = 0.88, P= 0.004 in monkeys), indicating a feedforward progression. Units that had a contiguous 10ms significant local oddball response were included in this analysis. **c**, Percent response increase to local oddball across cortical areas in mice and monkeys over all units (regardless of visual responsiveness) in each area. Error bars indicate 2 SEM across units. Local oddball effects show significant feedforward enhancement in mice along the cortical hierarchy. **d**, Percentage of local oddball encoding units along cortical hierarchy in mice and monkeys. Signaling of predictable local oddballs is robust (>50% of all cortical areas) and widespread (observed in all cortical areas) **e**, Local oddball responses are more pronounced in L2/3 in V1 in both mice and monkeys compared to units in other layers, suggesting feedforward signaling.

Can these local oddball responses, representing local signal change, be considered a simple form of prediction error signaling? To examine this, we compared the local oddballs in contexts that differed in their relative probability (local oddball sequence probability varied between 100%, 80%, and 12.5%, see Extended Data Fig. 1). If they represent prediction errors, local oddball responses should scale according to their deviance (H1). We found that local oddball signaling did not scale as a function of deviance in most areas (Extended Data Fig. 4, area V4 was an exception). Instead, we observed the opposite pattern just as often, and most prominently in V1 of mice (Extended Data Fig. 4). The enhanced neuronal responses to more predictable local oddballs in mouse V1 may have been caused by learning-driven potentiation^37^. In addition, local oddball responses did not change as a function of position in the main block in monkeys and were reduced slightly across the main block in mice (Supplementary Fig. 6). To summarize, local oddball sequences (x-x-x-y) release most neurons from adaptation and increase excitability independent of prediction, even though animals had been exposed to these sequences thousands of times.

### Unpredictable global oddballs do not generate feedforward prediction error

Next, we tested whether highly surprising global oddballs (x-x-x-x) drive neuronal spiking consistent with prediction error signaling. We found that global oddball signaling was restricted to only a few higher-order brain areas (LM, AM, and PM in mice; V3, MT, 8A, and PFC in monkeys, Fig. 3a). In monkeys, extensive habituation to the x-x-x-y sequence was necessary to evoke global oddballs in area MT and significantly boosted global oddball responses in 8A and PFC compared to without habituation (Extended Data Fig. 1d,e). This suggests that global oddballs developed because of habituation-induced learning. In the main habituated paradigm, at the individual neuron level, although we identified a handful of neurons and areas that responded to global oddballs (Supplementary Fig. 2), the percentage of units signaling global oddballs in each area was sparse (median across areas in mice: 7%, median across areas in monkeys: 8%, Fig. 3b), contradicting H3 (“Predictive coding circuitry is ubiquitous”). Unlike local oddballs, which showed a clear temporal progression of latencies across the hierarchy (Fig. 2b), the latency at which neurons signaled global oddballs did not scale with hierarchy (Fig. 3c). Instead of being a prominent part of the sensory response, global oddball responses were more often seen long after visual stimulation began (>250 ms post-stimulus) or even in the off-response periods which were periods with a blank screen, devoid of active sensory processing (Supplementary Fig. 7).

**Fig. 3.**
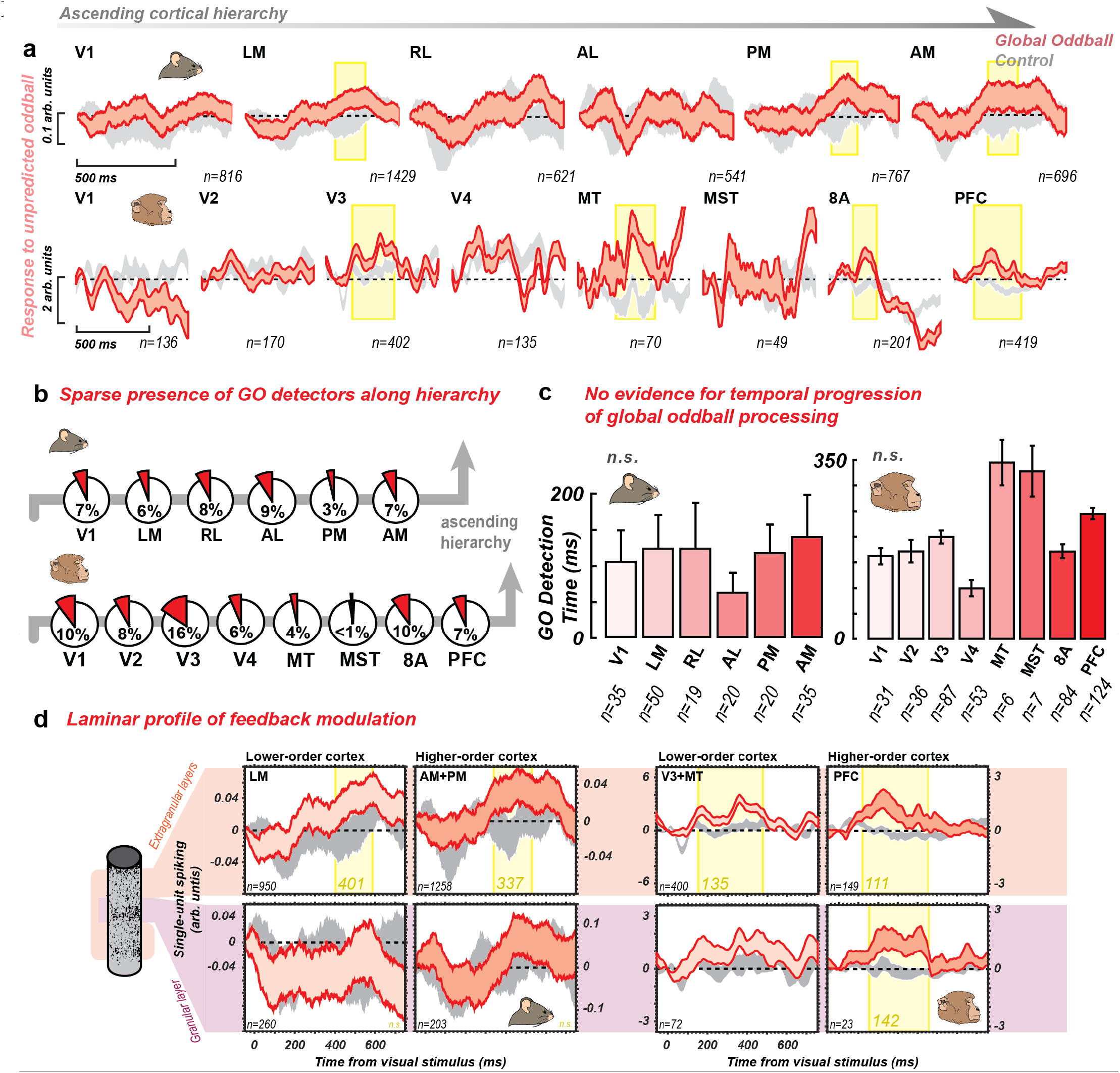
Unpredictable global oddball emerges in higher-order cortex and is fed back. **a**, Global oddball detection across cortical areas in mice and monkeys in visually responsive neurons. Unit responses are normalized by dividing instantaneous firing rate by average firing rate at the single-unit level in mice and via a z-score method for the multiunit data in monkeys (see *Methods*). Only visually responsive units are included. Bands are 95% confidence intervals across units in an area. Red bands are P4-P3 in the GLO block; gray bands are P4-P3 in the control block. For non-subtracted neuronal activity, see Supplementary Fig. 1. Yellow highlights reflect periods of significant population global oddball detection, P<0.05, corrected for multiple comparisons using nonparametric, cluster-based permutation tests. N’s in the figure indicate the number of single units (mice) or multi-units (monkeys) per analysis recorded across n=15 mice and n=2 monkeys. **b**, Percentage of significant global oddball encoding units/channels along the cortical hierarchy in mice and monkey. The percentages reflect units with significant response to global oddball over all units in each area (regardless of visual responsiveness). **c**, Onset times of significance for global oddball effect along the cortical hierarchy in mice and monkeys. Linear regressions show no significant change along cortical hierarchy, suggesting no strong feedforward signaling of global oddball prediction error. Units that had a contiguous 10ms significant global oddball response were included in this analysis. **d**, Granular versus extra-granular global oddball detection in areas with significant population-level detection in mice and monkeys. Red bands are P4-P3 in the GLO block; gray bands are P4-P3 in the control block. Yellow highlights reflect periods of significant population global oddball detection in the layer grouping, with the latency indicated.

We next examined global oddballs within L2/3 pyramidal neurons, which according to multiple PP models should transmit prediction error (H2). In mice, we found that putative layer 2/3 pyramidal cells did not signal global oddballs (Extended Data Fig. 5). In addition, current source density analysis of synaptic activity in L2/3 of mice did not reveal any reliable changes in synaptic activation (Extended Data Fig. 6) during global oddballs. In monkeys, L2/3 spiking activity did not signal global oddballs in area V3 (Extended Data Fig. 5c). There was L2/3 involvement in global oddballs in area MT, but this came 116 ms later than in area PFC, supporting a feedback^38,39^ rather than feedforward putative error computation (Extended Data Fig. 5c).

Without the hypothesized L2/3-specific signal, we investigated alternative sources for global oddball signaling. We limited the analysis to the areas with population global oddball signaling. We grouped areas into higher- and lower-order (higher-order areas AM and PM vs. lower-order area LM in mice; higher-order area PFC vs. lower-order areas V3 and MT in monkeys). We first divided the units into granular (L4) vs. extragranular (L2/3 and L5/6) compartments, which anatomically separate feedforward vs. feedback input^31,32,40^. In mice, we observed global oddball signaling only in the extragranular (feedback-associated) layers and later in the lower-order cortex (401 ms post-stimulus, Fig. 3d, left subpanels) than in the higher-order cortex (337 ms post-stimulus). In monkey visual areas V3 and MT, we observed global oddball signaling only in the extragranular (feedback-associated) layers (135 ms post-stimulus), later in time than the extragranular layers of higher-order area PFC (111 ms post-stimulus, Fig. 3d, right subpanels). These latency and laminar effects collectively support prediction error propagation from higher to lower areas of the hierarchy, contradicting H2.

Having not found the hypothesized signatures of prediction error coding of GOs, we next turned to a data-driven analysis. We examined whether global oddball signals may have been present in a more complex neuronal ensemble that may have been missed by our hypothesis testing. We applied functional clustering (see *Supplementary Text*) to the single unit spiking data in mice and tested whether global oddball encoding properties emerged in a well-defined cluster. We did not find evidence for any clusters specialized in GO detection (Supplementary Fig. 8).

Finally, we also analyzed whether global oddballs were more consistently signaled in the first few trials when they occurred early in the main block, when putative prediction errors should be strongest. Although area V4 in monkeys and PM in mice exhibited a profile consistent with a putative prediction error signal, areas before or after them in the hierarchy did not (Supplementary Fig. 9), inconsistent with the hypothesized feedforward (H2) or ubiquitous (H3) prediction error signal. Robust global oddball signaling also did not emerge when we analyzed only later periods in the control trials, allowing for more time to form predictions of the expected x-x-x-x stimulus in the sequence control (Supplementary Fig. 10).

### Directed connectivity of global oddballs does not indicate feedforward error propagation

In HPC models, prediction errors are sent feedforward (Fig. 1a, H2). Hence, neurons earlier in the hierarchy should increase their drive onto neurons later in the hierarchy to signal prediction error. We tested this by computing Granger causality between spiking activity time courses. Granger causality tests whether neuronal activity in area A can statistically predict activity in area B above and beyond predictions made by activity in area B alone^41^. If so, then area A “Granger-causes” B. By comparing the A-to-B vs. the B-to-A directions of Granger causality^42^ where A is below B in the hierarchy, we tested whether local and global oddballs evoked feedforward processing. We performed this analysis in mice, because recordings across areas were performed simultaneously, which is a pre-requisite for Granger calculation. In mice, compared to pre-stimulus baseline (Extended Data Fig. 7), the hierarchy became more feedforward-dominated in both early (100-300 ms) and late (300-500 ms) periods of local oddball processing (Extended Data Fig. 7a,b; Wilcoxon rank sum test, P<0.01). This feedforward activity propagated from V1 to the highest levels of the visual hierarchy, areas PM and AM. In contrast, during global oddballs, there was no change in feedforward vs. feedback asymmetry compared to baseline (Extended Data Fig. 7b-d, Wilcoxon rank sum test of GC asymmetry values across mice vs. pre-oddball baseline, all comparisons, P>0.01). To summarize, the latency, hierarchical areas of significant signaling, laminar, and Granger analyses collectively contradicted H2; namely, global oddballs did not drive feedforward processing. Instead, the data on hierarchical areas of significant signaling and laminar position supported a feedback model of prediction errors evoked by global oddballs.

### Global oddballs: A release from predictive inhibition?

In HPC models, predictions are hypothesized to subtract away predictable stimuli (H1), leaving sensory cortex less excited. During prediction error, this should result in a net decrease in inhibitory interneuron firing rates, allowing the overall population firing rate to increase and thereby report prediction error. We tested this by assessing whether predictions are neuronally instantiated by inhibitory interneuron activity. We used optogenetics to identify Somatostatin-(SST) and Parvalbumin-(PV) expressing interneurons in mice, viral vectors that restricted optogenetic expression to inhibitory interneurons in monkeys^43^, and waveform shape analysis to identify putative inhibitory interneurons in monkeys^44^ (Fig. 4a and Extended Data Fig. 8). In both species, we utilized Channelrhodopsin-2 to drive inhibitory interneurons to fire during light stimulation (see *Methods*). In mice, PV+/SST+ units were identified across all recorded areas. Inhibitory interneurons were identified in areas MT/MST, PFC, and 8A (Fig. 4b) in one monkey. Opto-tagged neurons in the monkey had, on average, narrow waveforms which is additional confirmatory evidence^45^ that these opto-tagged neurons were inhibitory (Fig. 4b; Extended Data Fig. 9). In both species, these inhibitory cell populations responded to the stimulus sequence and increased their spike rates during P4 of local oddballs (Fig. 4c). According to H1, inhibitory interneurons within areas that showed global oddball responses should decrease their activity to allow for prediction errors to be expressed. We did not observe a significant response to global oddballs at the population average of these subpopulations across areas combined (Fig. 4d). When considering individual areas and laminar compartments in mice, we found that PV neurons when combined in areas LM, PM and AM (areas which showed global oddball responses at the population level) and in individual areas LM and PM and in deep layers (Extended Data Fig. 10) increased firing rates during global oddballs – the opposite direction of the hypothesized effect. SST neurons did not respond to global oddballs in any area/laminar compartment (Extended Data Fig. 10). In the monkey, we also bolstered the number of neurons in the analysis by considering putative inhibitory interneurons defined by their narrow waveform shape (Fig. 4e). These putative inhibitory interneurons also did not respond to global oddballs, neither when considering the lower-order areas with a population global oddball response (V3 and MT, Fig. 4e, top) nor when considering the higher-order areas with a population global oddball response (8A and PFC, Fig. 4e, bottom, see also Extended Data Fig. 9). Inhibitory interneurons selective for the global oddball stimulus also did not signal global oddballs in mice (Supplementary Fig. 11) or in monkey (for putative inhibitory interneurons, Extended Data Fig. 9d). Collectively these analyses indicate a minor role, if any, for these inhibitory interneuron sub-populations in releasing predictive suppression to enable global oddball signaling, failing to support H1.

**Fig. 4.**
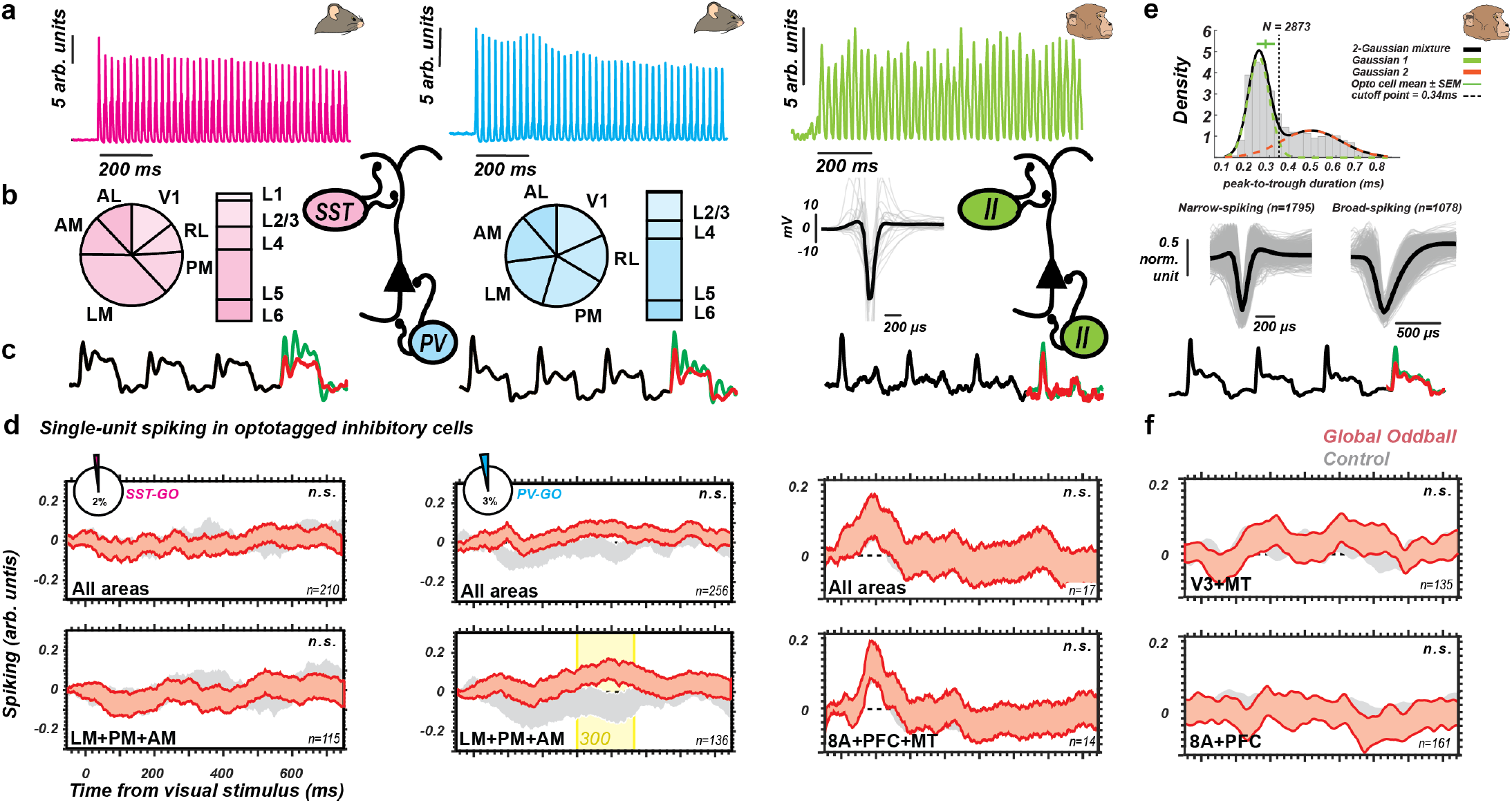
Optotagged inhibitory cells in mice and monkeys do not carry out hypothesized predictive coding functions. **a**, Optical stimulation with 40 Hz sinusoidal modulation of laser power. Tagged SST (magenta, n=215 neurons from n=9 mice) and PV (cyan, n=262 neurons from n=7 mice) responses are normalized by their average firing rate. In primates, an opsin targeting all GABAergic inhibitory cells was used; the responses shown were computed as a weighted mean (green, n=17 neurons from n=1 monkey), with each cell weighted by its modulation strength (average firing rate change from baseline, divided by the sum of the baseline and laser-timed response). **b**, Relative proportion of each inhibitory interneuron type, which have synapses at largely different sites on pyramidal cells (schematized here), across areas and layers in mice (left and middle). Average waveform shape and spike width in primate data (right) are consistent with those of narrow-spiking interneurons. **c**, Population average spiking response to the oddball sequences for these inhibitory interneurons. P1 to P3 in black, local oddball P4 in green, and global oddball P4 in red. **d**, Lack of global oddball detection in SST and PV cell subpopulations in mice and in pan-inhibitory cell populations in monkeys. Bands are 95% confidence intervals across units. Top sub-panels, the red band is P4-P3 in the main block; the gray band is P4-P3 in the control block for all neurons of the specific type. For P3 and P4 before subtraction, see Supplementary Fig. 1b. Bottom sub-panels: same as top sub-panels, but restricted to the areas that showed significant global oddball signaling at the population level. **e**, The histogram shows the distribution of all good units from both monkeys (n=2) from the main experiment and the optotagging experiment. A two-Gaussian mixture fit yielded a cutoff of 0.34 ms, which was used to identify putative inhibitory cells (waveform duration < 0.34 ms). An indication of the waveform duration from the monkey optotagged units (n=17) is shown above the histogram in green. Putative inhibitory cells were identified using waveform duration validated by the waveform durations of optotagged inhibitory cells. Cells with wide waveforms (right) were identified as putative excitatory cells, while cells with narrow waveforms (left) were identified as putative inhibitory cells. **f**, Identical to d, but with the putative inhibitory cell population of visual sensory areas V3 and MT (top) and the putative inhibitory cell population of higher-order areas 8A and PFC (bottom). For P3 and P4 before subtraction, see Extended Data Fig. 9.

### HPC and Predictive Processing model-data comparisons

The existing literature on models of classical HPC and a broader family of PP models is large, with many mechanistic proposals. To contextualize our experimental results within the broader corpus of these HPC and PP studies, we performed a survey of our results against claims from a representative set of previous experimental and theoretical works. To quantify the distance between our results and the literature, we utilized an agent-based reasoning system, leveraging the recent capabilities of Large Language Models^46^. We created a pipeline^47^ whereby a council of ten models (Supplementary Table 3) performed quantitative scoring of our results and the corpus of studies in Supplementary Table 1 on each study’s agreement (assigned a score of +1) vs. disagreement (assigned a score of −1) on each of the three hypotheses (defined by a series of factors, see Supplementary Tables 5-7). The models performed this scoring separately for local and global oddballs (see *Methods* and Supplementary Table 4). We found that for LOs, our paper diverged significantly from this literature on H1 (Fig. 5, green, Subtractive predictions, M=-0.13 SD=0.35 vs Literature M=0.53 SD=0.10, P=0.0005, Wilcoxon rank sum test) but not on H2 (Feedforward prediction errors, M=0.43 SD=0.38 vs Literature M=0.62 SD=0.10, P=0.2762, Wilcoxon rank sum test) nor H3 (Ubiquity, M=0.52 SD=0.35 vs Literature M=0.42 SD=0.14, P=0.2331, Wilcoxon rank sum test). This aligns with our findings of feedforward mechanisms throughout the cortical hierarchy in both species for local oddballs, but also that these local oddballs were not enhanced by their relative deviance (Extended Data Fig. 4). For global oddballs, our findings significantly diverged from the literature on all three hypotheses (Fig. 5, red, H1, M=-0.532 and SD=0.292 vs. Literature M=0.486, SD=0.141, P<1e-3, H2, M=-0.531, SD=0.284 vs. Literature M=0.437, SD=0.310, P=1e-4, H3, M=-0.158, SD=0.552 vs. Literature M=0.430, SD=0.147, P=0.01, Wilcoxon rank sum test). This aligns with our findings that global oddballs did not release predictive suppression (Fig. 4), did not evoke feedforward mechanisms (Fig. 3c,d, Extended Data Fig. 7) and were sparse, weak, and absent in half of the areas of the hierarchy (Fig. 3a,b). We graphically summarize the major findings of our study in Fig. 6.

**Fig. 5.**
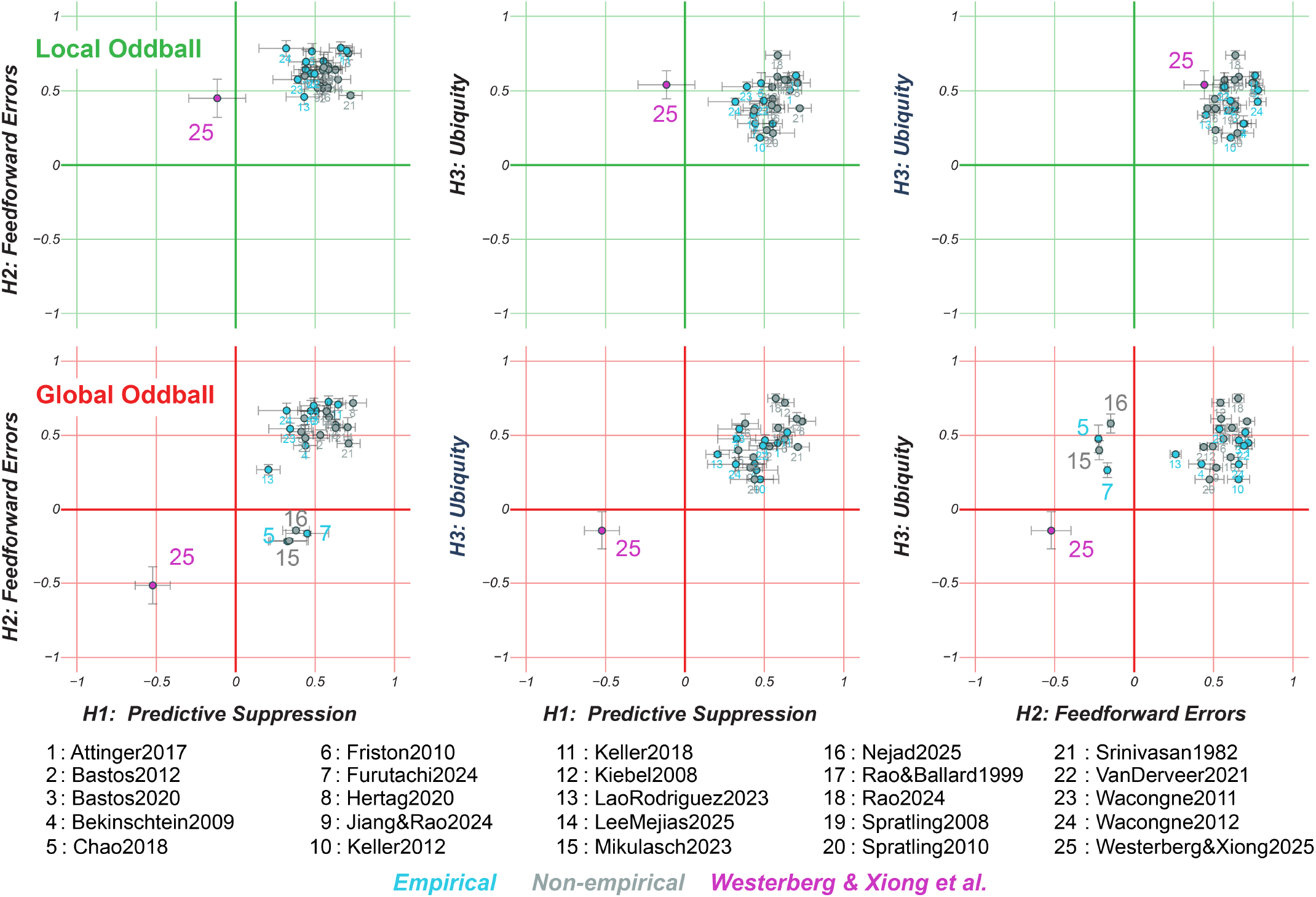
Model evaluations of the literature relative to our findings. Axes: Linear scales from −1 to 1 quantifying study alignment with the hypotheses. Negative values indicate disagreement; positive values indicate agreement. The metrics include H1: Predictive Suppression, H2: Feedforward Errors, and H3: Ubiquity. Individual publications correspond to the dots, are indexed in the study legend and in Supplementary Table 1. Green dots represent empirical studies, grey dots represent non-empirical papers, and the red dot (25) represents this work. Error Bars indicate SEM for each respective study evaluation across independent evaluations from n=10 distinct LLMs (see Supplementary Table 3) performed separately for LO and GO contexts (defined in Supplementary Table 4). For additional details see *Methods*.

**Fig. 6.**
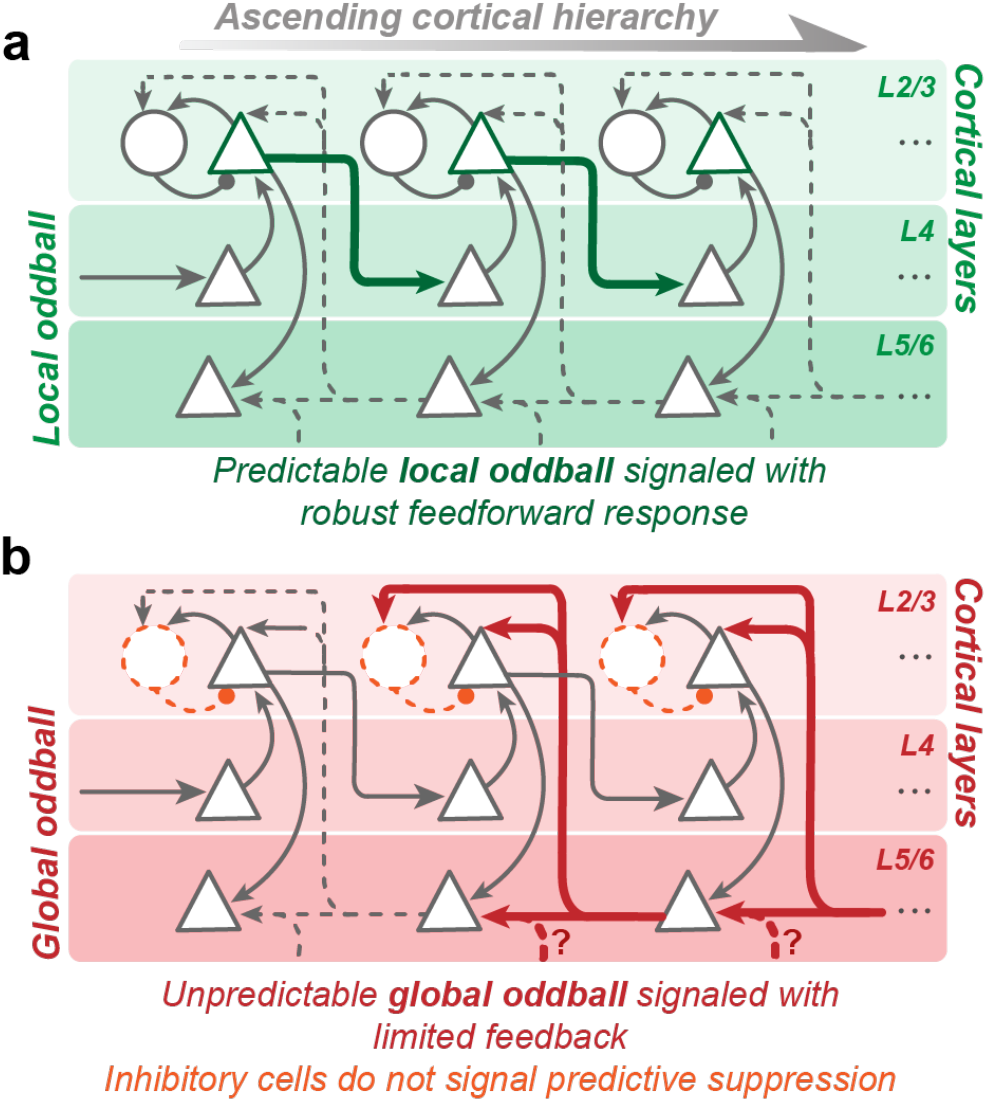
Schematic of findings represented in the canonical cortical connections. **a**, Highly predictable local oddball resulted in robust and widespread feedforward response. The timings of onset of significance as well as Granger causality results suggest strong feedforward processing. Thick green arrows represent strong cortical activity across layers and widespread across cortical hierarchy. **b**, Unpredictable global oddball resulted in a limited feedback response, particularly in select higher order areas. We did not observe L2/3 prediction error signals in mice and found non-specific L2/3 (compared to L5/6, except for in PFC at top of the hierarchy) prediction error signals in monkeys. Furthermore, optotagged inhibitory cells do not exhibit suppression specific to predictable stimuli. Question marks signify that additional areas we did not sample may contribute to GOs.

## Discussion

We tested several key hierarchical predictive coding (HPC) model hypotheses by performing an extensive survey of cortical spiking in mice and monkeys observing predictable or unpredictable stimulus sequences. We used MaDeLaNe (Multi-Area, high-Density, Laminar Neurophysiology) recordings^13^ and optogenetics to isolate different cortical layers, areas, and neuron types previously hypothesized to play a role in HPC^1–3,48^ (Fig. 1a). We tested whether neuronal spiking responses could be described as prediction error signals resulting from a release from predictive suppression (H1) that triggers feedforward processing (H2) with a ubiquitous signature (H3). HPC models and a wider family of predictive processing (PP) models and prior work based on theoretical, computational, and empirical perspectives largely agreed with these three core hypotheses (with some exceptions, see Fig. 5).

HPC models^1–3,48^ hypothesize that prediction errors, that is the difference between the bottom-up sensory signal and the top-down prediction, constitute a core, feedforward cortical computation (a hypothesis shared by many models in the PP model family, Supplementary Table 1, Fig. 5). Our subjects were habituated to a particular sequence (x-x-x-y) for thousands of trials such that its violation (x-x-x-x) should drive robust spike signaling of prediction error. In HPC, the habituated (and highly predictable) local oddballs (x-x-x-y) should be suppressed, driving less activity^49^. We found the exact opposite: spiking responses robustly signaled unsurprising local oddballs (engaging over 50% of all recorded neurons throughout cortex) but weakly signaled surprising global oddballs (engaging less than 10% of all recorded neurons). Signals associated with local oddballs emerged early in the hierarchy at fast latencies in layers 2/3 and fed forward up the cortical hierarchy (Fig. 6a). However, contradicting the interpretation of these signals as prediction errors, local oddball responses did not consistently scale with increased sequence deviance. Further, global oddballs did not emerge because of a release of inhibition from PV/SST+ interneurons in mice or inhibitory interneurons in monkey (contradicting H1). Therefore, local oddball detection is most likely a release from neuronal adaptation. Although this robust local oddball detection is not a prediction error in the classic sense (e.g., according to HPC models), it could function to signal local deviations from context, and the magnitude of deviation may depend on how many local repetitions have occurred. This could in principle encode local stimulus probability.

HPC models hypothesized that L2/3 pyramidal neurons transmit feedforward prediction errors. Responses to global oddballs, which should have triggered prediction error responses with this hypothesized feedforward flow, emerged late in the sensory response and in higher-order areas first, then lower-order regions (Fig. 6b). Putative excitatory pyramidal neurons in L2/3 (which project feedforward connections) and their associated current sinks did not signal global oddballs in mice and only signaled weakly in one area (monkey area MT) and later than in PFC. Instead, global oddballs were found to emerge at the population level in the non-granular, feedback-recipient layers. Feedforward neuronal communication assessed by Granger causality amongst spiking neurons was not observed during global oddballs. Therefore, we failed to detect feedforward processing during global oddballs by four independent metrics: hierarchical area, temporal order, Granger causality, and laminar compartment of spiking/transmembrane current flow (Fig. 6b, contradicting H2). Instead, we found that global oddballs were stronger in higher areas of the hierarchy and in agranular layers which processed the global oddballs initially at higher-order areas and only afterwards in lower areas of the hierarchy. These observations support a feedback model of global oddball signaling. Our results speak against the proposed ubiquity of prediction error signaling in sensory cortex (contradicting H3). Altogether, our results challenge multiple aspects of current HPC models^1–3,48^ especially for global oddballs (Figs. 5,6).

It is worth considering whether the lack of explicit task engagement in our no-report task limited our ability to detect global oddballs. However, given the implicit behavioral changes (pupillary changes in monkeys, running speed in mice) we detected (Extended Data Figs. 2,3), it is likely that the animals registered the global oddball stimuli as deviants. In addition, global oddballs were observed in monkeys’ prefrontal areas and higher-visual regions of both mice and monkeys. Thus, despite being available to the animals’ higher cortex and evident in behavior, they did not profoundly affect responses in visual cortex. Also, some HPC models have postulated that prediction error signals are modulated by a gain term that takes on a functional interpretation in terms of precision (Π in Fig. 1a), which can be modulated by attention. That being said, models that consider gain in terms of precision require that precision be positive and nonzero^50^, entailing that prediction error signals might be attenuated but never entirely quashed to zero by gain modulation. In addition, in some formulations, prediction errors are thought to be automatic features of the sensory-driven response and do not explicitly depend on gain^1^. Future studies should explicitly consider the role of gain modulation on prediction error responses by manipulating the attentional state.

We found that global oddball responses are largely a property of neurons in higher-order cortex (PFC in primates, and late stages of visual processing in both species). These higher-order areas, in which we found population level coding of global oddballs^13,20,34,51^, contain neurons with long timescales^52^ and therefore in a better position to represent the relatively long duration of a sequence and its context, driving predictions and prediction errors in this task. This property may also explain why we found neuronal signaling of global oddballs more prominently in the period after stimulus offset (in monkeys), which may be related to representing the violated context within working memory^53^.

Our failure to observe global oddballs signaling in multiple areas may be related to the intrinsic properties of neurons in early visual cortex responding to changes in the environment with fast intrinsic timescales.^52^ They therefore may be ill-suited for predictive computations that require flexible re-mapping of responses and contextual processing over longer time scales. Neurons in higher-order areas are better positioned to integrate longer timescales. These higher-order neurons can re-map their activity in real-time based on experience-dependent learning and display mixed-selectivity. We speculate that this property is necessary for predictive processing because predictions need to be sensitive to the statistical structure of a changing and context-dependent environment^11,21,54^. We hypothesize that mixed selectivity neurons flexibly remap cognitive spaces to signal predictions. Functionally, such neurons could form dynamic ensembles via neuronal oscillations^20,55^ to guide sensory processing.

Although our empirical findings failed to support key aspects of classical HPC models (Figs. 5,6), some PP models hypothesized circuit elements were supported by our data. For example, Nejad et al.^5^ propose that both deep and superficial layers cooperate for prediction error computations, an effect highly aligned with our laminar results. Other works in computational neuroscience considered how top-down error signals could instantiate predictive learning^55^. Our work is consistent with these proposals and with a feedback direction of information flow for prediction errors^5,10,56^ as well as feedback associated enhancements of neuronal activity via attention^57^, which may have been triggered by the global oddballs. Higher-order areas participated in PP, although the computation was sparse across neurons, suggesting these areas may build flexible prediction in real-time^20,34,58,59^ along sub-spaces of neuronal activity (rather than engaging prediction errors widely as a core, cortex-wide computation).

Finally, not all aspects of the classical HPC models have been tested here. We studied predictions in the temporal domain because a significant body of prior work (Supplementary Table 1) had emphasized temporal oddballs as a key marker for predictive coding. Whether spatial and/or sensory-motor based predictions^8^ operate through the circuitry envisioned by HPC models remains to be investigated. Furthermore, we did not investigate all inhibitory cell types, (e.g., VIP+ neurons^60^) in our study. Finally, our LLM-based comparison between our results and the literature should be complemented in future work by quantitative computational modeling of the proposed predictive computations and comparison to data. With our full mouse and monkey datasets openly available, this will be a rich resource as a community effort to establish the PP principles and circuitry that are more consistent with MaDeLaNe data. Using novel approaches to model building^61^ we suggest that PP models^8^ should be constrained and grounded in these and other emerging MaDeLaNe neuronal recordings^12,13,62^.

## Acknowledgments

This work was funded by the US National Institutes of Health (NIH) [grant numbers: R00MH116100 (AMB), U24NS113646 (JAL, ChK)], the Dutch Research Council (NWO) [grant number: VI.Veni.232.110 (JAW)], the International Human Frontier Program Organization (HFSPO) [grant number: LT0001/2023-L], the Vanderbilt Faculty Fellow Award (AMB), and Vanderbilt University Startup Funding (AMB). We thank Andrew Salyer for assistance with literature review. The Neuropixels dataset was obtained at the Allen Brain Observatory as part of the OpenScope program, which is operated by the Allen Institute, Neural Dynamics program. We thank the OpenScope steering committee for their support, Karel Svoboda, and the Allen Institute founder, Paul G. Allen and Jody Allen, for their vision, encouragement, and support.

## Author contributions

Conceptualization: JAW, AM, AMB; Data curation: JAW, AB, SD, BH, JAL; Formal analysis: JAW, YSX, ES, MA, AMB; Funding acquisition: JAW, ChK, JAL, AMB; Investigation (NHP studies): JAW, YSX, HN, AMB; Investigation (rodent studies): SD, BH, HC, HB, HL, WH, KN, VH, TJ, CG, AY, JS, RG, BO, SC, AW, PAG; Project administration: JAW, CaK, JAL, AM, AMB; Software: JAW, AB, CRP; Supervision: SRO, ChK, JAL, AM, AMB; Validation: JAW, AB, CRP, JAL; Visualization: JAW, YSX; Writing – original draft: JAW, AM, AMB; Writing – review, and editing: all authors

## Supplementary information

Methods included below. Supplementary information includes Supplementary text, Supplementary references, Supplementary Tables 1-7, and Supplementary Figs. 1-11. See also Extended Data Figs. 1-10.

## Methods

### Mouse experiments at the Allen Institute: Animals

All experiments were performed in 16 SSTAi32 and PVAi32 mice (*Mus musculus*) of both sexes (n_male_=6, n_female_=10), aged 95-128 days. Optogenetics experiments were conducted with Sst-IRES-Cre/wt; Ai32(RCL-ChR2(H134R)_EYFP)/wt mice (n=9) and Pvalb-IRES-Cre/wt; Ai32(RCL-ChR2(H134R)_EYFP)/wt) mice (n=7). Pvalb-IRES-Cre and Sst-IRES-Cre mice were bred in-house and crossed with an Ai32 channelrhodopsin reporter line. Pvalb-IRES-Cre;Ai32 breeding sets (pairs and trios) consisted of heterozygous Pvalb-IRES-Cre mice crossed with either heterozygous or homozygous Ai32(RCL-ChR2(H134R)_EYFP) mice. Pvalb-IRES-Cre is expressed in the male germline. To avoid germline deletion of the stop codon in the loxP-STOP-loxP cassette, Pvalb-IRES-Cre;Ai32 mice were not used as breeders. Sst-IRES-Cre;Ai32 breeding sets (pairs and trios) consisted of heterozygous Sst-IRES-Cre mice crossed with either heterozygous or homozygous Ai32(RCL-ChR2(H134R)_EYFP) mice. Cre+ cells from Ai32 lines are highly photosensitive, owing to the expression of Channelrhodopsin-2. All experiments on animals were conducted with the approval of the Allen Institute’s Institutional Animal Care and Use Committee.

### Macaque monkey experiments at Vanderbilt University: Animals

All procedures were approved by the Vanderbilt University Institutional Animal Care and Use Committee in compliance with the regulations set by the Association for the Assessment and Accreditation of Laboratory Animal Care and followed the United States National Institutes of Health guidelines. Two macaque monkeys were used in this study: one male adult bonnet macaque (*Macaca radiata*) aged 19 years (Ca, 7.4 kg), and one male adult rhesus macaque (*Macaca mulatta*) aged 11 years (Jo, 12.0 kg).

Monkey Ca was injected with AAV vectors carrying an inhibitory cell-specific Dlx-ChR2-mCherry transgene. We mapped the planned trajectory of these injections with co-registered CT and MRI images. While the monkey was awake and head-fixed, we advanced an injectrode (32-channel Plexon V Probe customized with a fluid channel, Plexon, Dallas, US) connected to a Hamilton syringe (Hamilton Company, Reno, US) containing the viral vector in areas MT, MST, 8A, and PFC. Each injection site received 3 injections spaced 1 mm apart in depth, starting from the deepest location. Injections were followed by a one-minute waiting period, after which the injectrode was retracted to the next injection depth. We injected in total 57.6 μL of AAV1-Dlx-ChR2-mCherry and 14.4 μL of AAV9-Dlx-ChR2-mCherry (9.6 μL AAV1 and 2.4 μL AAV9 in PFC, 19.2 μL AAV1 and 4.8 μL AAV9 in 8A, 14.4 μL AAV1 and 3.6 μL AAV9 in MT, 14.4 μL AAV1 and 3.6 μL AAV9 in MST). Each injection was confirmed by simultaneous recording of neuronal activity on nearby channels near the fluid port, where neuronal activity decreased during injections.

### Mouse experiments at the Allen Institute: Surgery

The surgery pipeline at the Allen Institute is described in detail in previous reports^12,63^. Briefly, A pre-operative injection of dexamethasone (3.2 mg kg^−1^, subcutaneously was administered 1 hour before surgery to reduce swelling and postoperative pain by decreasing inflammation. Mice were initially anesthetized with 5% isoflurane (1–3 min) and placed in a stereotaxic frame (Model 1900, David Kopf Instruments, Tujunga, US). Isoflurane levels were maintained at 1.5–2.5% for the duration of the surgery. Body temperature was maintained at 37.5 °C. Carprofen was administered for pain management (5–10 mg/kg, subcutaneous), and atropine was administered to suppress bronchial secretions and regulate heart rhythm (0.02–0.05 mg/kg, subcutaneous). An incision was made to remove the skin, and the exposed skull was leveled with respect to pitch (bregma–lambda level), roll, and yaw. The headframe was placed on the skull and fixed with White C&B Metabond (Parkell, Edgewood, US). Once the Metabond was dry, the mouse was placed in a custom clamp to position the skull at a rotated angle of 20° to facilitate the creation of the craniotomy over the visual cortex. A circular piece of skull 5 mm in diameter was removed, and a durotomy was performed. The brain was covered by a 5-mm-diameter circular glass coverslip, with a 1-mm lip extending over the intact skull. The bottom of the coverslip was coated with a layer of polydimethylsiloxane (SYLGARD 184, Sigma-Aldrich, Burlington, US) to reduce adhesion to the brain surface. The coverslip was secured to the skull with Vetbond (Patterson Veterinary, Houston, US). Kwik-Cast (World Precision Instruments, Sarasota, US) was added around the coverslip to further seal the implant, and Metabond bridges between the coverslip and the headframe well were created to hold the Kwik-Cast in place. At the end of the procedure, but before recovery from anesthesia, the mouse was transferred to a photodocumentation station to capture a spatially registered image of the cranial window.

Intrinsic signal imaging was performed to identify visual area boundaries^64^. An insertion window was designed for each mouse based on the identified visual areas. Mice were habituated to head fixation and visual stimulation over two weeks.

### Macaque monkey experiments at Vanderbilt University: Surgery

Monkeys were implanted with a headpost and recording chambers during an anesthetized procedure. Monkeys were anesthetized with isoflurane. Body temperature was maintained near 38°C with a circulating water or warm air recirculating heating blanket. Peripheral oxygen levels, heart rate, temperature, ECG, and expired CO2 levels were monitored continuously during surgery. After mounting the animal in a stereotaxic device, the animal’s head was shaved and scrubbed (with Betadine and ethanol). Local anesthetic (Lidocaine) was injected subcutaneously before opening and placed into the ear canals before fitting the animal into the stereotaxic device.

A skin flap was made over the skull. The skull was cleaned and dried, and orthopedic screws were placed in the skull. The head restraint device, held in position with a stereotaxic holder, was fixed in place with orthopedic screws and methyl methacrylate cement. Custom-made recording chambers (~20 mm in diameter, Crist Instruments) were placed on the skull to allow access to all targeted brain areas in one hemisphere and secured in place with orthopedic screws and dental cement. Craniotomies (10-20 mm diameter) were subsequently performed using piezosurgery drills (Piezosurgery, Columbus, US) to permit access to the underlying cortex. Monkey Ca had two recording chambers, one placed over the prefrontal cortex (giving access to areas 8A and LPFC) and one placed over the temporal cortex (giving access to areas MT, MST, V4, and TEO). Monkey Jo had one recording chamber positioned over the occipital cortex (giving access to areas V1, V2, and V3). Both animals were used for electrophysiology experiments, only monkey Ca was used for optogenetics (details below). After surgery, analgesics were given for pain management (Buprenorphine and NSAIDs for 3 days), and animals showed no signs of pain or distress.

### Mouse experiments at the Allen Institute: Electrophysiology experiments

The Neuropixels data was acquired at the Allen Institute as part of the OpenScope project that allows the community to apply for observation on the Allen Brain Observatory platform (https://alleninstitute.org/division/mindscope/openscope/). The Neuropixels pipeline at the Allen Institute is described in detail in previous reports^12,63^.

On the day of recording, the cranial coverslip was removed and replaced with the insertion window containing holes aligned to six cortical visual areas. Mice were allowed to recover for 1–2 hours after the window was placed before being head-fixed in the recording rig.

Six Neuropixels probes were targeted to each of the six visual cortical areas (V1, LM, RL, AL, PM, AM). The boundaries of these areas and their retinotopic maps were obtained through intrinsic signal imaging, and insertion locations were planned to target regions responsive to the center of the LCD monitor. Probes were doused with CM-DiI (1 mM in ethanol; V22888, Thermo Fisher Scientific, Waltham, US) for post hoc ex vivo probe localization. Each probe was mounted on a 3-axis micromanipulator (New Scale Technologies, Victor, US). The tip of each probe was aligned to its associated opening in the insertion window using a coordinate transformation obtained via a previous calibration procedure. XYZ manipulator coordinates were obtained via a previous calibration procedure where the tips of probes were aligned to the retinotopic centers of target visual areas using an image provided by intrinsic signal imaging. The operator then moved each probe into place with a joystick, with the probes fully retracted along the insertion axis, approximately 2.5 mm above the brain surface. The probes were manually lowered to the brain surface until spikes were visible on the electrodes closest to the tip. After the probes penetrated the brain to a depth of around 100 μm, they were inserted automatically at a rate of 200 μm/min (total of 3.5 mm or less in the brain). After the probes reached their targets, they were allowed to settle for 5–10 min.

Neuropixels data was acquired at 30 kHz (spike band, 500 Hz high-pass filter) and 2.5 kHz (LFP band, 1,000 Hz low-pass filter) using the Open Ephys GUI^65^. Videos of the eye and body were acquired at 60 Hz. Pupil size was measured via previously described measures^12^. Briefly, a universal eye tracking model trained in DeepLabCut^66^, a ResNET-50-based network, recognizes up to 12 tracking points each around the perimeter of the eye, the pupil, and the corneal reflection. The angular velocity of the running wheel was recorded at the time of each stimulus frame, at approximately 60 Hz.

The spike-band data was median-subtracted, first within-channel to center the signal around zero, then across channels to remove common-mode noise. The median-subtracted data file was sent through the Kilosort2 MATLAB (The Mathworks, Natick, US) package (https://github.com/mouseland/Kilosort^67^, which applies a 150 Hz high-pass filter, followed by whitening in blocks of 32 channels. Kilosort2 models this filtered, whitened data as a sum of spike ‘templates’. The shape and locations of each template were iteratively refined until the data could be accurately reconstructed from a set of N templates at M spike times. Finally, Kilosort2 output was curated to remove putative double-counted spikes and units with artefactual waveforms. All units not classified as noise were packaged into Neurodata Without Borders (NWB) files^68^ and analyzed in this paper.

PV and SOM cells were identified in their respective mouse lines through a brief opto-tagging session at the end of the recordings^69^. Cells were stimulated with an optical fiber at 40 Hz, 5 Hz, and with a raised cosine function in 1 s epochs. Blue light was delivered by a 473 nm laser (Laser Quantum, Stockport, UK) with a measured light power at the tip of the fiber between 12.5-30 mW. The light source was coupled to a 400 µm diameter fiber optic cable (Thorlabs, Newton, US), with the tip positioned such that blue light illuminated the entire cranial window. Light was provided using three different types of stimuli, randomly interleaved: 2 ms pulses at 5 Hz, 40 Hz, or a 1s raised cosine ramp. Mice had no exposure to the optotagging stimulus prior to the end of the recording session. Cells whose mean response during the stimulation epoch positively exceeded 2 standard deviations of their baseline prestimulus (100 ms epoch) activity to any of the stimulation types were deemed to belong to the associated interneuron group.

Current source density (CSD^70,71^ was estimated from the LFP band data by epoching the LFPs, median-filtering with a filter-width of 3 channels, and then downsampling to every 4^th^ channel to retrieve a single vertical/laminar column of responses. CSD was then estimated as the 2^nd^ spatial derivative across multiple contact points according to the equation^72^:

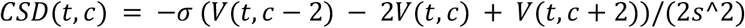

*V(t, c)* is the extracellular voltage at time *t* measured at an electrode contact at position *c, s* is the inter-contact distance of the electrode array, and σ is the tissue conductivity (here taken as 0.4 S/m). CSD contrasts were computed by grand-averaging the raw LFPs over each condition, then calculating the CSD as above and subtracting across conditions. We corrected for time in task as a confounder via double contrasts, first subtracting the 3^rd^ (fully adapted) presentation response from that of the 4^th^, and then pooling trials within task conditions. These subtractions were treated as normal random variates for a two-tailed, nonparametric spatiotemporal cluster-test that controls the family-wise error rate to *p* < 0.05 to determine significant cross-condition contrasts^36^. LFP and CSD are smoothed with a width 4 Gaussian filter in both the spatial (laminar) and temporal dimensions for plotting.

### Macaque monkey experiments at Vanderbilt University: Electrophysiology experiments

We acquired data using between 1 and 4 linear array multi-channel recording probes. In most experiments, we recorded with 128-channel “deep array” probes where the inter-contact spacing was 25 or 40 μm (Diagnostic Biochips, Glen Burnie, US) and the coverage was 3.175 mm and 5.08 mm, respectively. In some experiments, we used 32-channel V probes with an inter-contact spacing of 100 μm (Plexon, Dallas, US) with a coverage of 3.1 mm. We used a mechanical Microdrive (Narishige, Tokyo, JP) mounted onto the chambers to insert electrodes into the appropriate depth and area. A guide tube was lowered through a recording grid (Crist Instrument, Hagerstown, US) to penetrate the dura mater and granulation tissue. We then acutely introduced the recording probe into the respective area and waited ~1 hour for the signal to settle before starting the recordings. Recordings were acquired using an RHD System (Intan Technologies, Los Angeles, US) sampling at 30 kHz. Recordings were referenced to the guide tube, which remained in contact with tissue. Grid positions were pre-determined based on a pre-recording MRI scan with the recording chambers and grid in place, with water to mark the trajectories of each grid hole position. We advanced the electrode until there was visually responsive neuronal activity on most channels and until the local field potentials showed a distinct spectral signature of the cortical sheet, characterized by large amplitude alpha/beta (10-30 Hz) oscillations in deep channels and gamma (40-150 Hz) oscillations in superficial channels^73–77^. Offline, using MATLAB (The Mathworks, Natick, US), the data was filtered between 500-5000 Hz, full-wave rectified, low-pass filtered below 250 Hz, and downsampled to 1000 Hz. We refer to this signal as the MUAe signal, and it reflects the activity of nearby surrounding neurons^21,75,78–81^.

We allowed three to seven weeks after the optogenetic injections for viral expression before stimulating and recording. Monkey Ca was awake and head-fixed during opto-stimulation. A DBC 128-channel probe with a fiber (an “optrode”) attachment (0.2 mm in diameter with a numerical aperture of 0.37) at the end was connected to a laser with a wavelength of 473 nm and output power of 318 mW (before fiber). Laser power at the fiber tip was tested in a calibration run with a photodiode at 92.1 mW maximum (SM1-Threaded Mounted Photodiode, Thorlabs, Newton, US) in an Optical Power Monitor (Thorlabs, Newton, US). On recording days, the optrode was advanced into the injection site (in MT, MST, 8A, and PFC in separate sessions) with a metal guide tube puncturing the dura mater. Electrophysiological signals were recorded on the same optrode and a non-optrode neural probe simultaneously in another area. For analysis of the monkey optogenetics dataset, we used isolated single units and used the same spike sorting techniques that were applied to mouse data (Kilosort2).

Neurons were stimulated with a 5 Hz and 40 Hz raised cosine function at 3 V, 4 V, 5 V amplitudes, at maximum output power of 13.9 mW, 71.2 mW, and 92.1 mW, respectively. Instantaneous firing rate was estimated by smoothing binary spike trains, using a nonparametric EPSP filter (20 ms smoothing window). Mean spike waveforms, attributes (trough-peak duration, voltage distribution along the probe), and quality metrics (presence ratio, overall firing rate, signal-to-noise-ratio, and inter-spike interval violations) were manually checked for each Kilosort2 sorted optotagged unit, and noisy/artifactual units based on these attributes were removed prior to analysis. Each single unit’s firing rate during laser stimulation (2000 ms) was compared with baseline (1000 ms before onset and 1000 ms after offset). Trial-averaged mean firing rates for each laser frequency and power level were compared with baseline using one-tailed Wilcoxon signed-rank tests (α=0.05, right-tailed). Magnitude-squared coherence between the laser and each unit was computed over the full spectrum of laser stimulation frequencies. Coherence was defined as significant under two criteria: if coherence between spiking and the laser during the laser interval was higher than that between spiking and the laser during the pre- and post-baseline periods, and if coherence exceeded a critical value^82^. Units significant for increased firing rate and coherence were considered optotagged.

### Mouse experiments at the Allen Institute: Visual stimulation parameters

All Neuropixels recordings were according to the standardized Neuropixels visual coding pipeline. Visual stimuli were generated using custom PsychoPy scripts and displayed using an ASUS PA248Q LCD monitor with 1920 × 1200 pixels (55.7 cm wide, 60 Hz refresh rate). Stimuli were presented monocularly, and the monitor was positioned 15 cm from the right eye of the mouse and spanned 120 dva × 95 dva. Monitor placement was standardized across rigs so that the mouse’s right-eye gaze would typically be directed at the center of the monitor. Then, a sequence of drifting gratings appeared (full-screen, 4 Hz drift rate, 1 cycle/degree, 0.8 Michelson contrast; the angle of the drifting grating was 45 or 135 degrees from the vertical midline) for 500 ms and was replaced by a blank screen for 500 ms. This sequence was repeated 4 times per trial, for a total trial duration of 4500 ms (500 ms baseline + 4 x 1000 ms stimulus presentations). The inter-trial interval (ITI), the time between the end of the last P4 stimulus and the beginning of the next P1 stimulus, was 1.5 s. Every 100 sequences, a blank, black screen was presented for 10 s.

### Macaque monkey experiments at Vanderbilt University: Visual stimulation parameters

We used MonkeyLogic (developed and maintained at the National Institute of Mental Health) to control the behavioral task^83^. Visual stimuli were displayed using PROPixx Pro projectors (VPixx Technologies, Quebec, CA) with a resolution of 1920 x 1080 at a 120 Hz refresh rate. The projector screen was positioned 57 cm in front of the monkey’s eyes. Luminance calibration was performed using a Photo Research PR-650 on the projector screen, yielding a median value of 2.613 cd/m^2^.

The monkey was trained to fixate its eyes around a central fixation dot (radius of fixation window: 1.5 dva). Eye position and pupil diameter were monitored using an Eyelink 1000 (SR Research, Ottawa, CA) at 250 or 500 Hz. A task began with an isoluminant gray screen. Once the monkey maintained fixation for 500 ms, a sequence of drifting gratings appeared (12 visual degree radius, 2 Hz drift rate, 1 cycle/degree, 0.8 Michelson contrast; the angle of the drifting grating was 45 or 135 degrees from the vertical midline) for 500 ms, and was replaced by a blank screen with the fixation dot remaining for 500 ms. This sequence was repeated 4 times per trial, for a total trial duration of 4500 ms (500 ms pre-sequence fixation + 4 x 1000 ms stimulus presentations). After the offset of P4, there was an additional 500 ms of fixation, followed by reward, followed by a 1 s blank screen period, followed by 500 ms of fixation to start the next trial. Because monkeys self-initiated fixation behavior, this introduced jitter into the effective ITI length, which had a minimum of 2 s and lasted until the animals fixated on the central fixation dot. The monkey was rewarded with a small amount of juice for maintaining fixation throughout the trial.

### Mouse experiments at the Allen Institute: Habituation to predicted sequence

Mice were habituated over the course of 5 behavior-only sessions in which only the predicted sequence (either x-x-x-y or y-y-y-x) was presented. This resulted in 2000 habituation trials before recordings.

### Macaque monkey experiments at Vanderbilt University: Habituation to predicted sequence

Monkeys were habituated to a predicted sequence over the course of 3 behavior-only sessions in which only the predicted sequence (either x-x-x-y or y-y-y-x) was presented. This resulted in 2850 complete habituation trials before recording sessions. Subsequently, after a recording session, monkeys were again re-habituated to one of the predicted sequences before the next recording session.

### Mouse experiments at the Allen Institute: Data analysis

We recorded 29977 total putative single units, averaging 359 (+/-42) units per animal and 60 (+/-3) units per probe after quality control procedures. Specifically, we restricted the units for analysis, consistent with previous reports using the same methods for analysis^12^, to those that exhibit a visual response that exceeded 5 standard deviations of the baseline firing rate, minimum of 0.1 sp/s, inter-spike interval (ISI) violations below 0.33, a presence ratio above 0.95, an amplitude cutoff below 0.1, and were found to be in areas V1, LM, RL, AL, PM, or AM. This resulted in a sample of 5176 units from 15 of 16 mice. No units from 1 mouse passed the criteria due to a destabilizing event during the recording. Additionally, data from another mouse were omitted from the analysis due to issues with the visual stimulus timing. As a result, our final sample consisted of 4807 single units from 14 mice. Data were not recorded from all areas in all mice: V1, LM, RL, PM, and AM were recorded in 13 mice, and AL was recorded in 9 mice.

### Macaque monkey experiments at Vanderbilt University: Data analysis

We recorded a total of 6,592 sites with MUAe signals across 8 cortical areas across 19 recording sessions from two monkeys: 640 in PFC (monkey Ca), 416 in 8A (monkey Ca), 512 in MST (monkey Ca), 512 in MT (monkey Ca), 1056 in V4 (monkey Ca), 1536 in V3 (monkey Ca and Jo), 896 in V2 (monkey Jo), and 1024 in V1 (monkey Jo). These MUAe signals were extracted from broadband signals with a Butterworth filter of stopbands at 450 Hz and 5500 Hz and passbands at 500 Hz and 5000 Hz. For analysis, we restricted our analysis to sites that showed visually evoked activity, defined as the period from 50-250 ms post-stimulus being significantly greater than the pre-stimulus baseline (using a threshold of p<0.05, rank-sum test). The resulting population used for analysis consisted of 508 sites in PFC, 177 in 8A, 161 in MST, 252 in MT, 230 in V4, 569 in V3, 328 in V2, and 270 in V1.

### Macaque monkey experiments at Vanderbilt University: Laminar identification

For each penetration into monkey recordings, we examined its trajectory using MRI and postmortem Nissl-stained images. Penetrations that are not sufficiently perpendicular to cortical layers or did not have enough cortical coverage were excluded from layer-based analyses. We then examined the multi-unit activity, CSD, and spectrolaminar pattern^74^ across depth for each penetration. Layer 4 of each area was identified based on the best corresponding electrode location with the earliest MUA onset, earliest CSD sink, and spectrolaminar high/low band relative power crossing. Total area thickness was measured in MRI images of each monkey for each area. We also obtained percent thickness per layer for each area from existing data^84^. The number of electrodes per layer was then calculated by multiplying the total layer thickness by the layer thickness for each area. We then assigned the layer position of each electrode by counting from the previously identified layer 4 electrode up to layer 1 and down to layer 6.

### Granger causality analysis

In mice, we began by defining all available single units in each area. We included all cortical neurons in this analysis; for each mouse and probe, we included an area if at least 12 single neurons were simultaneously acquired. This resulted in six neuronal spiking time series, one per area. Granger causality was calculated separately per mouse and for local and global oddball trials, using all available trials. We calculated Granger causality using a non-parametric implementation^41,42^. For this, we computed Fourier coefficients using the Fieldtrip toolbox^85^, with a sliding-window approach with a 200 ms window. We computed the Fourier coefficients using multitapers (3 multitapers per trial, time window size = 200 ms, smoothing = 10 Hz, function ‘ft_freqanalysis’, using method ‘mtmfft’, frequencies 0-100 Hz were calculated). One time window was on the baseline, from 100 ms pre- and 100 ms post-oddball onset. Two time windows reflecting stimulus processing were also used, one centered from 100 ms to 300 ms post-oddball onset, another from 300 ms to 500 ms post-oddball onset. Bivariate Granger causality was computed pairwise between all available neuronal spiking time series (using ft_connectivity analysis with method = ‘granger’ and granger.sfmethod = ‘bivariate’).

### Statistics and visualization

In mice, spiking activity for each cell was normalized to its average activity level to obtain an arbitrary value that was less variable across units with differing firing rates. In monkeys, spiking activity for each channel was normalized to each channel’s standard error across trials. Unless otherwise noted, all statistics were performed using nonparametric, cluster-based permutation testing^36^ of the spiking activity at either the single-(multi)unit level or the population level, with alpha thresholds set to 0.05 for both test stages. We also required that a statistically significant difference persist for at least 25 ms as indicated by this test. Data were smoothed using a 50-ms moving-mean window. Confidence bands, where displayed, are computed for a 95% confidence interval across units’ condition-wise mean activity included in the population.

In the main contrast, the third stimulus (P3) was subtracted from the fourth (P4) stimulus to control for time in session. The oddball stimulus was compared to the control stimulus. Therefore, the main contrast follows this equation:

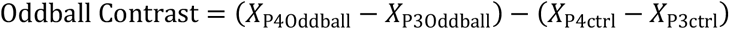

X in the above equation was a normalized quantity, with different definitions for the two datasets:

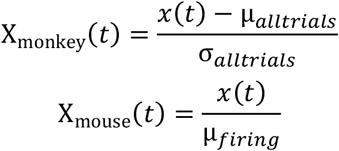

Where σ is the standard deviation and μ is the mean.

Feature selectivity was measured by first determining the mean response to each stimulus during visual stimulation (0-500 ms following stimulus onset) in a separate presentation block with randomized stimulus order. For each unit, we subtracted the mean response to the LO-orientation stimulus from the GO-orientation stimulus and divided it by the sum of those two means, defining the following selectivity index:

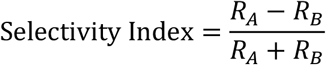

Such that R is the average neural response during the 500 ms of stimulus to A or B in the random control block in which A and B are presented randomly each with a 50% probability. The distribution of selectivity indices from each area was then examined and statistically tested for balance. For areas that showed a significant shift in median from zero on a one-sample Wilcoxon signed-rank test, stratification was performed to systematically remove extreme values until the distribution’s median was no longer significantly shifted from zero. The main contrasts were then performed again with the balanced distribution where indicated. In mice, single units with this index above 0.8 or below −0.8 were defined as highly selective for the GO orientation or the LO orientation, respectively.

For identifying the onset latencies of significance, a t-test was run at each time point during 500 ms of stimulus presentation for individual unit/channel (with the same selection criteria described above, only including unit/channel with significant visually evoked activity), and the first contiguous significant cluster larger than 10 ms was considered the onset of significance for that channel.

### Multi-model local large language model evaluation of Predictive Processing literature

The full methodology is described in Nejat et al., 2026^47^. Briefly, the three-dimensional hypothesis-space analysis was adapted from our glossary-constrained (Supplementary Tables 5-7) multi-LLM scoring pipeline for predictive processing literature evaluation. The glossary consisted of thirty-six factors organized into three hypotheses: predictive suppression (H1), feedforward error propagation (H2), and ubiquity (H3). This glossary defined the conceptual target space against which each study was evaluated. We considered a set of 25 studies (Supplementary Table 1). Each paper was scored separately in the local-oddball and global-oddball contexts (defined in Supplementary Table 4), allowing the same theoretical factors to be compared across both contexts. This design was important because it allowed us to ask whether evidence supporting predictive coding in local sensory deviance paradigms generalized to global, sequence-level prediction contexts. Rather than relying on unconstrained narrative summaries, the glossary forced each model to evaluate explicit hypothesis-relevant evidence using fixed definitions and a shared scoring scale.

Each study was processed through a deterministic local ingestion pipeline (see below). Full-text information was extracted from each source paper, and figure-based evidence was separately processed using a local vision-language model so that graphical results, statistical trends, and figure annotations could be included in the model-readable study representation. Text and figure descriptions were then merged into a unified markdown document for each paper. A constrained prompt (see *Data and code availability* section for full details) instructed each model to act as a neuroscientist and to score every glossary factor on a continuous scale from −1 to +1, where positive values indicated support for the factor, negative values indicated contradiction, zero indicated explicitly neutral evidence, and null indicated that the paper did not address the factor. For every non-null score, the model was required to provide an auditable reasoning log tied to textual, statistical, or figure-based evidence. This instruction layer was designed to reduce hallucinated interpretation and to make every score traceable to the source material.

The model council consisted of ten locally executed models (Supplementary Table 3). Models were run locally using the MLX-LM framework on Apple Silicon hardware^86^ (Apple, Cupertino, US), with matched inference settings across models. Critically, model context was cleared between every evaluation so that each paper-model run was independent and could not inherit information from a previous run. Outputs were validated for correct glossary keys, valid score formatting, and required reasoning fields before being entered into the final score table. The validated scores were then averaged across models to place each study into a three-dimensional hypothesis space defined by predictive suppression, feedforward error propagation, and ubiquity, separately for local and global oddball contexts. Variability was defined across models using standard error of the mean.

## Data and code availability

All mouse data used in this work are openly available through the DANDI Archive (dandiset id: 000253, doi: https://doi.org/10.48324/dandi.000253/0.240503.0152 for the mouse data and dandiset id: 001838, doi: https://doi.org/10.48324/dandi.001838/0.260616.2029). Software resources for the analysis of this dataset are provided here: https://alleninstitute.github.io/openscope_databook/projects/glo.html and here: https://github.com/yihan777/GlobalLocalOddball. The code to generate the global-local oddball sequences is provided here: https://github.com/AllenInstitute/openscope-glo-stim. The code and resources for the multi-LLM evaluation pipeline is provided here: https://github.com/HNXJ/mllm-public.

**Extended Data Fig. 1.**
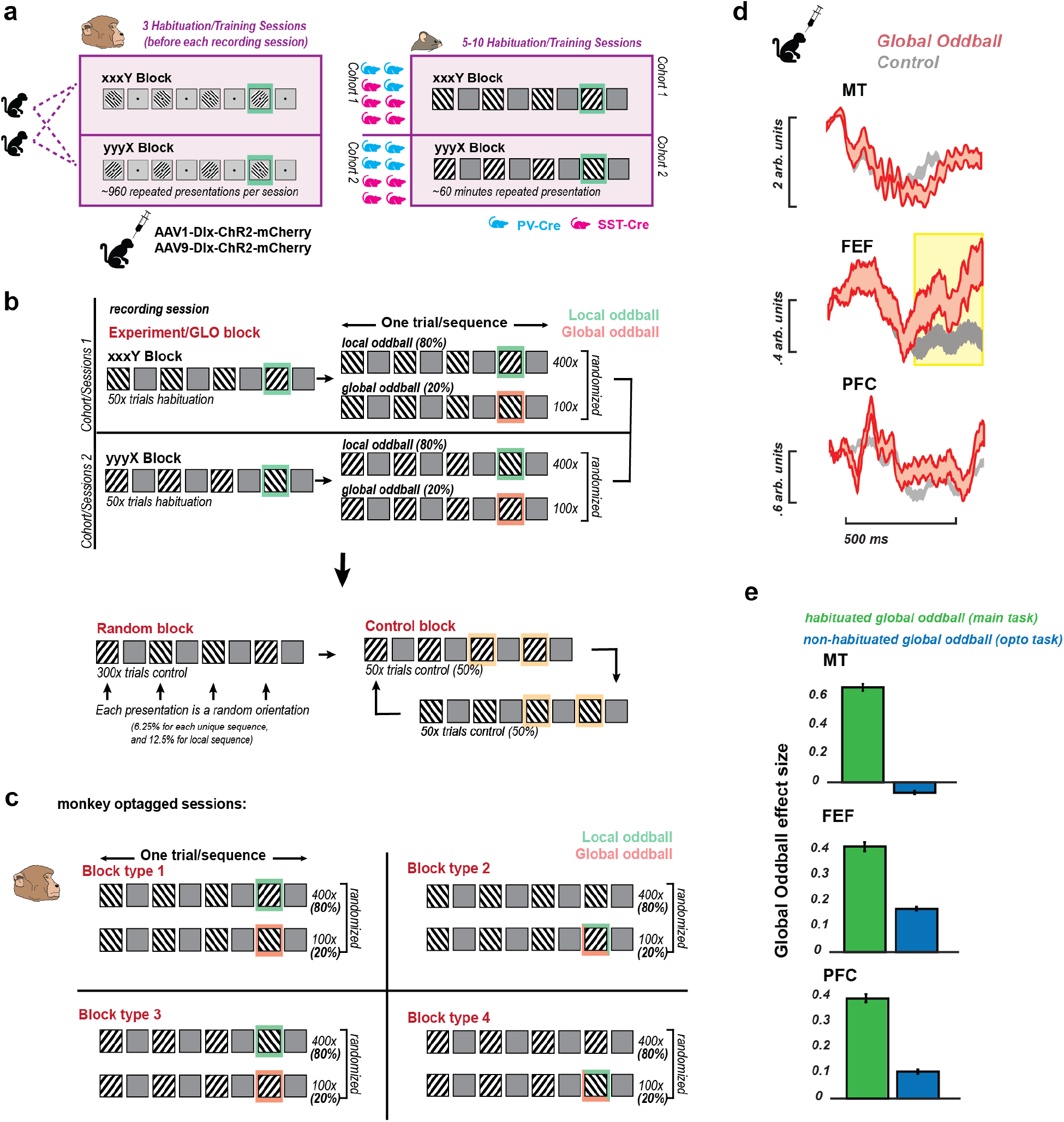
Experimental Design and task responses without habituation. **a**, 2 macaque monkeys were habituated to the local oddball pattern (x-x-x-y or y-y-y-y on a given recording week) for ~960 repeated presentations for 3 days prior to recording day. 16 transgenic mice were used in the experiment; genotypes were selected to allow for optotagging of either somatostatin-(magenta) or parvalbumin-positive (cyan) interneurons. Mice were split into 2 training groups: 1 group exposed to x-x-x-y sequences during the entirety of training and 1 group to y-y-y-x. **b**, Recording sessions were divided into 3 blocks. In the experiment (GLO) block, animals experienced their trained ‘Local’ oddball sequence (x-x-x-y or y-y-y-x) for a 50-trial habituation period. Then, on 20% of the next ~500 sequences, they observed the ‘Global’ oddball sequence (x-x-x-x or y-y-y-y). A block of random sequences of 4 stimuli followed the experiment block (resulting in a 6.25% probability for each unique sequence, with the sequence identical to the local oddball sequence upsampled to 12.5%). A control block followed where sequences of x-x-x-x (50%) and y-y-y-y (50%) were experienced. **c**, Opto-tagging sessions in monkeys were performed with a classical local/global oddball paradigm to ensure cell stability between oddball and control trials. **d**, Pure global oddball response in optogenetics experiment without habituation (paradigm shown in sub-panel **c**). MUA response to global oddball/local regular (P4-P3) in red and global regular/local regular (P4-P3) in grey. **e**, Global oddball effect size from the optogenetics experiment (local global oddball experiment without previous habituation, shown in sub-panel **c** compared to main experiment (~3 habituation sessions prior to recording, shown in sub-panels **a** and **b**.

**Extended Data Fig. 2.**
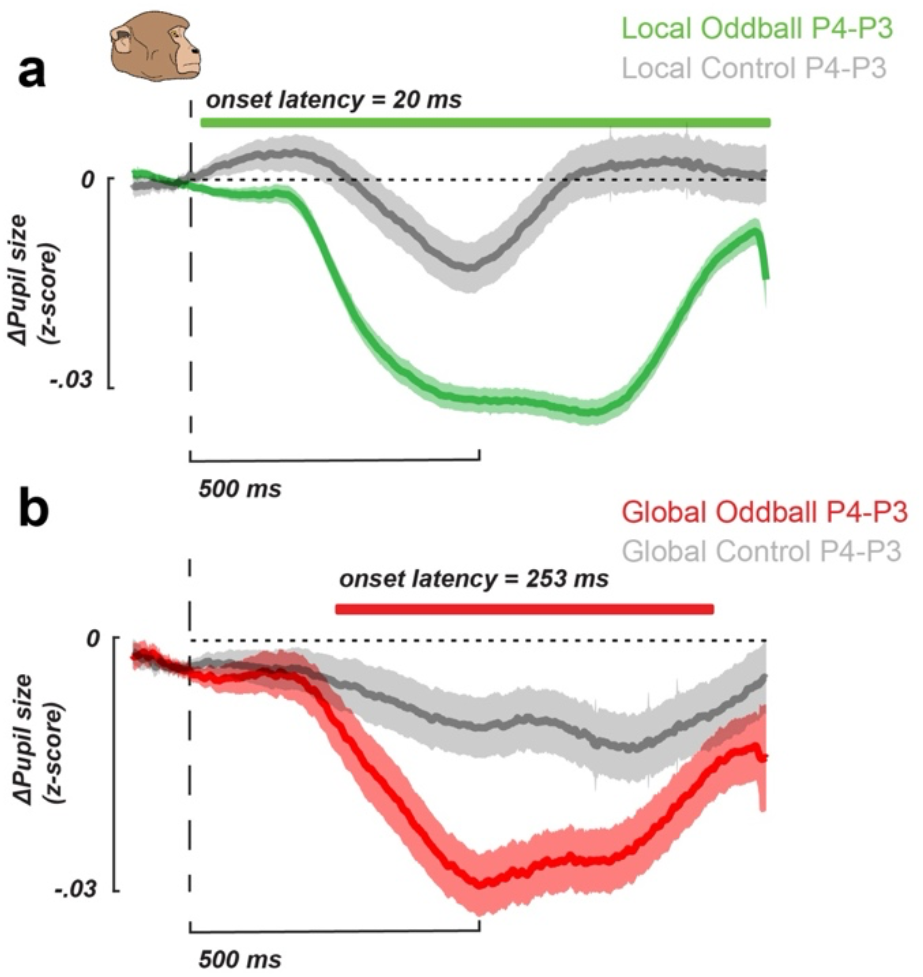
Global oddball detection indicated by pupillary response in monkeys. **a**, Average z-scored change in pupil size between P4 (oddball) vs. P3 (stimulus prior to oddball) during a local oddball sequence in green, and during a local control sequence in grey. The vertical dotted line shows stimulus onset. The horizontal green bar indicates the period during which the traces differ significantly. The onset time of this cluster was 20ms after the onset of the local oddball. First-level statistics were a t-test at alpha = 0.05, and the second-level statistic for the cluster-based permutation test was alpha = 0.05. **b**, Average z-scored change in pupil size between P4 (oddball) vs. P3 (stimulus prior to oddball) during a global oddball sequence in red, and during a global control sequence in grey. Colored bands represent 95% confidence interval. Vertical dotted line shows stimulus onset. The horizontal red bar indicates the period during which the traces differ significantly. The onset time of this cluster was 253ms after the onset of the global oddball. First-level statistics were a t-test at alpha = 0.05, and the second-level statistic for the cluster-based permutation test was alpha = 0.05.

**Extended Data Fig. 3.**
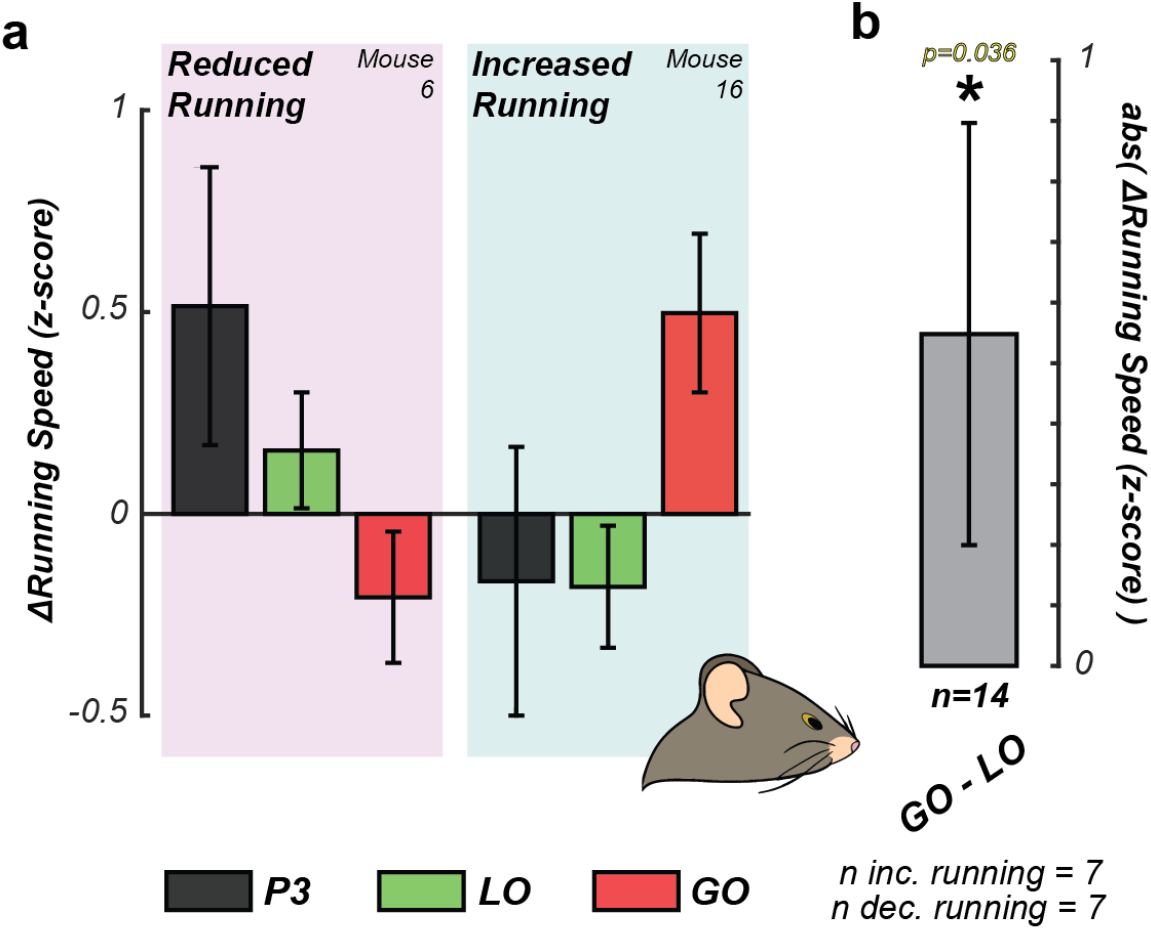
Changes in mouse running behavior indicate recognition of unpredicted stimulus. **a**, Average running behavior as change from prestimulus baseline (500 ms before stimulus onset) from two example mice (mouse 6, left; mouse 16, right) during the 1 s following stimulus onset for P3 (black), LO (green), and GO (red) during the main block. Error bars indicate +/-2 SEM. Some mice increased their running behavior in response to the GO (unpredictable) stimulus (n=7 mice) compared to predictable stimuli (P3, LO), and others decreased their running behavior (n=7 mice). **b**, Population-level quantification of change in running behavior in response to unpredictable stimulus. The average change in running behavior for the GO was contrasted with the LO (GO-LO), and the absolute value was taken for each mouse. Those mouse-level values were then averaged to get the population average shown as the gray bar. Error bars indicate +/-2 SEM. To determine whether there was a population-level significant change in running behavior as a function of stimulus predictability, a bootstrapped estimate contrast with stimulus-label-shuffled data was computed. More specifically, we generated a shuffle “GO” distribution and a shuffle “LO” distribution for each mouse that shared a label-shuffled pool of running measures from all GO/LO presentations in the main block. The number of samples taken for bootstrapping was done to match the original number of trials for the GO (~100 trials) and LO (~400 trials) distributions. The shuffle LO mean of that distribution was subtracted from the shuffle GO and the absolute value taken. This was then redone for 10000 bootstrap resamples to determine the shift from 0 per mouse in the absence of stimulus predictability information. That per-mouse estimate was then subtracted from the original per-mouse GO-LO (non-shuffled) contrast, thereby accounting for the 0-limit bias brought in via the absolute-value component. This additional step meant that, at the population level, we could determine whether there was a significant difference in the running behavior as a function of the stimulus predictability without biasing the result with the absolute value operation. We then performed a one-sample t-test across mice (n=14), yielding P=0.036, indicating a change in running behavior across the population as a function of stimulus predictability.

**Extended Data Fig. 4.**
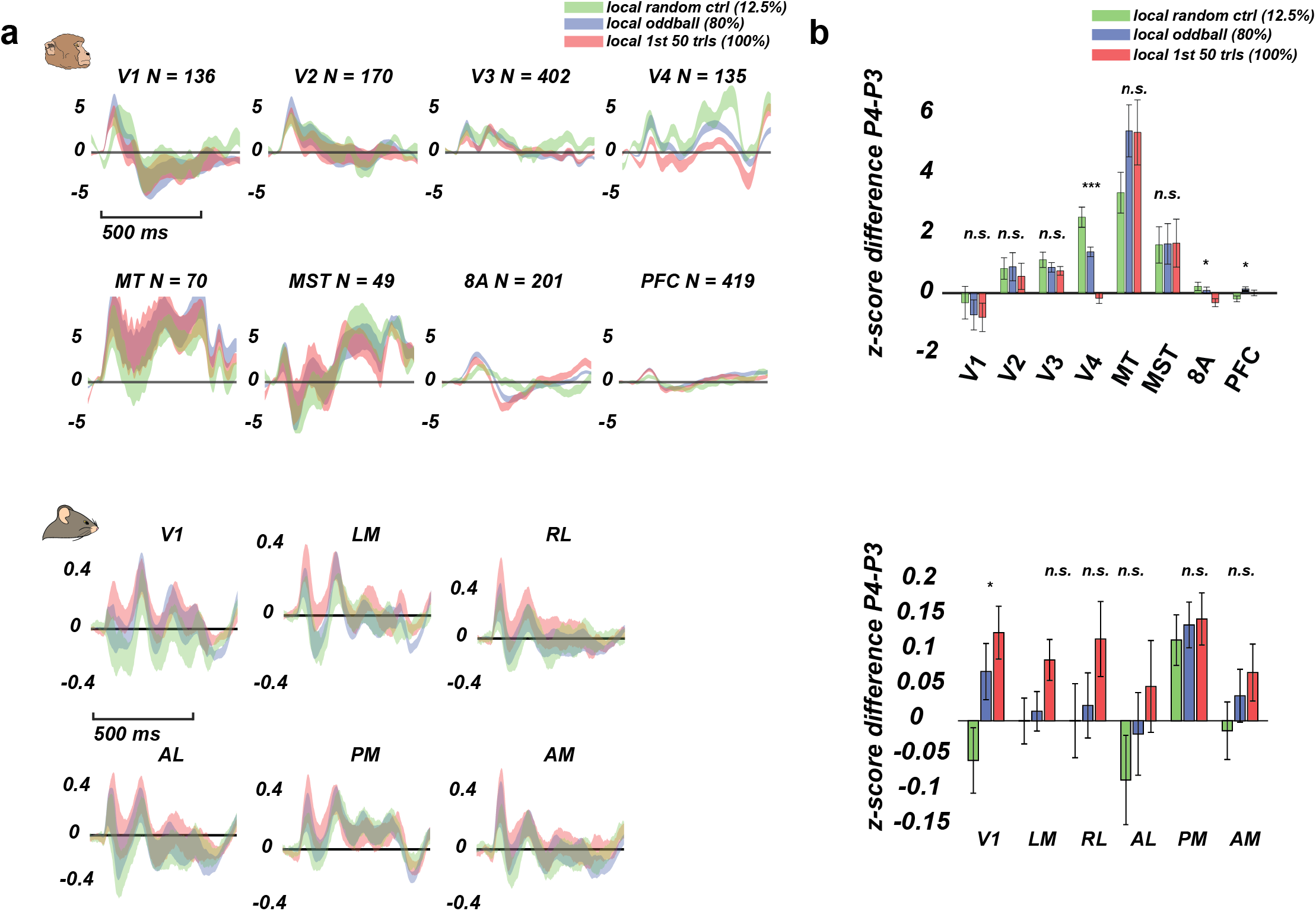
Lack of prediction effect in local oddball sequence of various probabilities. **a**, Z-score multiunit activity in monkeys (top) and mice (bottom) in response to local random control (12.5%, green), local oddball (80%, blue), and first 50 trials of local oddball (100%, red) across cortical hierarchy. Shadings show 2 SEM across units/channels. **b**, Z-score difference between P4 and P3 (effectively representing effect size) of local random control, local oddball, and first 50 trials of local oddball in monkeys (top) and mice (bottom). Single asterisk represents P<0.05 from a one-way ANOVA test. Triple asterisks represent P<0.001 from a one-way ANOVA test.

**Extended Data Fig. 5.**
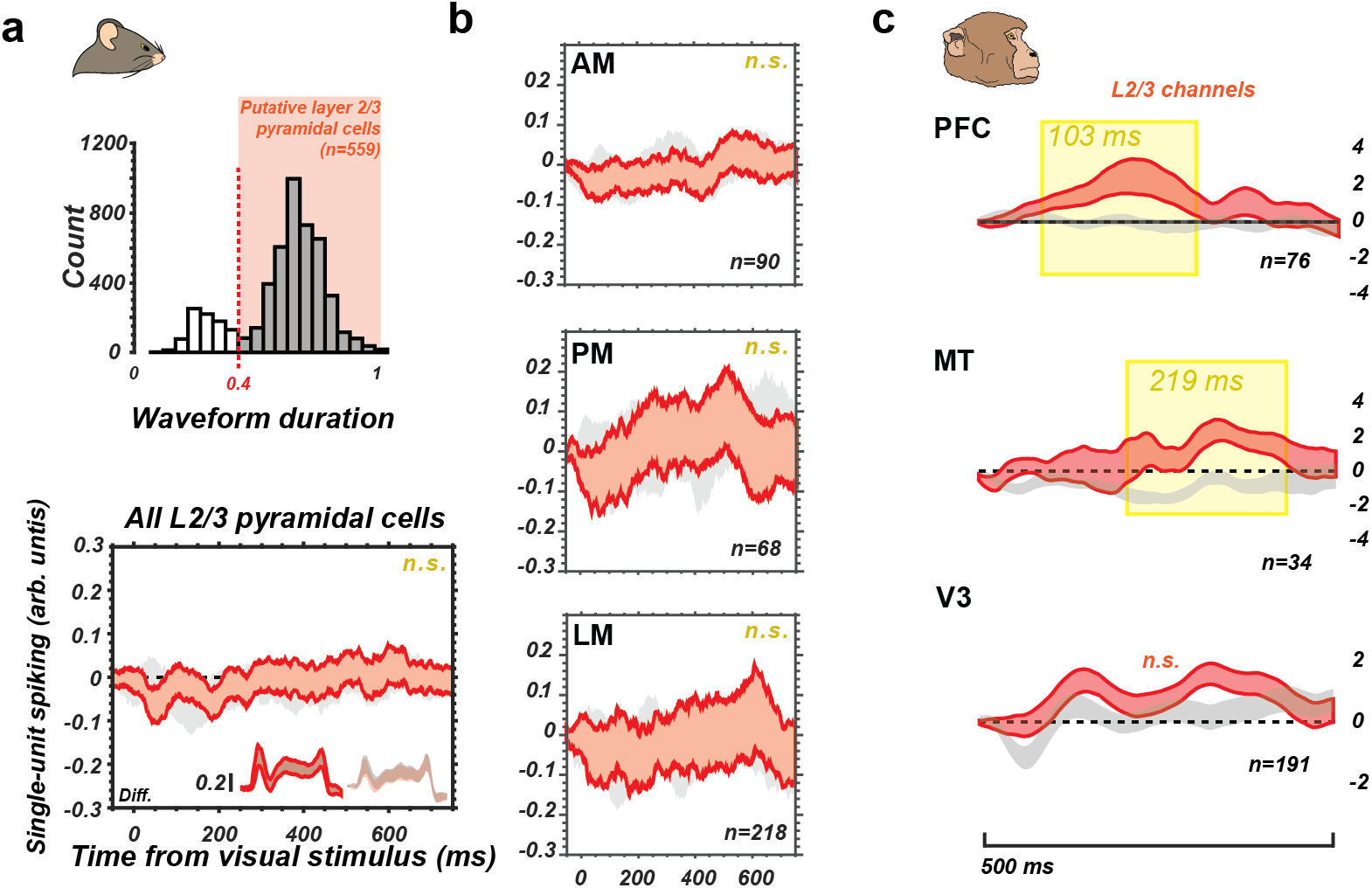
Putative layer 2/3 neurons do not harbor latent feedforward global oddball signals. **a**, Global oddball response in putative layer 2/3 pyramidal cells in mice. To identify layer 2/3 pyramidal units in mice, we eliminated single units that were found to be inhibitory interneurons via optotagging, we eliminated units found outside layer 2 or 3, and we ensured units that had a spike waveform duration of longer than 0.4 ms (waveform duration histogram is shown in upper subplot), which primarily reflects broad-spiking pyramidal cells. (lower subplot) Red bands are P4-P3 in the main block; gray bands are P4-P3 in the control block. Insets show raw traces of P4 and P3 presentation responses. **b**, same analysis as in **a**, but separately per area in mice that showed a population-level global oddball response. **c**, Global oddball responses in L2/3 in monkeys in areas that had population-significant global oddball responses (V3, MT, PFC) and in which laminar recordings were possible. Red bands are P4-P3 in the main block for global oddball trials compared to the control block (in gray), time-locked to oddball onset. Mean +/-2 SEM (the n in each sub-panel indicates number of L2/3 MUAs per area). Shading indicates significant clusters over time. Numbers within shaded region indicate the onset of the cluster in ms.

**Extended Data Fig. 6.**
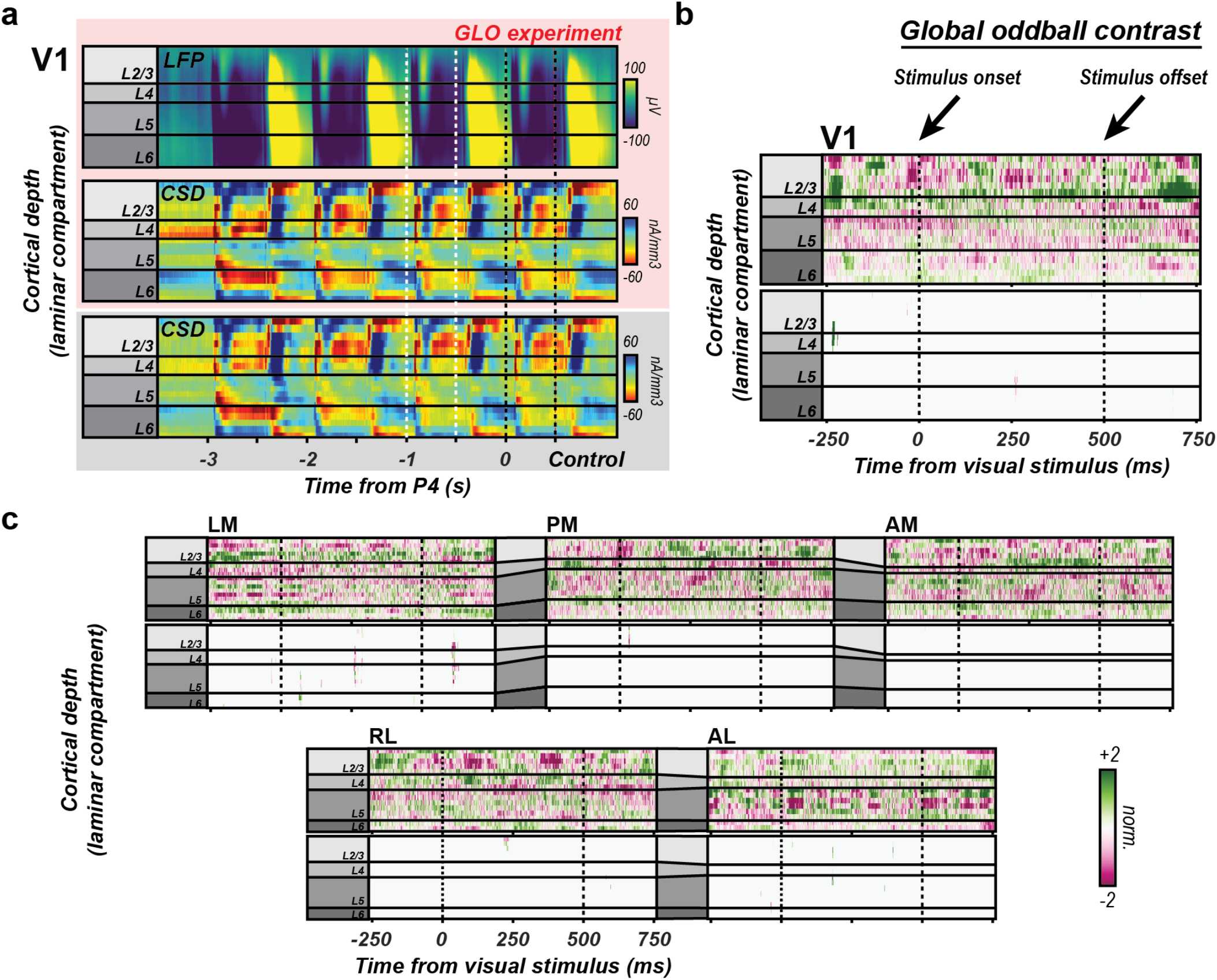
CSD reflecting synaptic activity does not indicate top-down predictions being fed back to the early sensory cortex. **a**, Laminar CSD is calculated from the LFP response (top) to sequence presentations averaged across trials and mice (n=9) for area V1 after aligning cortical depth between animals. CSD is calculated for the global oddball experimental condition (middle, red-highlighted region) and the control condition (bottom, gray-highlighted region). The epoch between the white dotted lines indicates presentation 3, and between the black dotted lines indicates the global oddball presentation (middle) or presentation 4 in the control (bottom). **b**, Evaluating the global oddball contrast for evidence of differences in synaptic activity with global oddball detection in V1. Difference of differences identical to previous global oddball contrasts (P4-P3) for the CSD data. The top image shows all values after contrast and normalization that fall between −2 and +2 standard deviations. The bottom image shows the same data with a mask applied to display only significant values (Welch’s t-test, p<0.01). Color map intensity is indicated by the scale bar at the bottom right of the figure. **c**, Difference of differences for global oddball contrast for the remaining visual cortical areas. Formatting identical to b.

**Extended Data Fig. 7.**
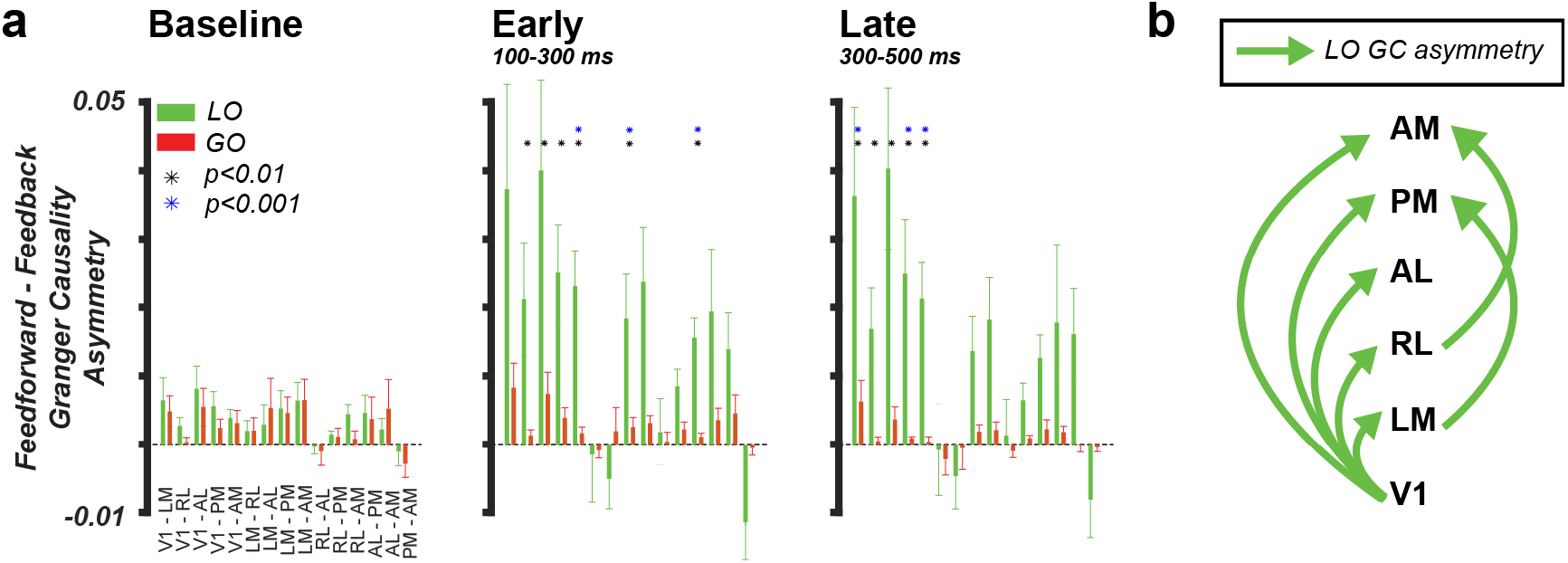
Cortical spiking causalogram for local and global oddballs: Granger causality as a function of time across brain areas. **a**, Granger causality asymmetry (Feedforward minus feedback GC) between all possible areas in the baseline (left, pre-oddball time window; middle, initial sensory processing period; right, late sensory processing period). Green bars denote local oddballs, and red bars denote global oddballs. Mean across mice, +/-SEM across mice. Asterisks indicate connections that are significantly different from the baseline at P<0.01 (black asterisks) and at P<0.001 (blue asterisks). **b**, Hierarchical summary of significant changes in GC asymmetry from pre-oddball baseline for local oddballs. Significant interactions at P<0.01 and P<0.001 are shown, and both early and late periods are depicted. Global oddballs are not shown because no significant changes in GC asymmetry were observed.

**Extended Data Fig. 8.**
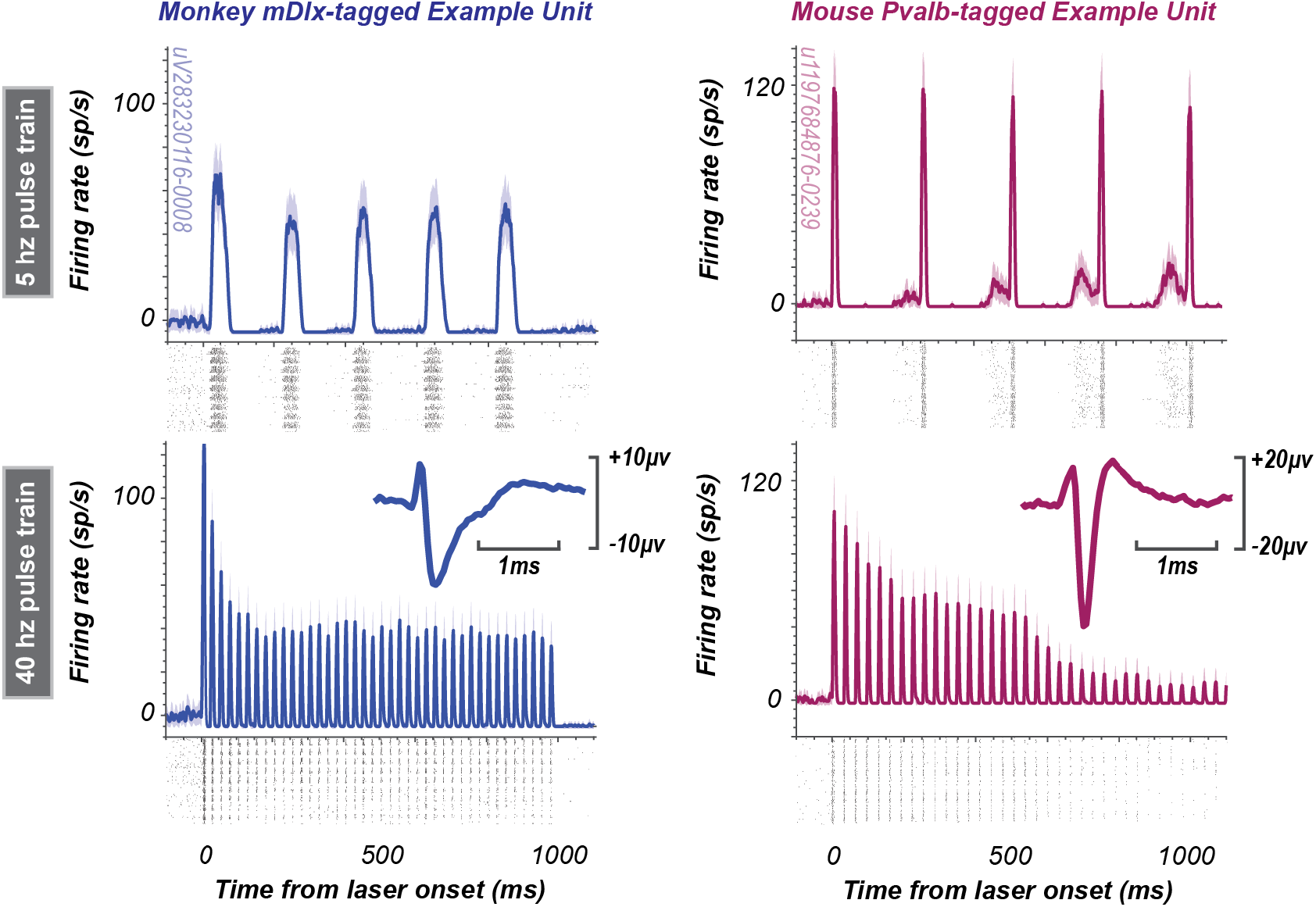
Optogenetic identification of single units. Example units with optogenetic modulation to 5 Hz (upper panels) and 40 Hz (lower panels) in monkeys (left panels, targeting pan-inhibitory interneurons) and in mice (right panels, in this example, targeting a Parvalbumin-positive inhibitory interneuron). Traces indicate average smoothed firing rates (spikes per second) across trials (+/-SEM). Rasters indicate individual spikes, stacked across trials. Insets indicate the average waveform shape of the identified unit.

**Extended Data Fig. 9.**
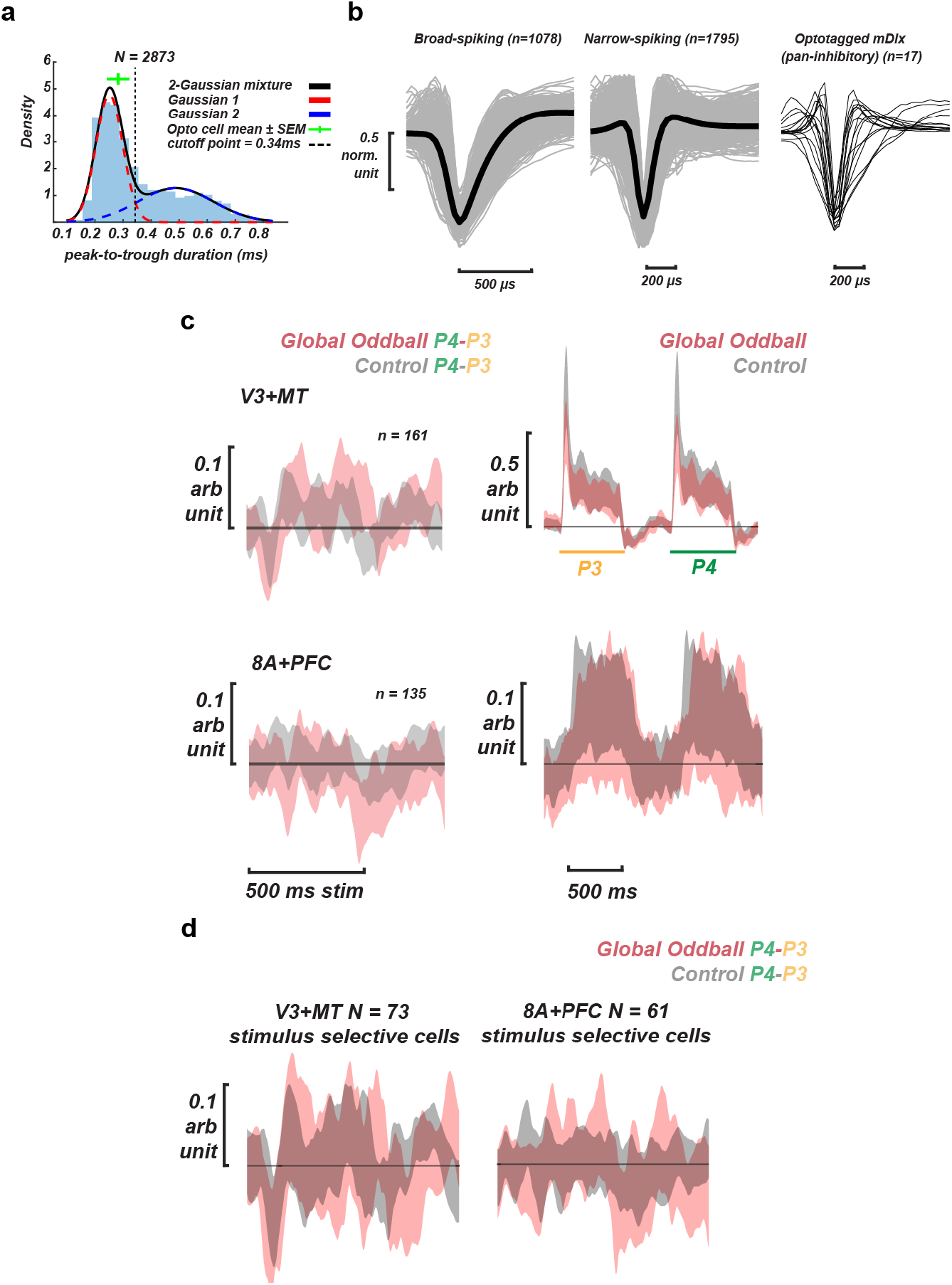
Putative Inhibitory interneuron identification by waveform shape in monkeys. Putative inhibitory cells in monkeys do not significantly decrease or increase in response to global oddball compared to the control condition. **a-c**, replication of Fig. 4e,f for reference. **a**, The distribution of peak-to-trough duration of all cells from the main experiment and the optogenetics experiment, and the optotagged cells. A two-Gaussian mixture fit yielded a cutoff of 0.34 ms, which was used to identify putative inhibitory cells. **b**, All normalized waveforms and waveform average for putative inhibitory and excitatory cells, and normalized waveforms for all optotagged cells. **c**, Top panel: neuronal response to global oddball (P4-P3) in red compared to control condition (P4-P3) in gray. Middle panel: neuronal stimulus response to global oddball, P4 in red and P3 in gray. Bottom panel: neuronal stimulus response to global control condition, P4 in red and P3 in gray. **d**, The neuronal response to global oddball (P4-P3) in red compared to control condition (P4-P3) in gray in only GO-stimulus-selective neurons.

**Extended Data Fig. 10.**
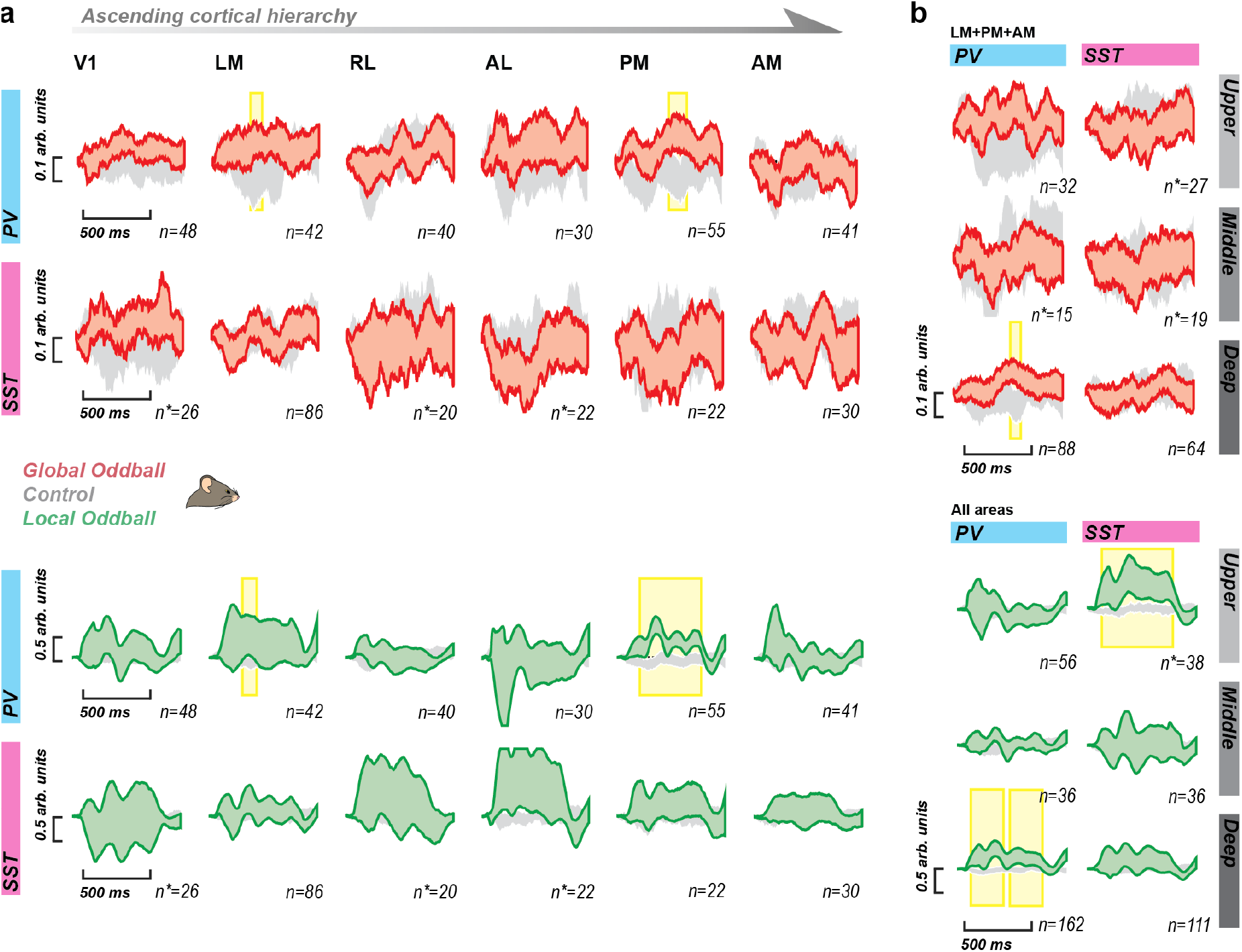
PV+ and SST+ neuron activity by area and layer in mice. **a**, GO contrast (P4-P3, GO sequence in main block, red bands) vs. control contrast (P4-P3; GO stimulus sequence in the control block, gray bands) (top) and LO contrast (P4-P3, LO sequence in main block, green bands) vs. control contrast (P4-P3, LO stimulus sequence in control block, gray bands) (bottom), averaged across units for each inhibitory cell type (PV and SST) across areas from all mice (n=14 mice). Bands are 95% confidence intervals across units in an area. Yellow highlights reflect significant population local oddball detection periods, P<0.05, corrected using nonparametric, cluster-based permutation tests. Number of units in each grouping, n, is indicated in the figure. **b**, identical to a, but for specific laminar compartments averaged across population-level global oddball-detecting areas (LM, PM, AM) (top) and population-level local oddball-detecting areas (i.e., all areas) (bottom). Specific data stratifications were susceptible to a population-level confound, where the population was imbalanced in the number of units that preferred one stimulus over the other. This was addressed by eliminating units from the analysis until the balance of units selective for one stimulus vs. the other was negligible. These are indicated for specific populations, with the sample size denoted by n* (the number of units after stratification). Typically, eliminating 1 unit was necessary to achieve balance.

## Supplementary information

### This file includes

Supplementary Text

Supplementary References

Supplementary Tables 1-7

Supplementary Figs. 1-11

## Supplementary Text

### Neuronal Selectivity analysis for local and global oddball processing (in mice)

The increased response observed during local oddballs (Fig. 2) cannot be explained by the fact that the oddball is a different stimulus (compared to the rest of the stimulus sequence) since we averaged across all neurons (thus eliminating any tuning-based preferences for one or the other stimulus) and about half of the animals saw the orthogonal orientation as the oddball stimulus. Moreover, across all recorded single units in the dataset, we found a highly similar number of units strongly preferring each of the two orientations (LO-orientation: 21.2% of all single units, GO-orientation: 22.3% of all single units). This indicates that there was not a strong response bias for one oddball feature over the other within the population of measured units. To summarize, even after completely randomizing neuronal biases, the local oddball stimulus consistently evoked a larger response when we examined spiking activity across layers, areas, and animals.

Previous reports have suggested that sensory prediction error signaling might be isolated to neuronal populations that are selective for the features that define the error-producing stimulus^87^. To test for this possibility, we isolated the population of GO-orientation selective neurons within our recorded populations and determined whether they specifically harbored a global oddball detection signal. Of the 22.3% of single units strongly selective for the GO orientation (relative to the LO orientation), only 6.1% were also significant GO detectors. This is comparable to the rate of GO detectors we observed in the general population (6.4% of single units). In other words, GO detection was not isolated from featuring selective populations, nor were all units highly selective for the GO stimulus found to be GO detectors.

### Additional details on Granger causality analysis

We calculated Granger causality (GC, see *Methods*) to quantify directed functional connectivity amongst neurons and address the directed propagation of oddball responses throughout the hierarchy. For each cortical area and mouse, we combined all available single neurons to create a population spiking time series (a minimum of n=12 neurons per area were used to create this population spiking time series). We then calculated GC between each pair of cortical regions, using the anatomical hierarchical ordering: V1, LM, RL, AL, PM, AM (in order from the earliest to the latest area in the mouse visual hierarchy^40^. This generated 30 inter-areal GC values, one for each area pair and direction of influence. We computed the GC hierarchical asymmetry for each pair of areas by subtracting the feedback GC from the feedforward GC. For example, the feedforward GC from V1 to RL was subtracted from the feedback GC from RL to V1. A positive difference indicates that feedforward GC > feedback GC. A positive difference for global oddballs would be consistent with the hypothesized (H2, Fig. 1a) feedforward propagation of prediction error. We compared oddball GC responses during stimulus processing in both the early (100-300 ms post-oddball onset) and late (300-500 ms post-oddball onset) time windows between local and global oddballs. We compared them to baseline (100 ms pre- and 100 ms post-oddball onset). GC became significantly hierarchically asymmetric in the feedforward direction for local, but not global, oddballs during both early and late periods (Extended Data Fig. 7, Wilcoxon rank sum test of GC hierarchical asymmetry values across mice vs. pre-oddball baseline, P<0.01). In contrast, GC hierarchical asymmetry did not change during global oddballs compared to baseline (Extended Data Fig. 7, Wilcoxon rank sum test of GC hierarchical asymmetry values across mice vs. pre-oddball baseline, all comparisons, P>0.01).

### Current source density analysis

To determine whether top-down prediction signatures might be latent in synaptic dynamics rather than spiking activity, we examined the Current Source Density (CSD, a measure of local net depolarization) profiles during global oddball detection in mice. CSD is frequently used as a proxy for synaptic activity occurring along the cortical column^77,81,88–95^, which reliably reflects columnar processing dynamics, including those that may be missed by spiking activity^75,76,96^. We computed CSD from the local field potential responses to the GLO and control sequences (Extended Data Fig. 6) and then performed the same global oddball contrast as with the spiking data. We hypothesized that a difference in current sinks in the upper or deep layers of the cortex might manifest during prediction errors (i.e., when the global oddball stimulus unpredicted in the main experimental block as compared to the control block). During global oddballs, we did not observe reliable changes in synaptic activation at the earliest stage of visual cortical processing (i.e., in V1, Extended Data Fig. 6b), neither before nor after visual stimulation. Specifically, we did not observe significant, reliable synaptic changes in the extragranular compartments, which would indicate top-down influences in deploying a prediction to earlier areas. We observed the same in downstream visual cortical regions (Extended Data Fig. 6c). This shows the predictions are not being deployed widely through synaptic modulations.

### Verifying statistical approach accounting for animal variability

For the bulk of statistical analyses, we used a cluster-based permutation test, which is reliable for detecting significant differences in electrophysiological data^36^. However, we treat all neurons as independent samples without considering the animal in which they were recorded. To verify that our results are not contaminated by animal-to-animal variability, we used a nested model design to reevaluate a central result of our study: that three brain areas in the mouse (LM, PM, and AM) detected global oddballs at the population level.

For each brain area separately, we tested whether the per-unit response differed between the GO (P4-P3 GO sequence in main block, contrast 1, C1) and Control (P4-P3 GO stimulus sequence in control block, contrast 2, C2) contrasts, accounting for animal-level variability using a two-level linear mixed-effects model in which units were nested within mice.

For unit *i* in area *A*, we computed the paired condition difference *d*_*i*_ = *C*1_*i*_ − *C*2_*i*_, where *C*1_*i*_ and *C*2_*i*_ denotes the average response of unit *i* in the specific contrast over a pre-specified analysis window. Because conditions were recorded from the same units, this paired difference removes any unit-level baseline.

The hierarchical model was:

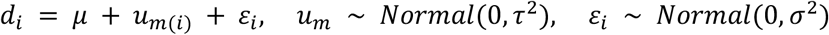

where *μ* is the across-mice fixed-effect mean condition difference (the parameter of interest), *u*_*m*(*i*)_ is a random intercept absorbing the systematic mouse-specific offset for the mouse from which unit i was recorded, and *ε*_*i*_ is the residual. The variance components *τ*^2^ (between-mouse) and *σ*^2^ (within-mouse, between-unit) were estimated by REML for the Wald inference.

The null hypothesis was H0: *μ* = 0. Because we had a directional prior expectation that GO evoked a greater population-level response than the control condition, all tests were one-sided (H1: *μ* > 0). We report p-value from a Wald test computed as 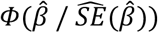 Fits in which the REML estimate of *τ*^2^ collapsed to the lower boundary 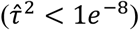 were flagged as unreliable and excluded from significance testing, because in that regime the Wald standard error degenerates to ordinary least squares on the unit-level data and substantially overstates significance.

To localize the period over which the condition difference was strongest, we repeated the nested model analysis above on a moving window. The per-unit response in each condition was the mean of the binned activity within a 25 ms window. The window was advanced in 1 ms steps across the analysis range (the period of visual stimulation, 0-500 ms following stimulus onset). For each window, we fit the model and recorded 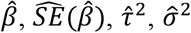, and the Wald p. Windows in which the model failed to converge or hit the variance-component boundary were excluded from significance summaries. We then determined which periods were consistently significant across windows >25 ms. We applied a clustering analysis in time. Because the test was one-sided (H1: μ > 0), we identified candidate clusters as runs of temporally contiguous windows whose statistic fell below a cluster-forming threshold of z = 1.645 (the one-sided 0.05 normal quantile).

In performing the analysis, we noted data from one animal was consistently anticorrelated (4/6 brain areas) with all other animals (mouse 12), so we excluded that animal from further analysis. After that exclusion, we identified three clusters, one in LM (283-344 ms, Wald p = 0.0398, n = 12 mice, n = 1359 units), one in PM (365-484 ms, Wald p = 0.0146, n = 12 mice, n = 719 units), and one in AM (359-400 ms, Wald p = 0.0445, n = 12 mice, n = 626 units). These clusters are consistent with the clusters we identified in the cluster-based permutation test analysis (Fig. 3), indicating the population-level statistical framework we chose to use throughout the manuscript is reliable.

### Functional clustering to identify potential global oddball detector population

We were interested to know whether we missed any global oddball detector populations in our analysis as we specifically evaluated along neuroanatomical axes (e.g., brain area, layer, cell type). It is possible that a global oddball detector population does not appear along these specific groupings. To address this, we performed an unsupervised functional clustering analysis to determine whether a global oddball detector population could be identified in a data-driven manner (Supplementary Fig. 8). We used an approach identical to that used in previous work in predictive processing^97^. The specific methodological details are provided in earlier works^98,99^.

Briefly, for each of the 4,807 neurons in mice (pooled across all recorded areas of the visual cortical hierarchy), firing-rate profiles were taken over a −100 to 1,000 ms peri-stimulus window for each of the ten conditions of the global–local oddball protocol. Each neuron’s ten condition profiles were concatenated into a single feature vector and divided by its standard deviation, so that clustering was driven by the shape of the response across conditions rather than by absolute firing rate.

Pairwise dissimilarities between neurons were quantified as Euclidean distances and agglomerated using Ward’s linkage. The resulting dendrogram was cut into an initial overclustering of 25 clusters, following the strategy of splitting until further subdivisions no longer revealed qualitatively distinct response profiles. Each candidate cluster was then evaluated for internal consistency using its homogeneity, defined as the mean pairwise Pearson correlation among the response profiles of its member neurons; clusters with homogeneity below [0.25], or containing fewer than 30 neurons, were considered non-robust and discarded, with their neurons left unassigned. A relatively low homogeneity threshold of 0.25 (as compared to the previous works) was chosen to avoid missing a weakly correlated cluster that may serve as a global oddball detecting population. The surviving clusters were then plotted across conditions for qualitative evaluation.

While several strongly correlated clusters were identified, e.g., #18 and #22 which appear clustered as stimulus-selective, we did not find any cluster where the response to the GO in the main block was notably different from the comparison control conditions. This lends further support to the idea that global oddball detection is not a ubiquitous feature of cortical processing.

### Neural instantiation of predictive coding

Predictive coding theorizes an internal model of the environment’s statistics; these statistics are typically taken to include the mean expected stimulus (typically denoted mathematically as μ) and the precision of the predicted stimulus (typically denoted mathematically as Π). In other words, PC is both subtractive and multiplicative: when sensory signals do not match the prediction, subtraction results in a larger value, called the prediction error, which is weighted by precision (Fig. 1). This error signal has been proposed to be computed in superficial layers 2 and 3 of the sensory cortex, which propagate the signal in the feedforward direction up the cortical hierarchy^2,3,100^. As a second-order statistic, the precision Π is always a positive, nonzero quantity, ensuring that prediction errors are fed up with the cortical hierarchy, even if attenuated. As they ascend, prediction errors instigate updates to the internal model to improve predictions, and associated prediction update signals flow back down the hierarchy.

### On hierarchical predictive coding

PC has also been proposed to occur over a series of computational steps, each comparing predictions to prediction errors from a previous step^1^. This “hierarchical predictive coding model” has since been modified to fit the neocortex’s structural and functional architecture^1^ (Fig. 1a). These hierarchical models efficiently predict and explain the cortical responses to first-order pattern violations (“local oddballs”), such as a change from repeated stimulation (e.g., a stimulus sequence such as x-x-x-y)^13,27^. Such local oddball responses depend on a simplified predictive model that the current stimulus will continue to repeat. However, local oddball responses persist during states of deep anesthesia when feedback connections are functionally weak^23,24^. As such, they do not provide strong evidence for PC, which envisions rich and multi-faceted predictions feeding back and modulating early sensory processing^2,3,29,30^. It is an open question whether more complex prediction errors engage the sensory cortex^13,22^. More complex prediction error computations and higher-order predictions can be studied using “global oddballs.” These second-order pattern violations occur when a repeated presentation of first-order pattern violations is interrupted (e.g., x-x-x-y / x-x-x-y / x-x-x-y / x-x-x-x). Global oddballs are a repeated stimulus, so responses cannot be explained by recent stimulus history, i.e., release from adaptation. As of now, there is only indirect evidence from human and non-human primate studies using functional Magnetic Resonance Imaging (fMRI), magneto- and electroencephalography (M/EEG), and local field potential (LFP) for global oddball responses in the sensory cortex^13^.

### Experimental design notes and rationale

Two cohorts of mice, each habituated to a different stimulus sequence, participated (Extended Data Fig. 1). To counterbalance the stimulus set across the population of animals, one cohort was habituated to a sequence of 3 rightward oriented gratings followed by a leftward-oriented grating (x-x-x-y, we refer to each stimulus in order as P1, P2, P3, P4), the other cohort to 3 leftward then a rightward (y-y-y-x). Following a 500 ms baseline, each stimulus was presented for 500 ms (with a 500 ms interstimulus interval) for a total trial duration of 4500 ms. Mice were habituated to these sequences (n=8 mice for x-x-x-y and n=8 mice for y-y-y-x sequences) throughout five sessions totaling 2000 trials. Two macaques were habituated to these sequences with the additional requirement of central fixation of the eyes on the screen. Monkeys were habituated over three sessions for 2850 complete trials before recordings. Then, in the experimental session, mice/monkeys were exposed to the not-yet-experienced global oddball sequence with a lesser frequency (20%). After that main experimental block, mice/monkeys experienced a block of random orientation sequences (“Random” block) before the “Control” block, where animals viewed alternating x-x-x-x and y-y-y-y sequences, rendering the same stimuli used in global oddball trials predictable. These controls were included to evaluate oddball detection in conjunction with the responses in the GLO experimental block.

### Additional considerations

While our findings are inconsistent with the hypothesized ubiquitous elements of Predictive Processing, additional issues should be considered. For one, we have chosen to study pure global oddballs, deviations from predictions induced by the lack of a stimulus change. Other forms of prediction that depend on local stimulus change (e.g., a stimulus that is both a local and global oddball) have been found in the mouse visual cortex in other experiments^12,87,97^. However, more complex forms of prediction processing envisioned by PC models, such as the global oddball and omission responses^11,28,30^, do not appear to engage visual sensory cortical responses with stimulus-specific activity^101^, as we have found. Another important consideration is the degree of attention paid to the oddballs by the animals. Previous work suggested that global oddballs are not observed under certain states of unconsciousness^19,20^. We did not explicitly manipulate attention, but behavioral analyses suggested that animals recognized the global oddballs (Extended Data Figs. 2,3). Future studies should explicitly consider the role of gain modulation on prediction error responses by manipulating the attentional state.

---

**Supplementary Table 1.** Overview of existing predictive processing models.

| **Study Number** | **MODEL** | **Reference** | **Model Type** |
| --- | --- | --- | --- |
| 1 | Attinger et al. (2017) | Cell, 169(7), 1291-1302.e14 | Empirical |
| 2 | Bastos et al.(2012) | Neuron, 76(4), 695–711 | Theoretical |
| 3 | Bastos et al. (2020) | PNAS, 117(49), 31459-31469 | Empirical |
| 4 | Bekinschtein et al. (2009) | PNAS, 106(5):1672-7 | Empirical |
| 5 | Chao et al. (2018) | Neuron, 100(5), 1252-1266 | Empirical |
| 6 | Friston (2010) | Nature Rev. Neuro., 11(2), 127-138. | Theoretical |
| 7 | Furutachi et al. (2024) | Nature, 633(8029), 398-406. | Empirical |
| 8 | Hertäg and Sprekeler, 2020 | Elife, 9, e57541 | Computational |
| 9 | Jiang & Rao (2024) | PLOS Computational Biology, 20(2). | Computational |
| 10 | Keller et al. (2012) | Neuron 74(5):809-15 | Computational |
| 11 | Keller and Mrsic-Flogel (2018) | Neuron, 100(2), 424-435 | Theoretical |
| 12 | Kiebel & Friston (2008) | PLoS Comput Biol, 4(11): e1000209. | Computational |
| 13 | Lao-Rodriguez et al. (2023) | Science Advances, 9, eabq8657 | Empirical |
| 14 | Lee, Pennartz & Mejias (2025) | PLoS Comput Biol 21(9): e1013469 | Computational |
| 15 | Mikulasch et al. (2023) | Trends in Neurosciences, 46(1), 45-59. | Theoretical |
| 16 | Nejad et al. (2025) | Nature Comms, 16(6178) | Computational |
| 17 | Rao & Ballard (1999) | Nature Neuroscience, 2(1):79-87 | Computational |
| 18 | Rao (2024) | Nature Neuroscience, 27(7), 1221-1235. | Computational |
| 19 | Spratling (2008) | Vision Research, 48(12), 1391-1408 | Computational |
| 20 | Spratling (2010) | J. Neurosci., 30(9),3531–3543 | Computational |
| 21 | Srinivasan et al. (1982) | Proc R Soc Lond B Biol Sci, 216(1205):427-59 | Empirical |
| 22 | Van Derveer et al. (2021) | Schizophrenia Bulletin, 47(5), 1385-1398. | Empirical |
| 23 | Wacongne et al. (2011) | PNAS, 108(51), 20754–20759; | Empirical |
| 24 | Wacongne et al. (2012) | J. Neurosci., 32(11), 3665–3678. | Computational |
| 25 | Westerberg, Xiong, et al. | *Current Study* | Empirical |

**Supplementary Table 2.** Mouse distribution of local-vs. global-oddball-stimulus selective units across data subsets. All data stratifications across mice (n=14) were evaluated to determine whether there were significant differences in the number of units that were selective for either the global or local oddball stimulus. Percentage of units from median split showing any preference for the GO vs. LO stimulus shown in the GO and LO stim preference columns. Cohen’s d was computed on the selectivity distributions across the populations of units (number of single units, N, indicated in 4^th^ column from the left) with effect size interpretation using standard thresholds (i.e., <0.2=negligible, 0.2-0.4=small, etc.). Most stratifications showed no disproportionality in the number of units that were selective for one stimulus over the other. Those that were disproportionate had only a small effect (highlighted in yellow) and were restricted to specific inhibitory interneuron subpopulations. Those were corrected for in relevant analyses (see Extended Data Fig. 10) via stratification.

| Area | Depth | Cell type | *N* | GO stim pref | LO stim pref | Cohen's *d* | Effect size | *N* * | Cohen's *d* * | Effect size* |
| --- | --- | --- | --- | --- | --- | --- | --- | --- | --- | --- |
| Any | Any | Any | 4870 | 49.75 | 50.10 | 0.013 | Negligible | — | — | — |
| Any | Any | PV | 256 | 49.22 | 50.78 | 0.026 | Negligible | — | — | — |
| Any | Any | SST | 210 | 46.19 | 53.81 | 0.077 | Negligible | — | — | — |
| Any | Deep | Any | 3005 | 49.58 | 50.32 | 0.016 | Negligible | — | — | — |
| Any | Deep | PV | 162 | 47.53 | 52.47 | 0.009 | Negligible | — | — | — |
| Any | Deep | SST | 111 | 43.24 | 56.76 | 0.126 | Negligible | — | — | — |
| Any | Middle | Any | 878 | 50.91 | 49.09 | 0.034 | Negligible | — | — | — |
| Any | Middle | PV | 36 | 47.22 | 52.78 | 0.078 | Negligible | — | — | — |
| Any | Middle | SST | 36 | 58.33 | 41.67 | 0.140 | Negligible | — | — | — |
| Any | Upper | Any | 766 | 49.87 | 49.61 | 0.008 | Negligible | — | — | — |
| Any | Upper | L2/3 Pyr | 559 | 50.27 | 49.02 | 0.004 | Negligible | — | — | — |
| Any | Upper | PV | 56 | 53.57 | 46.43 | 0.061 | Negligible | — | — | — |
| Any | Upper | SST | 40 | 42.50 | 57.50 | 0.254 | Small | 38 | 0.194 | Negligible |
| LM+PM+AM | Deep | PV | 88 | 44.32 | 55.68 | 0.068 | Negligible | — | — | — |
| LM+PM+AM | Deep | SST | 64 | 40.62 | 59.38 | 0.122 | Negligible | — | — | — |
| LM+PM+AM | Middle | PV | 16 | 50.00 | 50.00 | 0.249 | Small | 15 | 0.172 | Negligible |
| LM+PM+AM | Middle | SST | 20 | 60.00 | 40.00 | 0.214 | Small | 19 | 0.154 | Negligible |
| LM+PM+AM | Upper | PV | 32 | 40.62 | 59.38 | 0.163 | Negligible | — | — | — |
| LM+PM+AM | Upper | SST | 31 | 38.71 | 61.29 | 0.344 | Small | 27 | 0.180 | Negligible |
| LM+PM+AM | Extragranular | Any | 2208 | 47.83 | 52.04 | 0.021 | Negligible | — | — | — |
| V1 | Any | Any | 816 | 50.98 | 48.77 | 0.048 | Negligible | — | — | — |
| V1 | Any | PV | 48 | 54.17 | 45.83 | 0.098 | Negligible | — | — | — |
| V1 | Any | SST | 27 | 62.96 | 37.04 | 0.219 | Small | 26 | 0.173 | Negligible |
| V1 | Deep | Any | 553 | 49.01 | 50.99 | 0.016 | Negligible | — | — | — |
| V1 | Middle | Any | 150 | 56.67 | 43.33 | 0.171 | Negligible | — | — | — |
| V1 | Upper | Any | 113 | 53.10 | 45.13 | 0.044 | Negligible | — | — | — |
| V1 | Upper | L2/3 Pyr | 93 | 50.54 | 47.31 | 0.014 | Negligible | — | — | — |
| LM | Any | Any | 1429 | 48.15 | 51.85 | 0.030 | Negligible | — | — | — |
| LM | Any | PV | 42 | 47.62 | 52.38 | 0.054 | Negligible | — | — | — |
| LM | Any | SST | 86 | 44.19 | 55.81 | 0.028 | Negligible | — | — | — |
| LM | Deep | Any | 674 | 47.33 | 52.67 | 0.041 | Negligible | — | — | — |
| LM | Middle | Any | 260 | 51.15 | 48.85 | 0.044 | Negligible | — | — | — |
| LM | Upper | Any | 276 | 48.19 | 51.81 | 0.066 | Negligible | — | — | — |
| LM | Upper | L2/3 Pyr | 218 | 50.00 | 50.00 | 0.058 | Negligible | — | — | — |
| RL | Any | Any | 621 | 53.14 | 46.86 | 0.049 | Negligible | — | — | — |
| RL | Any | PV | 40 | 52.50 | 47.50 | 0.121 | Negligible | — | — | — |
| RL | Any | SST | 21 | 47.62 | 52.38 | 0.241 | Small | 20 | 0.184 | Negligible |
| RL | Deep | Any | 379 | 53.30 | 46.70 | 0.039 | Negligible | — | — | — |
| RL | Middle | Any | 152 | 51.32 | 48.68 | 0.058 | Negligible | — | — | — |
| RL | Upper | Any | 90 | 55.56 | 44.44 | 0.080 | Negligible | — | — | — |
| AL | Any | Any | 541 | 53.60 | 46.03 | 0.099 | Negligible | — | — | — |
| AL | Any | PV | 30 | 56.67 | 43.33 | 0.139 | Negligible | — | — | — |
| AL | Any | SST | 24 | 37.50 | 62.50 | 0.295 | Small | 22 | 0.192 | Negligible |
| AL | Deep | Any | 371 | 54.18 | 45.28 | 0.121 | Negligible | — | — | — |
| AL | Middle | Any | 113 | 50.44 | 49.56 | 0.017 | Negligible | — | — | — |
| AL | Upper | Any | 57 | 56.14 | 43.86 | 0.102 | Negligible | — | — | — |
| PM | Any | Any | 767 | 47.33 | 52.54 | 0.021 | Negligible | — | — | — |
| PM | Any | PV | 55 | 43.64 | 56.36 | 0.043 | Negligible | — | — | — |
| PM | Any | SST | 22 | 45.45 | 54.55 | 0.117 | Negligible | — | — | — |
| PM | Deep | Any | 570 | 46.49 | 53.51 | 0.027 | Negligible | — | — | — |
| PM | Middle | Any | 98 | 51.02 | 48.98 | 0.015 | Negligible | — | — | — |
| PM | Upper | Any | 99 | 48.48 | 50.51 | 0.009 | Negligible | — | — | — |
| PM | Upper | L2/3 Pyr | 68 | 50.00 | 48.53 | 0.058 | Negligible | — | — | — |
| AM | Any | Any | 696 | 48.28 | 51.44 | 0.001 | Negligible | — | — | — |
| AM | Any | PV | 41 | 43.90 | 56.10 | 0.131 | Negligible | — | — | — |
| AM | Any | SST | 30 | 43.33 | 56.67 | 0.198 | Negligible | — | — | — |
| AM | Deep | Any | 458 | 50.66 | 49.13 | 0.049 | Negligible | — | — | — |
| AM | Middle | Any | 105 | 41.90 | 58.10 | 0.161 | Negligible | — | — | — |
| AM | Upper | Any | 131 | 45.04 | 54.20 | 0.054 | Negligible | — | — | — |

**Supplementary Table 3.** Open-weight LLMs used in this work.

| Model Name (handle-mlx) | Size | Release Date and info |
| --- | --- | --- |
| gemma-3-27b-it | 27B | Google DeepMind, 2025-03 |
| gemma-4-31b-it | 31B | Google DeepMind, 2026-04 |
| deepseek-r1-distill-llama-70b | 70B | DeepSeek-AI / Meta AI (Llama Base), 2025-01 |
| gpt-oss-claude-4.5-sonnet | 20B | Community Distillation (Teacher: Claude 4.5), 2026-03 |
| mistral-nemo-12b-thinking | 12B | Mistral AI & NVIDIA, 2024-07 |
| olmo-3-32b-think | 32B | Allen AI (Ai2), 2025-12 |
| phi-4-reasoning-plus | 14B | Microsoft, 2025-03 |
| qwen3-14b-gemini-3-pro | 14B | Qwen (Community Fine-tune), 2025-12 |
| qwen3-14b-claude-4.5-sonnet | 14B | Qwen (Community Fine-tune), 2025-12 |
| qwen3.5-40b-claude-4.5-opus | 40B | Qwen (Community Fine-tune), 2026-01 |

**Supplementary Table 4.** Technical Glossary for Local vs. Global Oddball context. S={.} represents the sensory input, E[S] represents the expected sensory input. When S={A} and E[S] = {A}; the prediction error is zero (match). When S={B} and E[S] = {A}; the prediction error is non-zero (mismatch). LO and GO differ by the temporal scale of expectation.

| Term/Context | Description | Definition (notation) |
| --- | --- | --- |
| Oddball | A surprising event, often a mismatch | $S = \{B\}$ given $E[S] = \{A\}$ |
| Local Oddball (LO) | An oddball that is immediate or short-term, often confounded by low-level adaptation mechanisms. | $S = \{AB\}$ given $E[S] = \{AA\}$ |
| Global Oddball (GO) | An oddball that is not LO, often contextual, that is not caused by adaptation mechanisms. | $S = \{AA\}$ given $E[S] = \{AB\}$ |

**Supplementary Table 5.** Technical Glossary for Hypothesis 1 (Predictive Suppression). LO = local oddball. GO = global oddball.

| Factor Number | Factor Name | Definition | Context Scope |
| --- | --- | --- | --- |
| 1 | Subtractive Inhibition (SST) | Linear input suppression via somatic inhibition (Somatostatin-mediated). | LO+GO |
| 2 | Divisive Inhibition (PV) | Multiplicative gain reduction (Parvalbumin-mediated). | LO+GO |
| 3 | Inhibition (GABA) | Chloride-mediated inhibition for prediction suppression. | LO+GO |
| 4 | Habituation to Sequence | Habituation suppresses local oddball response relative to a novel local oddball. | LO |
| 5 | Synaptic Depression (Adaptation) | Passive fatigue of synaptic efficacy due to repetition of stimulus. | LO |
| 6 | Activity Suppression | Reduction in firing rates for expected (predictable) stimuli. | LO+GO |
| 7 | Selective Sharpening | Signal-to-noise increase for predictable stimuli, where noise is selectively suppressed to highlight relevant information. | LO+GO |
| 8 | Alpha/Beta Mediated Suppression | Low-frequency bands (8-30Hz) associated with predictions, inhibiting prediction error signals. | LO+GO |
| 9 | VIP-Mediated Disinhibition | VIP-mediated inhibition of SST/PV neurons, leading to disinhibited pyramidal activity during prediction error. | LO+GO |
| 10 | Precision Weighting (Gain) | Top-down amplification of error units, leading to disinhibited gain in response to attended or highly precise stimuli. | LO+GO |
| 11 | E/I Balance Shift | Dynamic adjustment toward higher inhibition for predictable stimuli, leading to suppressed neural firing. | LO+GO |
| 12 | Omission Response | A generated internal signal due to absence of expected input | LO+GO |

**Supplementary Table 6.** Technical Glossary for Hypothesis 2 (Feedforward Error Propagation).

| Factor Number | Factor Name | Definition | Context Scope |
| --- | --- | --- | --- |
| 13 | Feedforward Deviance Detection | Detection of mismatch signals that explicitly propagate in the feedforward (ascending) direction to higher cortical areas. | LO+GO |
| 14 | Feedforward AMPA | Fast excitatory drive mediated by AMPA receptors, specifically conveying prediction-error signals in the feedforward direction. | LO+GO |
| 15 | Feedforward NMDA | Voltage-dependent amplification (bursting) of error signals, facilitating their robust feedforward propagation through the hierarchy. | LO+GO |
| 16 | Feedforward Ascending Gamma | Rhythmic synchronization (30-90Hz) that temporally packages prediction-error signals for efficient feedforward transmission. | LO+GO |
| 17 | Absence of Feedback Error | The functional requirement that the prediction-error signal be feedforward-directed, distinguishing it from local or feedback signals. | LO+GO |
| 18 | Feedforward Non-local Supragranular Activity (L2/3) | Error-signaling neurons in Layers 2/3 projecting into the next area, demonstrating feedforward signal propagation. | LO+GO |
| 19 | Feedforward Non-local Granular Activity (L4) | The arrival of prediction error signals in Layer 4, non-local representing the canonical entry point for feedforward propagation into the cortical column. | LO+GO |
| 20 | Feedforward Non-local Directed Connectivity | Directional measures (Granger Causality/Transfer Entropy) confirming the directed feedforward propagation of prediction error from lower to higher areas. | LO+GO |
| 21 | Feedforward Non-local Activation | Functional activation patterns specifically tracing the feedforward propagation path through the hierarchy. | LO+GO |
| 22 | Ascending Latency Shift | Systematic increases in response latency that characterize the feedforward propagation of prediction error through successive levels of the hierarchy. | LO+GO |
| 23 | Feedforward Error Propagation | Explicit experimental evidence that the prediction error signal is transmitted primarily via feedforward propagation pathways. | LO+GO |
| 24 | Subcortical Feedforward Relaying | Relaying or generation of prediction error signals by subcortical structures (Thalamus/BG), initiating or sustaining their feedforward propagation to the cortex. | LO+GO |

**Supplementary Table 7.** Technical Glossary for Hypothesis 3 (Ubiquity).

| Factor Number | Factor Name | Definition | Context Scope |
| --- | --- | --- | --- |
| 25 | Canonical Microcircuit Ubiquity | The ubiquitous presence of the L2/3 Error and L5/6 Prediction motif repeating across most or all levels of the cortical hierarchy. | LO+GO |
| 26 | Hierarchical Mechanism Invariance | The ubiquitous nature of the predictive coding mechanism, functioning similarly in the hierarchy from V1 to PFC. | LO+GO |
| 27 | Hierarchical Activity Ubiquity | The ubiquitous presence of prediction-error activity, detectable across all or most levels of the system's brain hierarchy. | LO+GO |
| 28 | Hierarchical CSD Ubiquity | The ubiquity of current source density profiles, which show consistent laminar patterns across most or all hierarchical levels. | LO+GO |
| 29 | Cross-Scale Hierarchical Ubiquity | The ubiquity of effects observable across scales (single units to LFP) throughout most levels of the cortical hierarchy. | LO+GO |
| 30 | Hierarchical Presence (V1-PFC) | The ubiquity of the mechanism across all levels of the hierarchy, such as low-level (V1), mid-level (V4), and high-level (PFC) areas. | LO+GO |
| 31 | Cross-Modal Ubiquity | Presence of the mechanism across multiple sensory modalities (visual, auditory, somatosensory) throughout the hierarchy. | LO+GO |
| 32 | Interspecies Hierarchical Ubiquity | Conservation of the hierarchical predictive coding mechanism across different species (e.g., mouse vs. primate). | LO+GO |
| 33 | Temporal Stability of Ubiquity | The consistent and non-transient presence of these mechanisms across the hierarchy over long recording sessions. | LO+GO |
| 34 | Hierarchical Order Stability | Evidence that the predictive mechanism is ubiquitous and not localized to a single hierarchical pole. | LO+GO |
| 35 | Population-Wide Ubiquity | Consistency of the mechanism across diverse neural populations and cell types throughout the hierarchy. | LO+GO |
| 36 | State-Independent Ubiquity | Presence of the mechanism across different brain states (e.g., wakefulness, sleep, anesthesia) throughout the hierarchy. | LO+GO |

**Supplementary Fig. 1.**
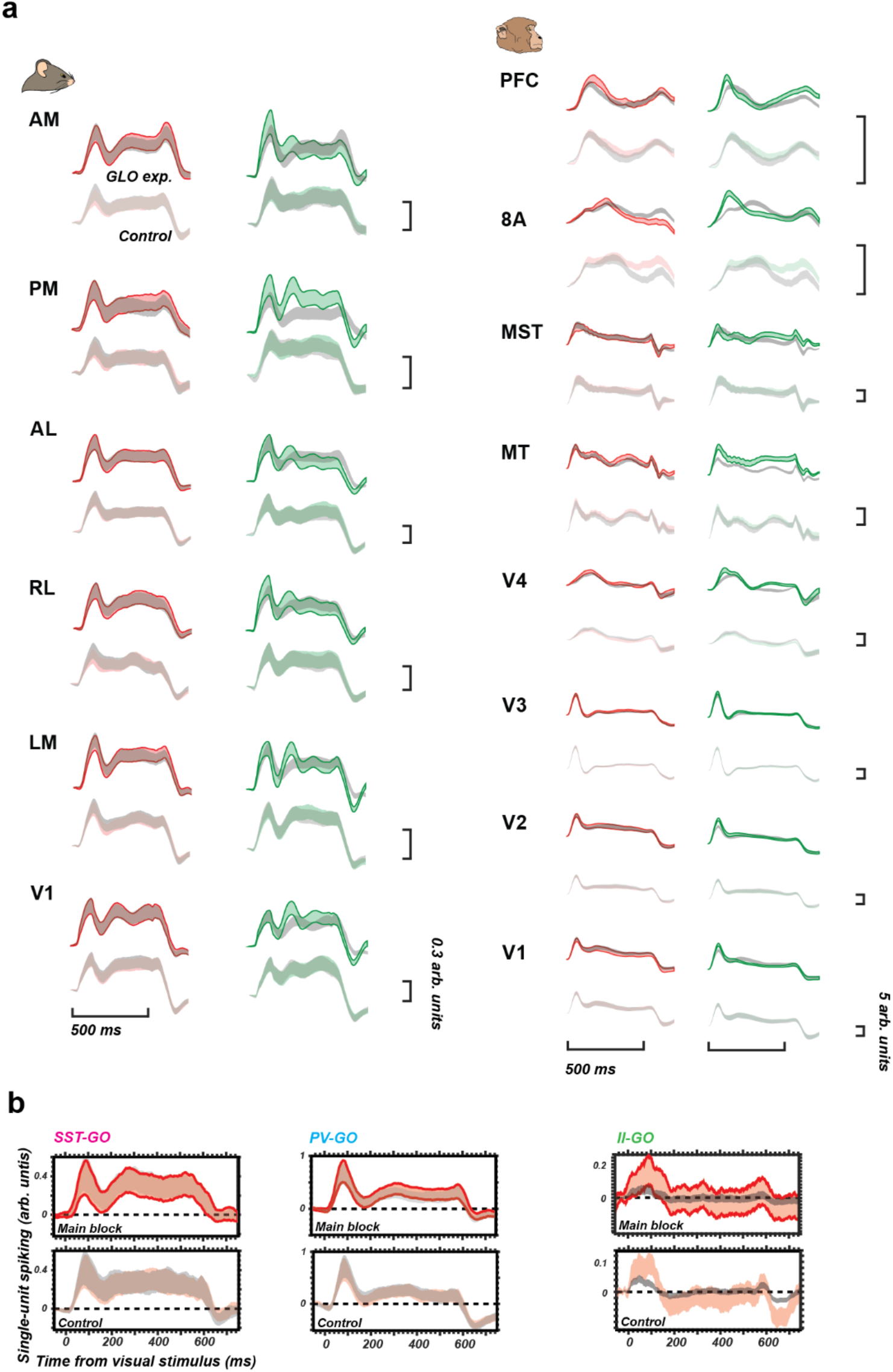
Component traces of global and local oddball contrast. **a**, For each area, red and green outlined traces (top) show 95% confidence interval of P4 of global oddball (left, red) and local oddball (right, green), and overlapping grey traces show 95% confidence interval of P3 of the same sequence. Red and green non-outlined (bottom) show 95% confidence interval of P4 of control sequence, and overlapping grey traces show 95% confidence interval of P3 of the same sequence. The green traces shown the corresponding local oddball responses. All visually responsive units from Fig. 2 and Fig. 3 included: mouse n=14, V1: n=816, LM: n=1429, RL: n=621, AL: n=541, PM: n=767, AM: n=699; monkey n=2, V1: n=136, V2: n=179, V3: n=402, V4: n=135, MT: n=70, MST: n=49, 8A: n=201, PFC: n=419. **b**, single unit normalized spiking responses in optotagged inhibitory interneurons (left panel, SST cells in n=7 mice: n=210 units; middle panel, PV cells in n=7 mice: n=256 units; right panel, inhibitory interneurons in n=1 monkey n=17 units), Bands are 95% confidence intervals across units. Top, main block spiking response with red P4 and gray P3. Bottom, control block spiking response with red P4 and gray P3.

**Supplementary Fig. 2.**
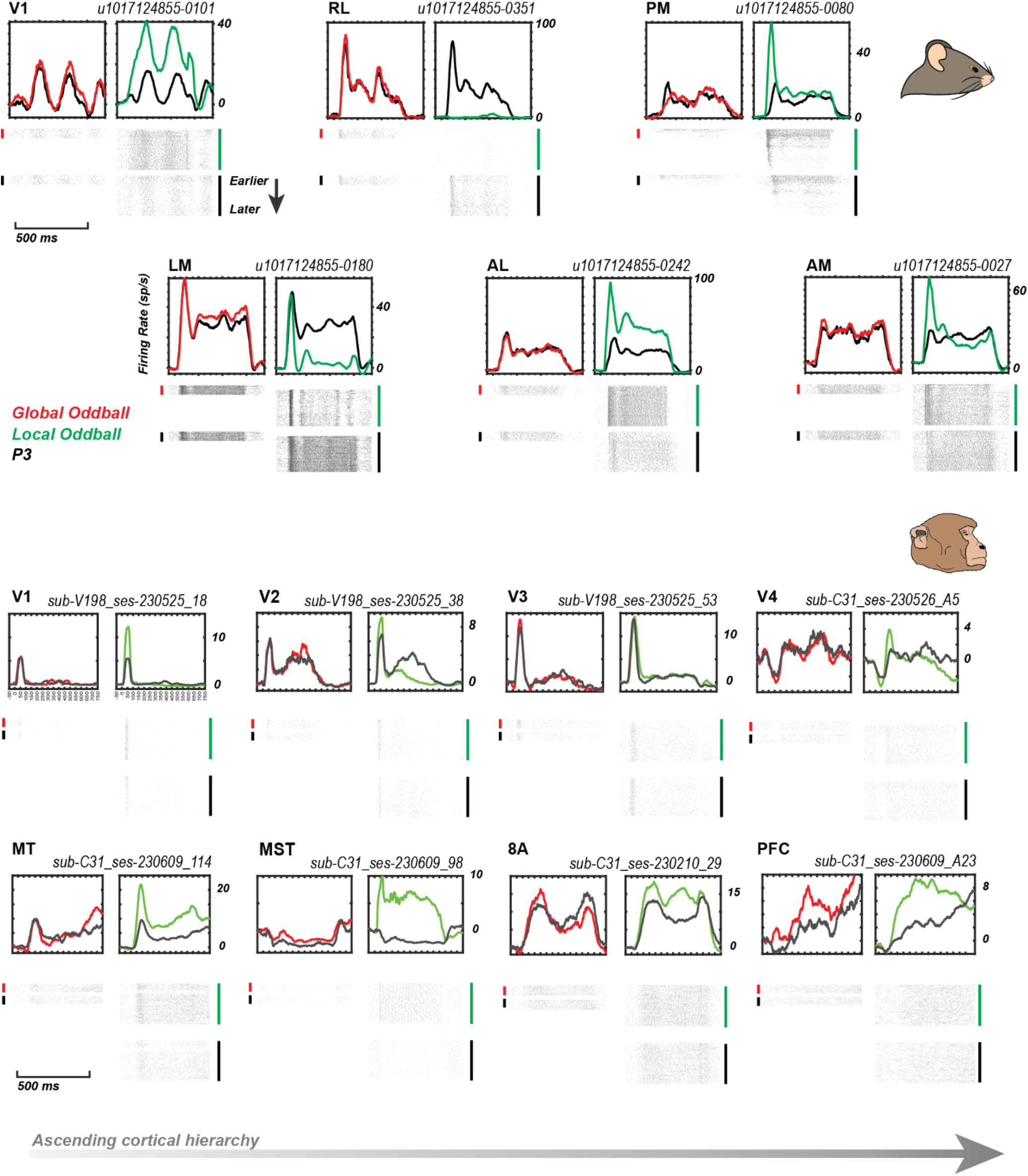
Examples single-unit responses in mice and monkeys. For each area in the mouse and monkey, one example single-unit is displayed. For the mouse areas, all single-units come from one mouse (1017124855). For the monkey, units are taken from two monkeys (V198, C31). The top left subpanel of each unit displays the convolved spike train for the P4 global oddball response (red) and the local oddball response (green) on the right. The respective P3 responses (black) immediately prior to each oddball type are displayed on the same axes. Below the convolutions are the raster plots for each condition. Top left, global oddball; bottom left, P3 on global oddball sequences; top right, local oddball; bottom right, P3 on local oddball sequences. Topmost line in each raster is the first trial of each condition with the subsequent lines being later in the session. There are fewer rasters for the left side plots as global oddball occurred in only 1 in 5 trials.

**Supplementary Fig. 3.**
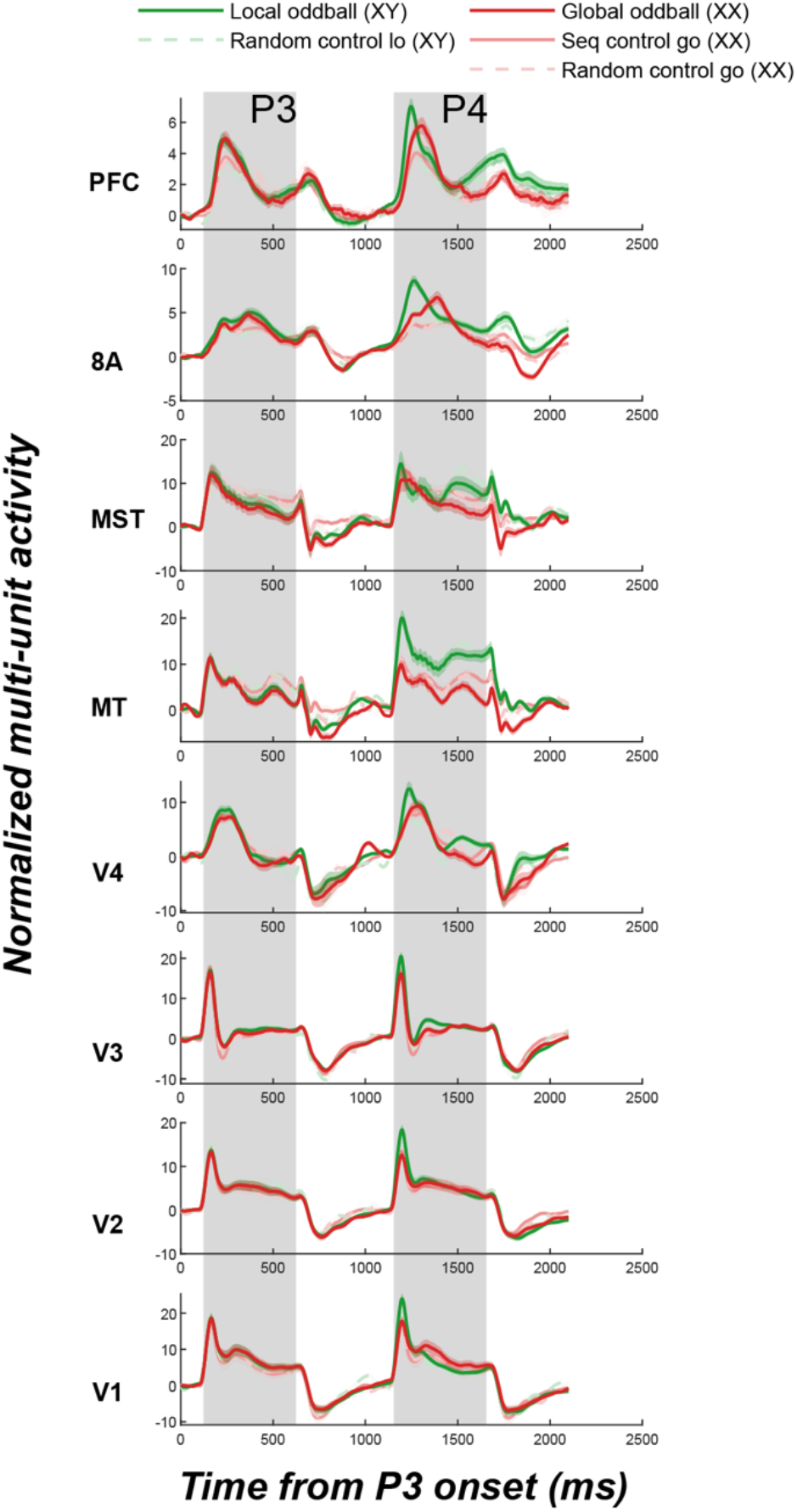
Average response to Presentation 3 (P3) and Presentation 4 (P4) in various experimental contexts of the session. Third stimulus (P3) and fourth stimulus (P4) across brains areas from different trial types.

**Supplementary Fig. 4.**
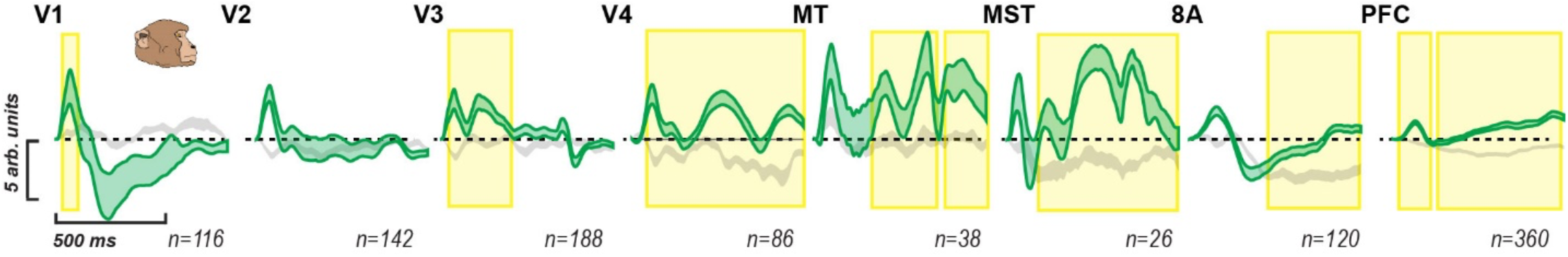
Local oddball data in mice and monkeys, stratified to equalize unit selectivity. The population of multi-units were stratified to have a balanced selectivity for ‘x’ vs. ‘y’ responses across the population of multi-units per area (i.e. median not significantly shifted away from zero for response to ‘x’ vs. ‘y’ when presented in the randomization control block). Bands are 95% confidence intervals across units in an area. During stratification, units with extreme values in selectivity were systematically removed until the selectivity distribution becomes balanced. Green bands are the P4-P3 local oddball in the main block; gray bands are the P4-P3 local oddball in the control block (same local oddball contrast as in Fig. 2 – except performed on the stratified population). Yellow highlights reflect significant population local oddball detection periods, P<0.05, corrected using nonparametric, cluster-based permutation tests.

**Supplementary Fig. 5.**
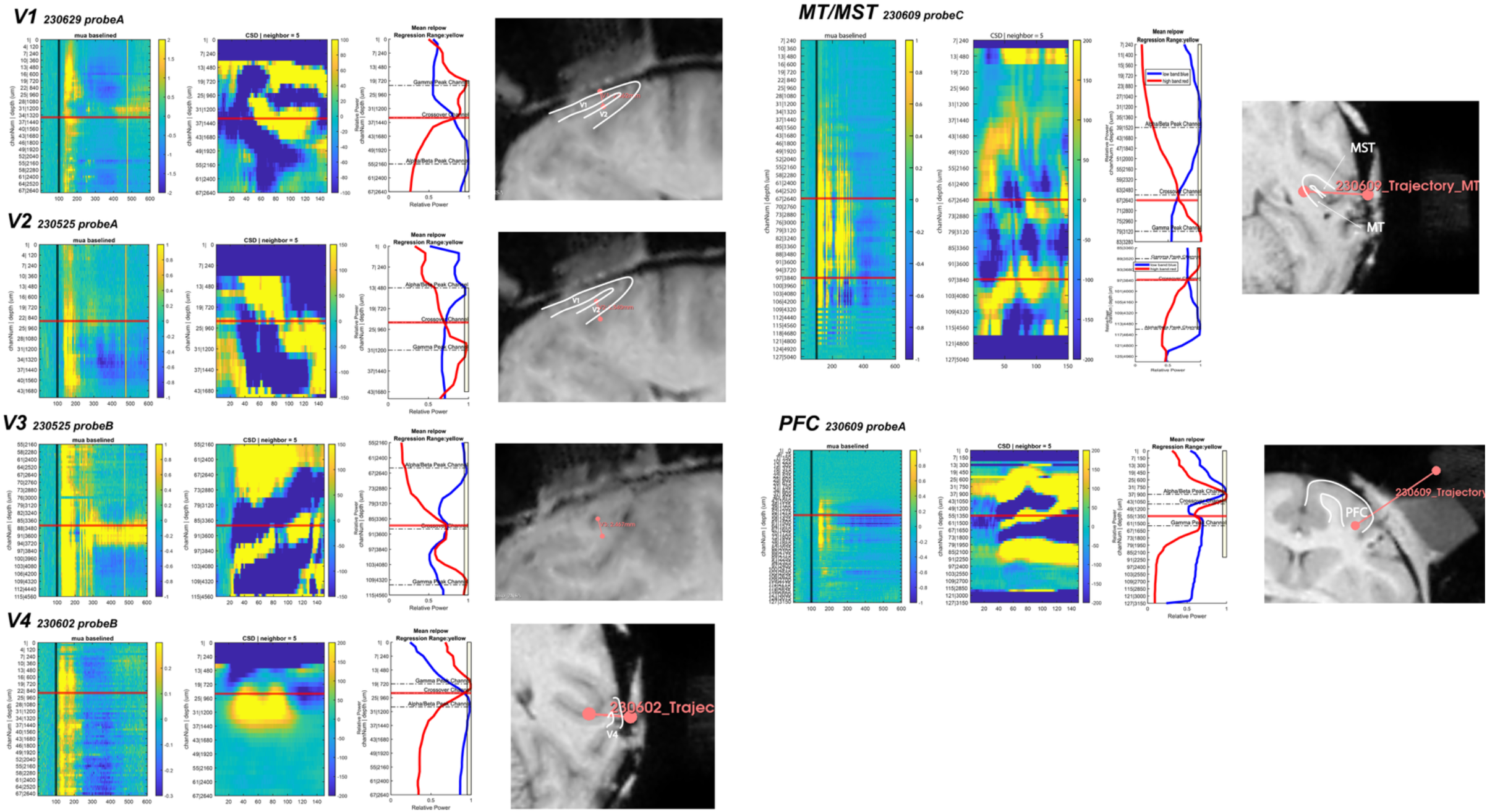
Spectrolaminar profile of monkey electrophysiology recordings included in layer-specific analyses. Panels for each area, from left to right, show corresponding evidence for layer 4 identification using the earliest onset of MUA channel, earliest sink in current source density, beta/gamma relative power crossing, and MRI anatomical reconstruction of the corresponding electrode trajectory.

**Supplementary Fig. 6.**
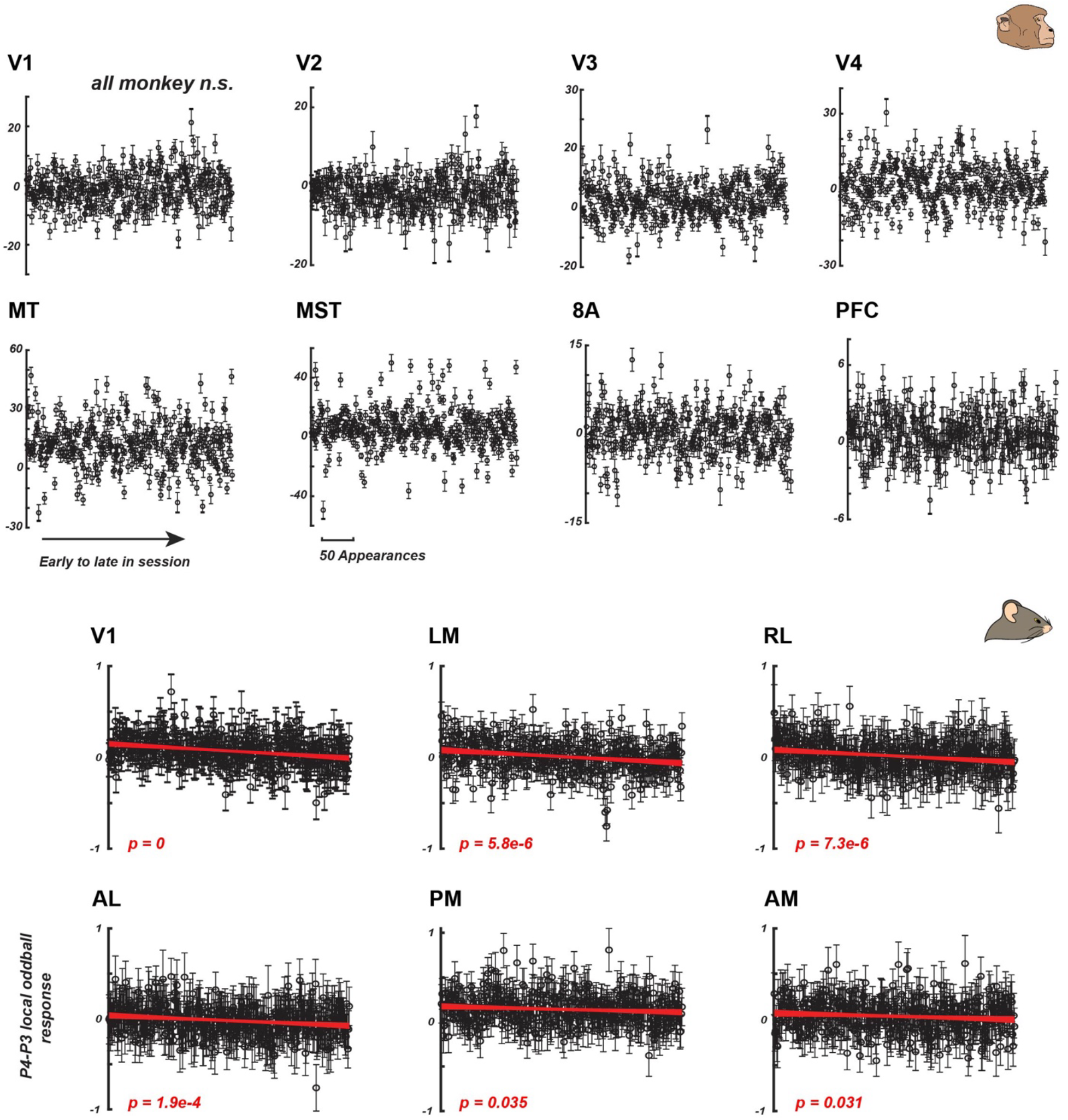
Local oddball responses over the main block are stable in monkeys but not in mice. Local oddball trials by position in the main block. The effect of local oddballs was examined by progression in the main block. The order of occurrence of local oddball trials is plotted on the x-axis. The neural responses of P4-P3 are plotted on the y-axis (mean across channels +/-2SEM). Linear regressions between these variables were tested separately for each indicated area, and the resulting fits are plotted in red. No significant regression was found in monkeys.

**Supplementary Fig. 7.**
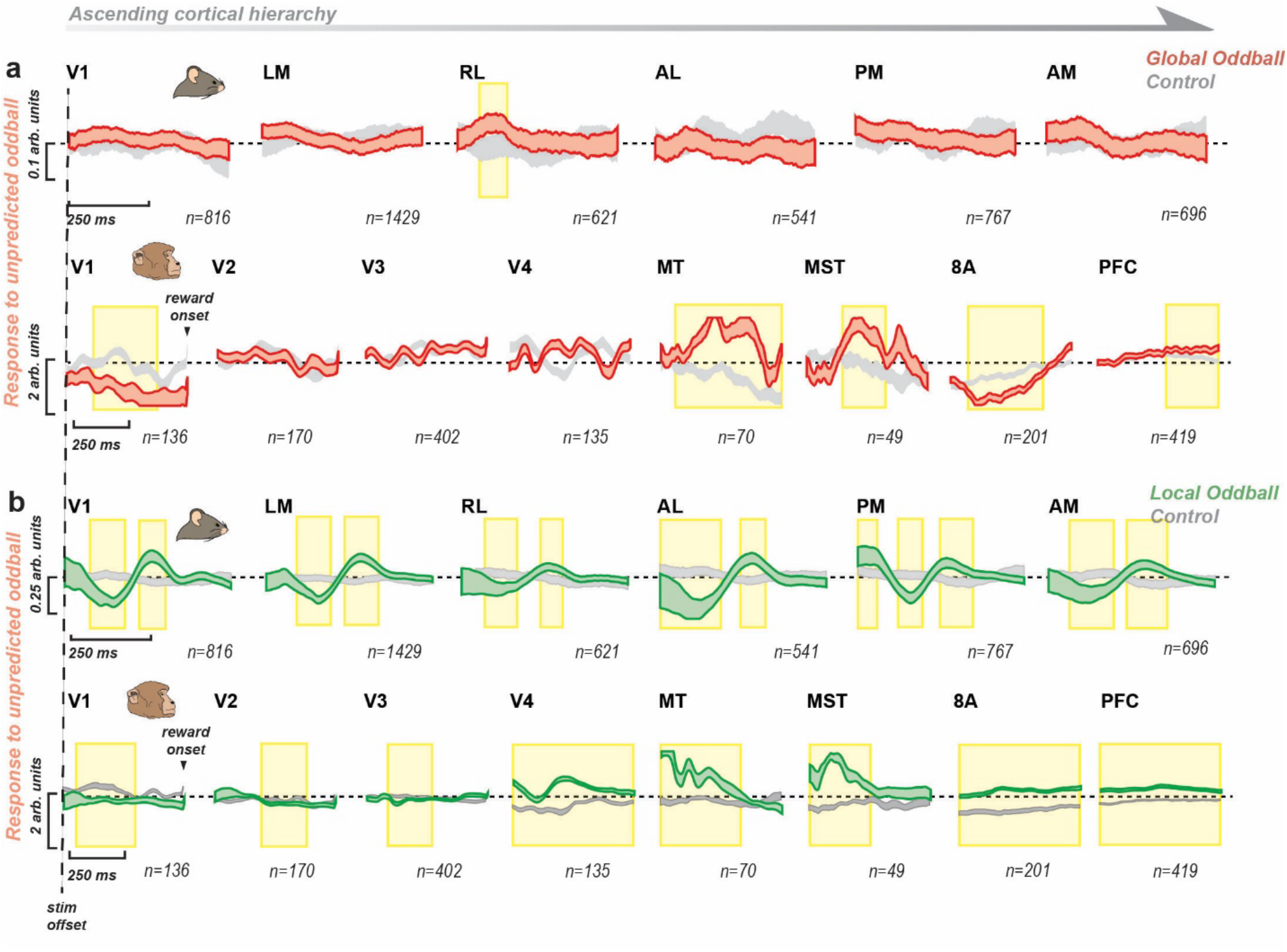
Global oddball responses during the off-response period. Off-period response to unpredictable global oddball and predictable local oddball. **a**, Red bands are the P4-P3 global oddball in the main block for 500ms after the offset of the stimulus; gray bands are the P4-P3 global control in the control block for 500ms after the offset of the stimulus. Shaded areas indicate 95% confidence interval. Yellow boxes indicate significant time periods during which the oddball is different from the respective control (FWE-corrected). The dotted line indicates stimulus offset. In monkeys, rewards occurred at the end of 500ms after stimulus offset (indicated by arrow, where data shows beginning of reward artifact) **b**, Green bands are the P4-P3 local oddball in the main block for 500ms after the offset of the stimulus; gray bands are the P4-P3 local control in the control block for 500ms after the offset of the stimulus. Shaded areas indicate 95% confidence interval.

**Supplementary Fig. 8.**
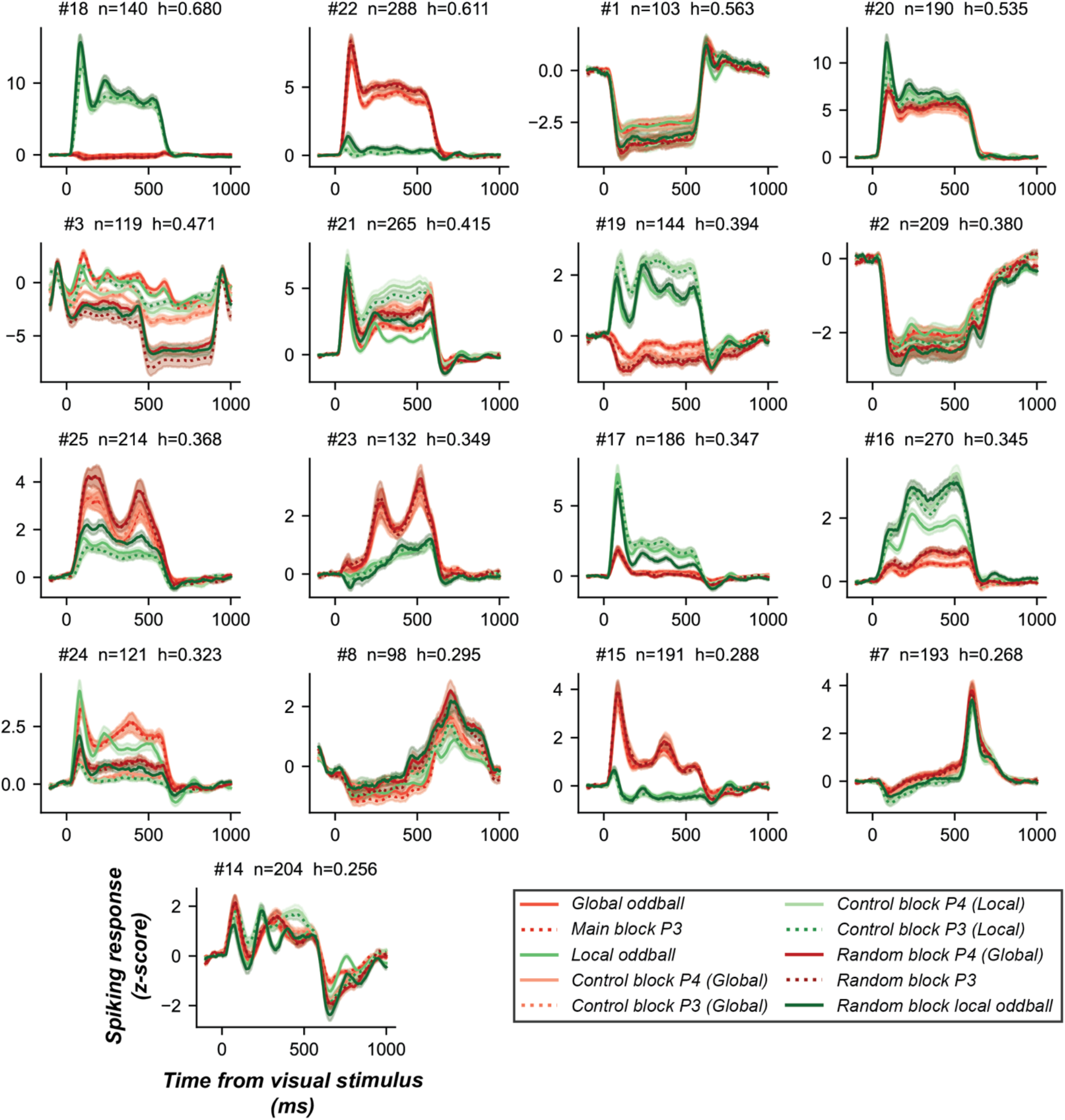
Data-driven functional clustering does not reveal a global oddball detector functional cell class. Functional clustering analysis (see *Supplementary Text*) was performed across 10 stimulus-response conditions, independent of the neuron’s recording location, to determine whether a data-driven approach reveals a global oddball-detecting functional class. Cross-validated analysis revealed 17 clusters that met the homogeneity criterion (h>0.25). Traces are the cluster-mean responses after z-score normalization. Red traces indicate conditions where the GO stimulus was presented, and green traces indicate conditions where the LO stimulus was presented. Solid traces indicate P4 presentations and dashed traces indicate P3 presentations. Lighter-to-darker traces indicate the specific block of the stimulus-response condition (lightest = control block, middle = main block, darkest = random block).

**Supplementary Fig. 9.**
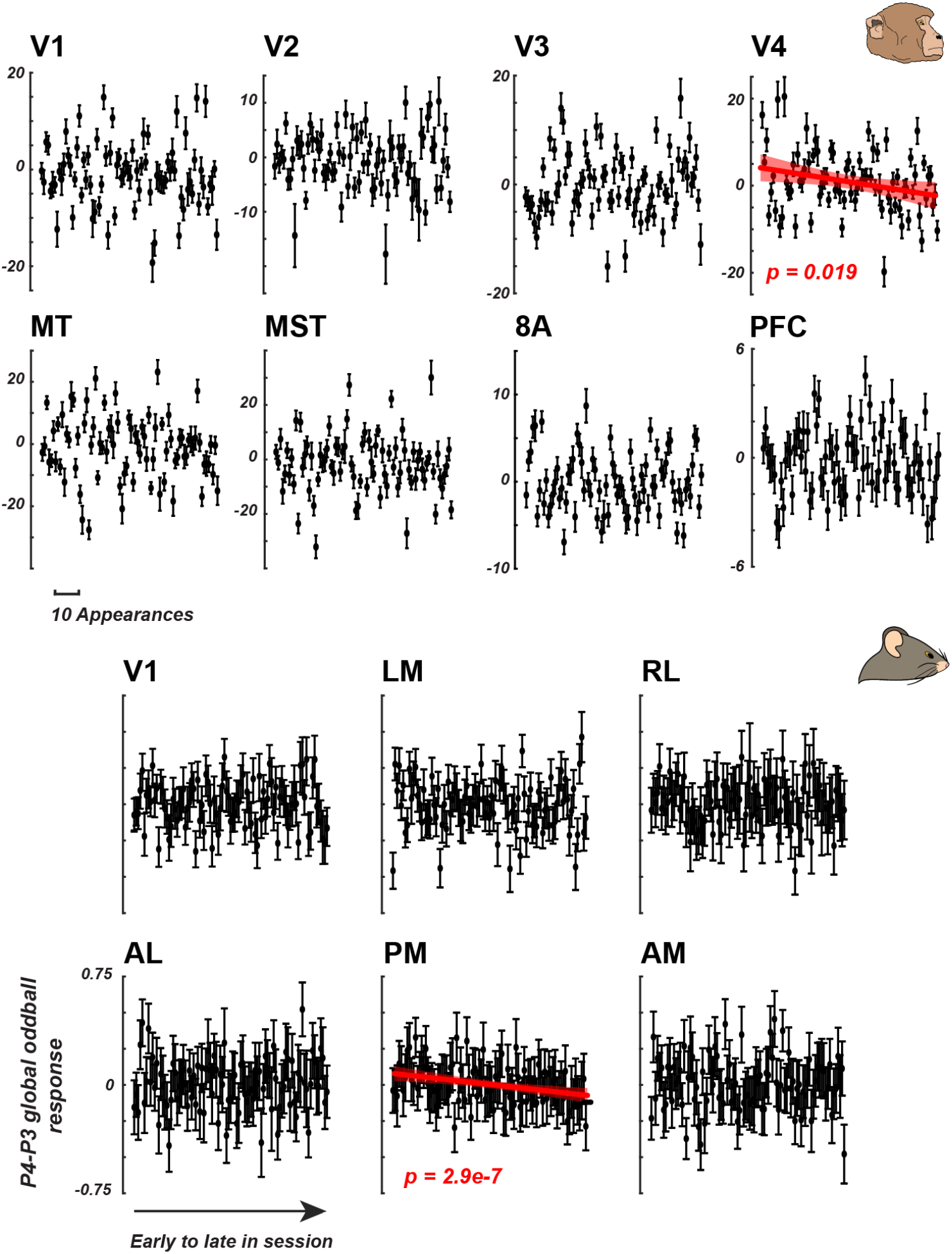
Global oddball trials by position in the main block. Effect of global oddballs were examined by progression in main block. The order of occurrence of global oddball trials are plotted on the x axis. The neural responses of P4-P3 are plotted on the y-axis (mean across channels +/-2 SEM). Linear regressions between these variables were tested separately for each indicated area and found to be significant in areas V4 in monkeys and PM in mice (P value indicated in graph). Only significant regression fits are plotted.

**Supplementary Fig. 10.**
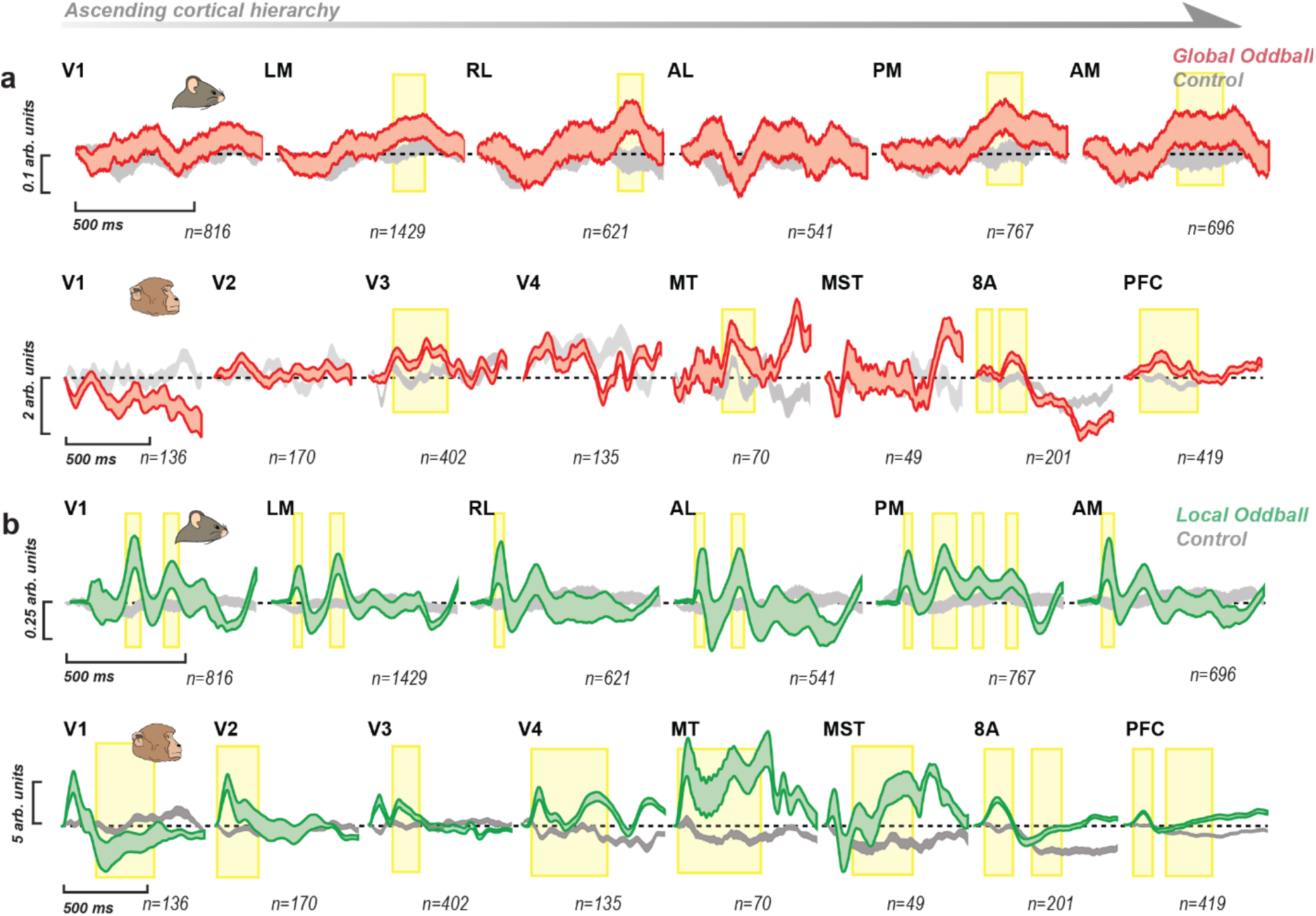
Late sequence control effects. Unpredictable global oddball and predictable local oddball responses when compared to control sequences after first 20 trials. (A) Red bands are the P4-P3 global oddball in the main block; gray bands are the P4-P3 global control in the control block, excluding the first 20 trials in the control block. (B) Green bands are the P4-P3 local oddball in the main block; gray bands are the P4-P3 local control in the control block, excluding the first 20 trials in the control block.

**Supplementary Fig. 11.**
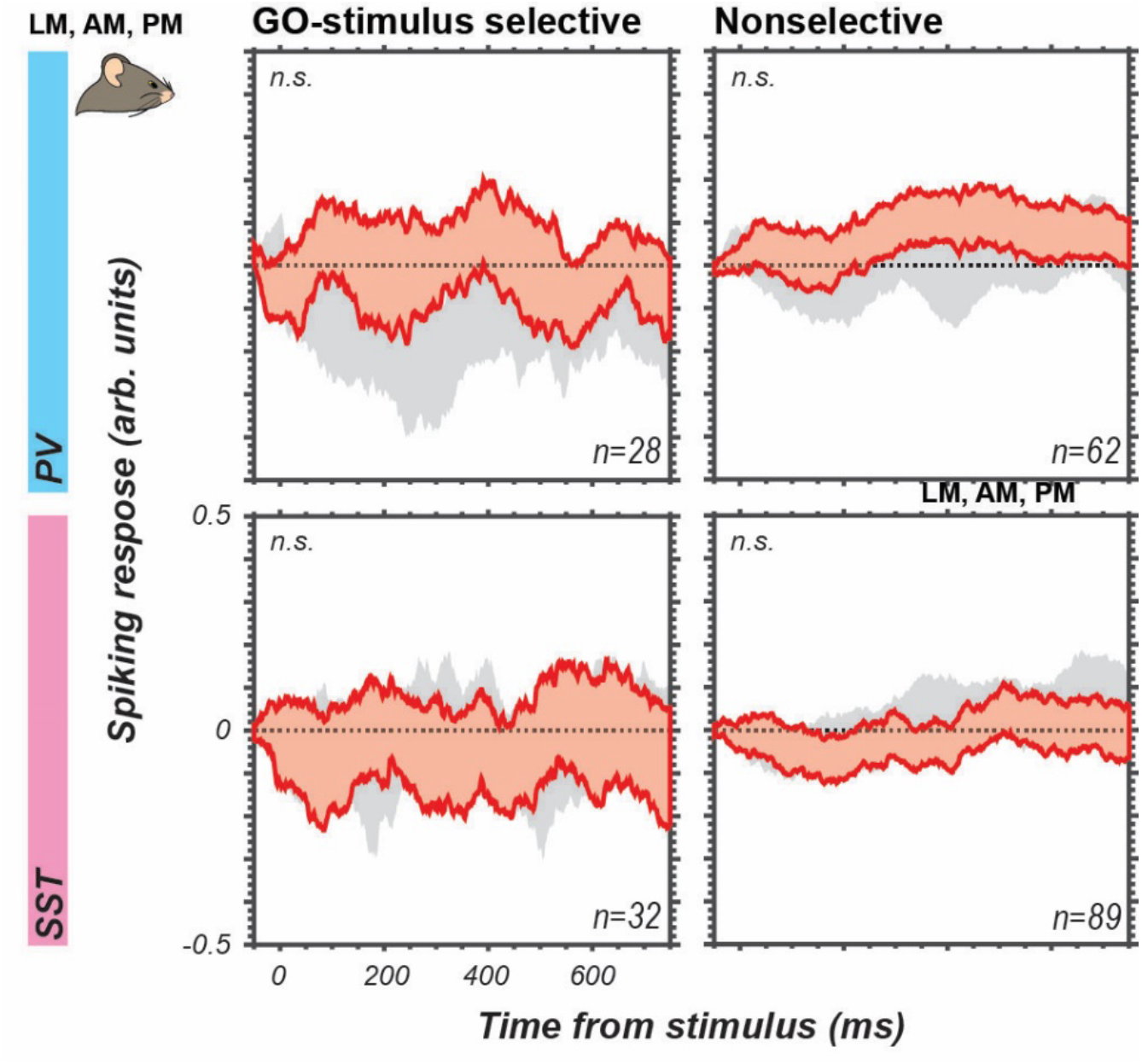
Global-oddball-stimulus-selective inhibitory interneurons are not global oddball detectors. GO contrast (P4-P3, GO sequence in main block) vs. control contrast (P4-P3; GO sequence in the control block), averaged across units for each inhibitory cell type (PV, top, cyan; SST, bottom, magenta) across the areas in all mice (n=14 mice), which were global oddball-detecting at the population level (LM, PM, AM). Data were further parsed to specifically evaluate units that were selective for the GO stimulus (see *Methods*) (left) vs. those that were found to be nonselective (right). Bands are 95% confidence intervals across units (number of units, n, indicated in the figure) in an area. No significant differences were found across any of the four data subsets using the cluster-based permutation test, indicated by n.s. in the upper lefthand corner of each subplot.

